# MSLASpheroidStamp: 3d cell spheroids for everyone

**DOI:** 10.1101/2024.05.12.593682

**Authors:** A. Minin, T. Semerikova, A.V. Belousova, O. Karavashkova, V. Pozdina, M. Tomilina, I. Zubarev

## Abstract

3D cell cultures, such as cell spheroids, are actively used in biology for modeling biological processes, studying intercellular interactions and pharmacological compounds screening and are becoming indispensable objects in cell culture laboratories. There are many methods for producing spheroids, varying in cost and convenience. One of the most handy and affordable is the use of agarose microwells. We have developed approaches to fabricate agarose microwells in standard culture plastic with the assistance of a hobby-grade MSLA 3D printer. The use of 3D printing allows you to customize microwells in a wide range of shapes and sizes and scale the production process from a few spheroids to tens of thousands. We have shown that it is possible to create gel microwells in a dish with a glass bottom, which allows us to easily realize time-lapse confocal microscopy of spheroids, and we have also performed in situ optical clearing in the same dishes to study the spheroid structure. We demonstrated the ability to study the cytotoxicity of various substances and nanoparticles in commonly used 96-well plates.

And finally, in this article we describe the difficulties and limitations of our approach and suggest ways for solving them, allowing the reader not only to reproduce it, but also to adapt it to the specific needs of a certain laboratory, using provided 3D models and instructions.

## Introduction

Recently, three-dimensional cell cultures like spheroids and organoids have gained equal importance in laboratory research compared to traditional suspension or adherent cell cultures. The behavior of cells in 3D culture is similar to their interactions and growth in the animal body, and data obtained from experiments on 3D cultures can be more valuable for drug development [1] and research in developmental [2] and cancer biology fields [3]. Despite the benefits, 3D culture demands increased effort and expenses in production. In this study, we suggest employing a do-it-yourself (DIY) method to lower costs and enable the affordable production of cell spheroids in any biological laboratory.

There are many ways to make cell spheroids: using nonadherent cultural plastic, hanging drop technique, agarose 3D microwell technique [4] or using specialized viscous media such as Matrigel [5].

Matrigel and other analogs are an almost mandatory matrix for culturing organoids, but its use is expensive and not always suitable for high-throughput drug screening. Nonadherent petri dishes do not provide control over the spheroid’s size, which negatively affects the reproducibility of the experiment [6]. Hanging drop technique is a rather labor-intensive and poorly reproducible procedure [7]. Agarose 3D microwell is a simple, accessible and reproducible technique that does not require large expenses, which is why we chose this method [8]. There are two approaches: the first is a plastic stamp which creates deepenings - microwells - in agarose poured into a container (e.g. a cell culture dish or a well of a culture plate) [9,10]; the second is a silicone mold with micropins that is filled with agarose [11,12]. We decided to try both approaches and study how and where they are applicable. At the same time, our task was not just to implement such a project, which has already been partially done, but also to make it accessible for reproduction by the any scientist, make it DIY (Do It Yourself).

In recent years, the term DIY (in science) often involves the use of a 3D printer as a device that allows the reliable and reproducible production of complex objects. A 3D printer in a modern scientific laboratory, including a biological one, is a relevant device that allows you to save money and time by manufacturing a variety of laboratory equipment directly [13]. In the problem that is solved in our article, 3D printing is not something new. For example, a 3D printed (using an FDM printer) system for producing spheroids using the hanging drop method has already been described [14].

However, FDM printing is a relatively rough technology for producing such miniature objects as microwells for cell spheroids. It is possible to use such technology as micro-CNC to make master molds from which the impressions will be made [15], but the such equipment costs a lot of money, and its operation requires very qualified specialists. Hence, the preferable method is much more precise the FDM and relative affordable photolithography 3D printing of various types. Unfortunately, in this area of research, authors rarely delve into the details of the printing process itself, which makes these approaches difficult to reproduce. For example, there is an article about a 3D printed stamp for the production of microwells in a 96-well culture plate [16], but the authors do not report the printing process in detail, limiting themselves to only mentioning the company that made the stamp. From the illustrations, it can be assumed that the technology used was laser stereolithography (SLA), which is an expensive method, compared to MSLA, which has now become substantially cheaper. In another research project such stamps were made with a professional MSLA 3D printer, but the cost is extremely high (more than 150,000 USD) which makes this product unaffordable for many laboratories [17]. At the same time, the features of the printing process are not reflected in this work either. A slightly different technique was used by the authors of the article, who used a 3D printing form (made using AutoDesk Ember STL, the price of which is approximately 6000 USD) [18] to produce wells in a polyacrylamdine gel applied to a glass substrate in which they were culturing spheroids. In a recent paper [19], the authors used a 3D printing stamp to make deepenings in agarose, however, with the exception of the printer model (far from the cheapest FormLabs Form3, USA) they did not provide any details about the printing process or the design features of their stamps, which limits the ability to reproduce this approach. It is worth noting that there are some notable exceptions. For instance, in the article [20] the authors describe the fabrication of microwells from PDMS using DLP 3D printer (approximately $19,000) and the authors have provided a comprehensive illustration of the printed objects in the supplementary materials.

In this article, we set out to study the limitations and possibilities of low-cost MSLA 3D printing technology (below 500 USD) and develop a set of approaches to affordable and customizable production of cell spheroids. We also emphasize the importance of documentation, not only showcasing what we have done but also providing detailed instructions on how to replicate and adapt it to your specific needs, highlighting potential pitfalls along the way. The article and supporting materials show the designs of stamps for the production of microwells in a glass bottom dish, in 96 well plates, as well as molds for mass production of spheroids in 6 well plates and in culture flasks. STL files and editable 3D models of all objects described in the article are given in supplemental materials.

## Materials and Methods

### 3d printing

The models were designed in the open-source software FreeCAD (0.21.0 and 1.0 in the late stages of development), and the Chitubox (1.9.4) slicer was used for printing.

The MSLA 3D printer Anycubic Photon Mono was used in the work (pixel size 50x50 microns, resolution 1620x2560, print area (130x165x82 mm). It is possible to use any other MSLA 3D printer available on the market. We chose Anycubic Basic Translucent Green resin because it allows us to study objects with optical transmission and LCSM microscopes. Specific printing settings were selected by printing test objects (“Cones of Calibration” test was used) and are given in the supplemental materials (section S1). The authors note that the settings may vary depending on the printer model and resin and must be tuned for specific tasks and conditions.

After printing, the parts were separated from the bed and washed with isopropyl alcohol. Initially, Firstly, in the portion of alcohol in a plastic container, then in the fresh alcohol in the Cure and Wash Station (15 minutes) and, finally, completely cleaned using a rinse bottle. Alcohol was removed from the printed products with compressed air from compressed air dusters, after which the products were cured, also using a Cure and Wash Station. Finished products were stored in plastic zip-lock bags. Stamps for 96 well plates were kept in empty boxes of 10-200 µl tips (Figure S1).

### Silicone molds

For the manufacture of secondary molds, two-component cast silicone based on tin Alcorsil 315 (China) was used. The silicone base was mixed with the hardener using a wooden spatula in a ratio of 100:2 base to hardener for 4-5 minutes. A dye was added to visually assess the quality of mixing. In addition, the dye allows to create a color-coding system to separate finished molds by well size (it is impossible to sign silicone products with a sharpie). Next, the silicone was poured into a printed mold, evenly distributed with a wooden spatula and vacuumed for 5-7 minutes in a vacuum chamber (lowest pressure was approximately 6 kPa). Vacuuming removes small air bubbles that spoil the finished product. During vacuuming, some of the silicone may be lost, and to compensate for the losses, another smaller portion of silicone with a hardener was added to the mold. In this case, you can use silicone with the addition of another dye, which increases the color palette available for marking finished silicone molds. The cast silicone thickens within 20–40 minutes, depending on the temperature, and acquires final strength in 24 hours. After this, the mold was extracted and washed with isopropyl alcohol and stored in zip lock bags to protect from dust. A video demonstrating the process of making molds and removing them, as well as the protocol are attached to the supporting materials and uploaded to GitHub.

### Optical and confocal microscopy

Macrophotos were taken with the macro camera of the Redmi Note 10 Pro mobile phone (MIUI Global 14.0.2).

For microscopy of test objects, stamps, wells, spheroids and cell cultures, an Olympus IX-71 (Olympus, Japan) inverted optical microscope with a Toupcam (ToupTek Photonics, China) UCOMS03100KPA digital camera with ToupView (4.11.19782.20211022) software was used.

Also, the objects were analyzed with the help of a laser scanning confocal microscope LSM-710 (Carl Zeiss, Germany). This method allows to examine the objects in three dimensions, which is critical for microscopy of large objects such as printed mold parts and spheroids. Z-stacks of 3D printed objects were observed at an excitation wavelength of 405 nm and emission detection in the range of 410-550 nm, using an EC Plan-Neofluar 10x/0.30 M27 objective. During microscopy, the brightness correction function was used to correct the height of the subject to acquire the optimal image.

To observe the formation of a spheroid in dynamics, cells were cultured in an Incubator PM S1 (Carl Zeiss, Germany) at 37C.

### Cell culture

Rhabdomyosarcoma cells (RD), Human Skin Fibroblasts (HSF) were obtained from Cell Bank of Institute of Cytology of the Russian Academy of Sciences (Russia), MSC ASC52telo hTERT, HEK-293t, MCF-7, A541, HeLa, Hacat and Vero cells were obtained from Cell Bank of MSU (Russia). Cells were cultured in a T25 flasks (JetBiofill, China) or 6 well plates (JetBiofill, China) in DMEM (BioInnLabs, Russia) containing 10% fetal bovine serum (Intl Kang, China). Cell cultures were maintained in an incubator at a temperature of 37°C in a humidified atmosphere with 5% CO_2_. Cells were passaged after 70% confluency of the monolayer, every three days and every five days for MSC ASC52telo hTERT and HSF cultures. Cell detachment was performed using 0.25% trypsin-Versen solution (BioInnLabs, Russia). HEK-293t culture does not require trypsin solution and was separated by vigorous shaking in PBS after washing. Cells were routinely checked for mycoplasma.

### Agarose molds production

We measured the sizes of culture plastic (dishes, plates, etc.) and showed significant differences, down to a millimeter from one manufacturer to another. The only truly reproducible value is the distance between wells in culture plates (9 mm for a 96 well plates). Accordingly, all stamp sizes, agarose volumes, and other things may vary slightly when using plastic other than ours. Key dimensions are given in the supporting materials (section S2) and in the project’s git repository, .stl files of stamps and molds are given, as well as .fcstd (freeCAD software format) and .step files that can be edited to the size of the culture plastic used, which requires a certain amount of trial and error.

For the production of molds in glass bottom dishes in a 6 well and 96 well plates, 2.7% agar (Panreac Applichem, Spain) or agarose (Helicon, Russia) was used, both prepared in phosphate-buffered saline (Bioinnlabs, Russia). To prepare solution 810 mg of agar/agarose was weighed into a 50 ml test tube, then 30 ml of PbS (pH 7.4) was added. After shaking, the agarose was gently heated in the microwave until melted (90-95C).

Two approaches were used to construct the agarose molds. First, agarose was poured into a culture plastic (dish, culture plate), and then a MSLA printed, or silicone stamp was placed there. About 1.5-2 ml of agarose should be poured into glass bottomi dishes (here it is convenient to use a disposable Pasteur pipette). For a 96 well gel was dispensed into 96-well plates using a multi-channel pipette (50 µl each), drawn from a standard polypropilene reagent reservoir (JetBiofill, China).

After agarose solidification (about 10 minutes at room temperature around 25 degrees) the stamp was carefully removed. This approach was used to make molds in glass bottom dishes to visualize spheroids using CLSM and in 96-well plates for cytotoxicity tests.

In another option, molten agarose was poured into a silicone mold. To avoid the formation of small air bubbles, the silicone mold under the gel layer was spread out with a 10-200 μL pipette tip. After cooling, the agarose mold was carefully separated and transferred to a culture container and filled with culture medium for 30 minutes. This was used to make spheroids on a large scale in six-well plates or T-125 Tissue Culture Flask with re-closable lid (TPP, Switzerland) flasks. When using molds with microwells smaller than 400 µm, due to surface tension the liquid did not fill all the microwells and bubbles were captured below the layer of liquid (section S4). In this case, the agarose mold was briefly washed with isopropyl alcohol to reduce surface tension and then rigorously washed three times with DMEM.

### Spheroid fabrication

In order to get the necessary number of cells, they were preliminarily seeded in six-well plates, where they were grown to 70-80% confluency. Cells were then detached using trypsin-Versene solution and centrifuged in microtubes (60RCF, 5 min, 4C) to remove excess trypsin, which can negatively affect spheroid formation. After removal of trypsin, the number of cells was counted using a hemocytometer and the suspension was diluted with nutrient medium to the required amount per microwell. Cells were transferred into large molds (in vials, in dishes, or in six ell plates) using a 1 ml single-channel pipette, carefully distributing the cell suspension over the entire area. In the case of small molds in 96-well plates, the cell suspension was transferred to a polypropylene reagent reservoir and then dispersed into wells using a multi-channel pipette (50-70 ul of cell suspension per well). Cells fell into microwells either by gravity or using a vortex with a plate rotor (at minimum possible speed). Cell seeding was monitored using a microscope. If there were too many cells in the free space outside the microwells, the excess cells had to be removed. For this purpose, the plate or dish was tilted and the medium together with the cells suspended in it was removed, after which new nutrient medium was added to the plate. Protocols and illustrations showing this process can be found in the supplementary materials (Section S5) and on the GitHub.

For comparison with commercial products, spheroids were also made in 3DSphearo dishes (Jet-Biofill, China) and using commercial silicone molds Microtissues (Microtissues, Inc., USA). The dishes and agarose molds made with Microtissues (according to the manufacturer’s protocol) were seeded with an amount of cells equal to those in a silicone mold of the same area.

### Spheroids staining for confocal microscopy

Spheroids were fixed with 4% formaldehyde for 1 hour. Before and after fixation, spheroids were washed twice with a PBS. The fixed spheroids were stained with Hoechst 33342 (Bio-Rad Laboratoties, USA) (10 μl of the dye (1.1 mg/ml) were dissolved into 190 μl of PBS, add 200 μl to the well for 1 hour).

### Optical clearing of spheroids

To improve the quality of the images obtained with CLSM the spheroids were subjected to an optical brightening procedure, for this purpose different protocols described in the literature were used. To show the potential of the microwell approach, optical brightening was performed directly with spheroids in agarose microwells in the wells of glass-bottomed dishes.

**Incubation in 88% glycerole solution** [21]

A 99.5% solution of glycerole (Reachem, Russia) was used for the preparation of a saline solution. The agarose mold containing spheroids was filled with 2 ml 88% of the glycerole solution and was incubated in the dark at room temperature. The clearing solution was changed twice for every day of incubation. Clearing takes at least 48 hours.

**ClearT method** [22]

Spheroids were incubated sequentially in 20, 40, 80, and 95% (v/v in PBS) formamide solutions (Fluka Analytical, Sweden) for 5 min each. Finally, incubate in a fresh 95% solution, then leave in the dark at room temperature for 15 minutes.

**ClearT2 method** [23]

The ClearT2 №1 clearing solution with a volume of 13 ml, contained 25% (v/v) of formamide and 10% (w/v) of PEG 8000 (Panreac, Spain). Clearing solution ClearT2 №2 with the same volume contained 50% formamide (v/v) and 20% (w/v) PEG 8000. Both solutions were carefully mixed until the PEG precipitate dissolved.

Spheroids were incubated for 10 minutes in solution №1, and two times for 5 minutes each in clearing solution №2. The spheroids were incubated in the dark at room temperature for one hour.

**ScaleA2 method** [24]

To prepare 13 ml ScaleA2 clearing solution, 3.12 g of urea (Diaem, Russia), 26 μs of Triton X-100 (Loba Chemie, Austria) and 1 ml of glycerole were mixed with distilled water. Thus, a solution with a concentration of urea 0.24 g/ml, glycerole - 0.1 ml/ml, Triton X-100 - 0.002 ml/ml was obtained. The solution was heated to 40°C in a water bath and thoroughly mixed. Spheroids were incubated in the clearing solution in the dark for 72 hours, the solution was changed once a day

**ScaleS4 method** [24]

To prepare ScaleS4 clearing solution in 13 ml, 5.2 g sorbitol-D (Labochem international, Germany), 3.12 g of urea, 26 µl Triton X-100, 1 ml of glycerole and 1.95 ml of DMSO were taken. Total, a solution with a concentration of urea of 0.24 g/ml, glycerole - 0.1 ml/ml, Triton X-100 - 0.002 ml/ml, DMSO - 0.15 ml/ml, sorbitol-D - 0.4 g/ml was obtained. The mixture was also heated to 40°C in a water bath and thoroughly mixed. It took 72 hours to incubate the spheroids in the clearing solution, the solution was changed once a day.

### Cell viability assays of spheroids

To study the possibility of cytotoxic assays on the spheroids in microwells, we used DMSO (PanReac Applichem, Spain), different substances with cytotoxic and cytostatic properties: paclitaxel, everolimus, doxorubicin, topotecan, docetaxel, camptothecin, oxaliplatin (Thermo Fisher Scientific, USA) and suspensions of Fe@C-NH2 and Fe@C-COOH magnetic nanoparticles. DMSO was mixed with the culture medium to the desired concentration. Cytotoxic compounds were dissolved in DMSO (except for oxaliplatin, which was dissolved in saline (NaCl 0.9%) solution and subsequently mixed with culture medium to the required concentration. Magnetic nanoparticles were synthesized and modified in the Laboratory of Applied Magnetism of Institute of Physics of Metal RAS, the synthesis was carried out using the gas-phase method [25], surface modification was done using aryl-diazonium derivatives [26]. The nanoparticle suspension was filtered before adding to the cells through 0.22 μm PTFE syringe filters (JetBiofill, China) and diluted with nutrient medium to the desired concentration.

To study the effect of the known pharmacological agent, spheroids were formed in microwells in a 96-well plate according to the method described above. The drug solution (70 µl) was added to the well, the concentration was calculated taking into account the volume of gel occupied in the well. Incubation with the substance was carried out for 72 hours, after which the metabolic activity of spheroids was evaluated using the resazurin or MTT assay.

#### Resazurin assay

After incubation with the substance, a solution of resazurin (Thermo Scientific, USA) in PBS at a concentration of 0.42 mM was added to the wells of the plate at a concentration of 20 µl per well. The medium was removed from the corner wells and DMSO was added for 5 minutes to kill the cells to use these wells as blank. The DMSO was then removed, the wells were rinsed with phosphate buffer and fresh medium was poured into them in the volume corresponding to the other wells. Chemidoc MP (Biorad, USA) was used to record the fluorescence intensity in three spectrum bands: Green Epi with Green 605/50 filter to measure intensity of resafurin fluorescens, Red Epi with 695/55 filter for resazurin and Blue Epi with 530/28 filter to get a picture of the plate itself. The obtained images were processed using CellProfiler (4.2.5) [27] to segment the cells of the plate and measure the fluorescence intensity in each of them.

#### MTT-assay

To obtain the blank data, the optical density of the wells with spheroids was measured using a plate reader (Elx808, Bio-Tek Instruments, USA) at a wavelength of 570 nm (optical density from the plate itself, agarose and spheroids).

One day after incubation, the culture medium was replaced with an MTT solution at a concentration of 1 mg/ml for 4 hours. After incubation, the nutrient medium was removed with a multichannel pipette and 100 μl of DMSO was poured into the wells of the plate, extracting the colored reaction product from the spheroids. With DMSO, the spheroids were incubated for 60 minutes in an incubator at 37 degrees, after which the optical density was recorded using a plate reader.

### Image processing

#### Image processing

The automatic processing of images of both stamps and cell spheroids was conducted using the CellProfiler (4.2.5) and Ilastik [28] software. As a rule, CellProfiler was used, but in cases when satisfactory segmentation could not be achieved, Ilastik (1.4.0) was utilized. Manual image processing was performed using FIJI (1.54f) [29].

The effect of substances on spheroids viability was also evaluated not only by metabolic tests, but also using visual data. For this purpose, spheroids in each group (substance, drug concentration and incubation time) were imaged with a microscope. Since it would be quite labour intensive to take pictures of each one, a random number generator was used to select some of them. Images were acquired every 24 hours. The images were then segmented by ilastik and analyzed using CellProfiler. Built-in functions were used to evaluate shape, size, texture, and granularity in the images; the importance of specific visual features for spheroids after vairous incubation times with different substances was evaluated using the ML approach. For each substance (or substance concentration), the most important features were determined using a random forest method. This is an ensemble algorithm consisting of multiple solver trees. The difference in accuracy of models with different hyperparameters was determined by cross-validation with F1_score metric on the training sample. The most accurate models were selected. The quality of feature selection by importance and distribution of features by classes was assessed on test data.

#### 3d model processing

In order to clarify some peculiarities of MSLA printing, we present images simultaneously comparing the model created in CAD, the model obtained after the slicer and the real appearance of the object obtained with the help of CLSM. Obtaining these images and corresponding videos turned out to be a relatively non-trivial task, so we decided to describe our pipeline in a bit more detail.

To illustrate the features of MSLA printing, the work contains images that simultaneously compare the model created in CAD, the model obtained after the slicer, and the real appearance of the object obtained using CLSM. One of the approaches we used was to export a .stl file from the CAD and then use the stl-to-voxel (0.9.3) library for Python to convert it into a layered set of .tiff images (with resolution=512). Using a slicer, the same stl model was converted into a .pwmo file, which was again converted into a .tiff set using UVTools (4.3.2). After collecting images with a confocal microscope, a multilayer .tiff file was also obtained. In the ImageJ program, the images were manually aligned relative to each other and transferred to napari (0.4.19) [30], where 3D images and animations were already produced. The attenuated_mip rendering mode was used for CAD and slicer models, and mip for images from a confocal microscope.

Another approach involves using Blender (4.1) with the tif2blender (0.1.1) plugin. This plugin made it possible to import both data obtained from a confocal microscope and a 3D model converted to tiff with UVTools, as in the previous method. In Blender, the 3D printed object model was converted into a mesh, and the microscopic data was visualized using the emission volume shader.

### Repeatability

To check the reproducibility of the approach described in this article 3d models of the stamps, printed stamps, silicone molds and all the instructions were provided to independent laboratories at other institutes for testing. These laboratories worked with different culture plastic, different agar/agarose, and various cell cultures. Feedback from these organizations was used to update the protocols and make them more accessible.

## Results and discussion

### Capabilities and limitations of the MSLA 3d printing

MSLA (masked stereolithography) printer is based on stereolithography to produce three-dimensional models. The process is carried out through the polymerization of photosensitive substance (resin) under the influence of UV light, and the mask controls which part of the field will be illuminated. During the printing process, a platform (bed) is lowered into the resin-filled vat. Light from photodiodes is then projected through an LCD screen onto the bed. The screen’s individual pixels control the passage of light, functioning as a Mask (the “M” - in “MSLA”) in the printing process. After one layer of the object is polymerized, the bed is raised, fresh resin flows under it and it is lowered again, to a height sufficient for the next layer to form under the first one. The process is repeated until the object is printed entirely. MSLA is a surprisingly cheap 3d printing method, for its accuracy and print quality. Thus, it is often used in laboratories to print miniature objects

In greater detail the process of MSLA 3d printing the main complexities and limitations of this method are outlined in the supporting materials (Section S6) with illustrations and diagrams.

In order to determine the limits of the technology in terms of the shapes and forms needed to produce microwells, a test object was printed containing many deepenings and elevations of different sizes and shapes: cones, pyramids, cylinders (Figure 1). Detailed description of geometric dimensions and features it contains is given in the supplement (Section S7). All features have been studied with CLSM, which proved to be a very convenient tool for studying such objects.

**Figure 1.**
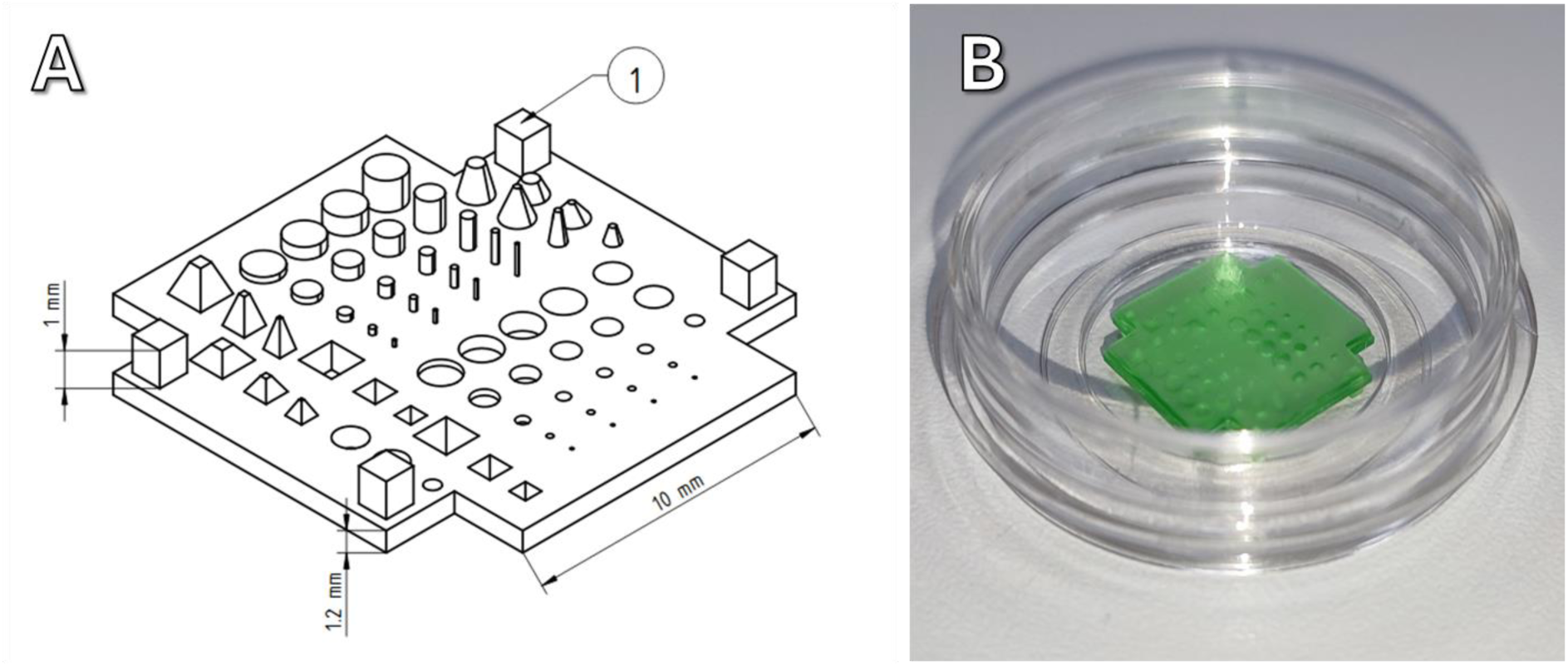
A scheme of the test object. The cubes (1) at the corners are necessary to serve as supports, because the test object is installed with the relief downwards and supports are needed to make it stand flat. Detailed dimensions of each feature are given in the supplementary materials. B - photo of the test object located in the glass bottom dish

The smallest features of the test object used are cylinders with diameter 100 μm and height 200 μm and corresponding recesses. Neither of them could be printed. The cylinders are most likely not printable due to their brittleness and simply collapse when torn from the film. The recesses have another problem - high viscosity of the resin leads to the fact that it does not flow out of the holes and is illuminated there by parasitic light from neighboring pixels of LCD screen.

Larger sized features were printed and as expected, the shape of the printed objects is significantly different from what was conceived and designed in CAD. Some examples are shown in the figure, others are included in the supporting materials. Cones (Figure 2 A, B and C) become pyramids of rather complex shape, with many irregular steps, pyramids with smooth sides become stepped. Cylinders and cylindrical recesses are distorted the least (Figure 2 D, E, F), merely acquiring a characteristic pixelated profile. Animations generated with napari that allow us to experience this in 3d are presented in the supplementary materials.

**Figure 2.**
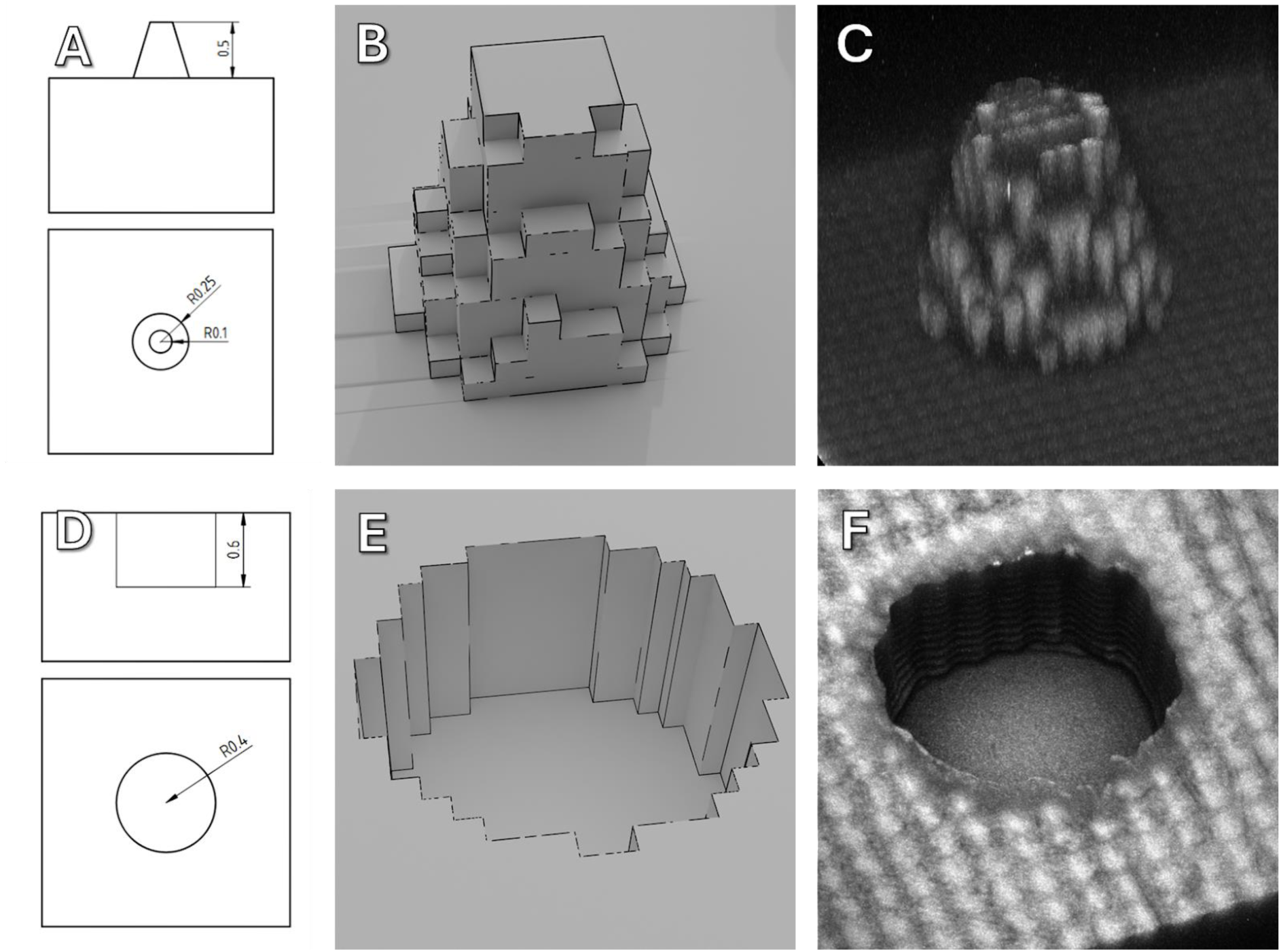
Scheme with dimensions, 3d rendering of the model formed by the slicer and 3d reconstruction obtained by confocal microscopy for objects of different geometry.

However, the fact that a printer can print single objects, even if it distorts their geometry, does not mean that it will do so reliably. Quite quickly it was discovered that when tens or hundreds of 200 µm diameter cylinders were printed, a noticeable part of them would end up damaged and broken when printed, as shown in the Figure. Another limitation is the distance between cylinders, if it is too small, bridges of cured polymer form between them during printing (Figure 3). As a result, it is optimal to print cylinders with diameters of 300 µm or more at a distance of about 300-400 µm from each other, as shown in Figure 3 (C and D).

**Figure 3.**
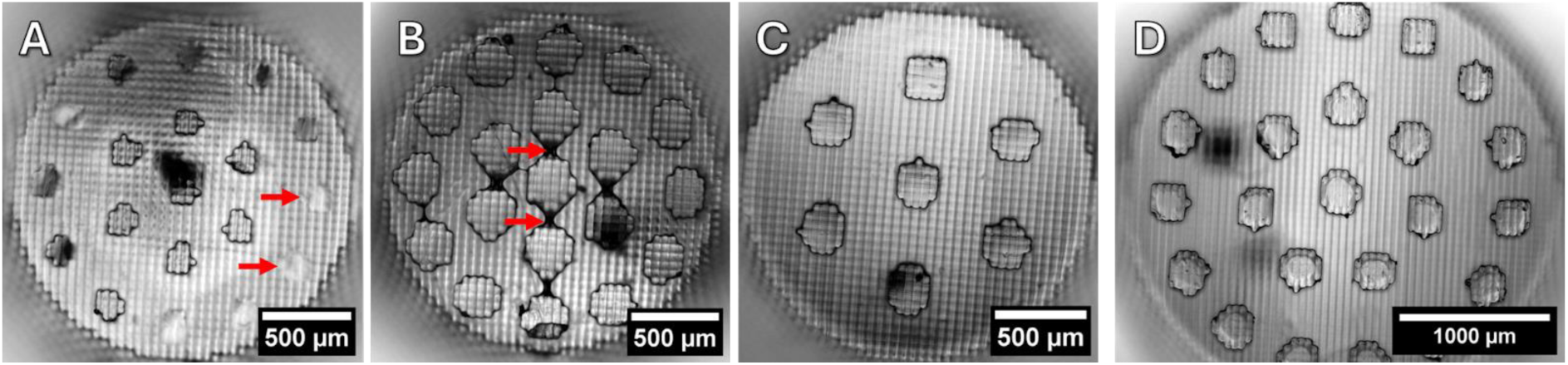
Stamps for making microwells in a 96-well plate. A - 200 µm cylinders, arrows indicate cylinders that broke off during printing; B - 300 µm cylinders with bridges between them; C - 400 µm cylinders at a greater distance from each other and D - 300 µm cylinders on a larger diameter stamp.

Similar difficulties were encountered when making silicone molds. Despite the fact that in the 3d printing mold the 200 µm diameter recesses are obtained successfully when making a silicone impression from this mold, some of the cylinders are damaged. An image of a silicone mold with cylinders of different sizes including the defective 200 µm is shown below (Figure 4).

**Figure 4.**
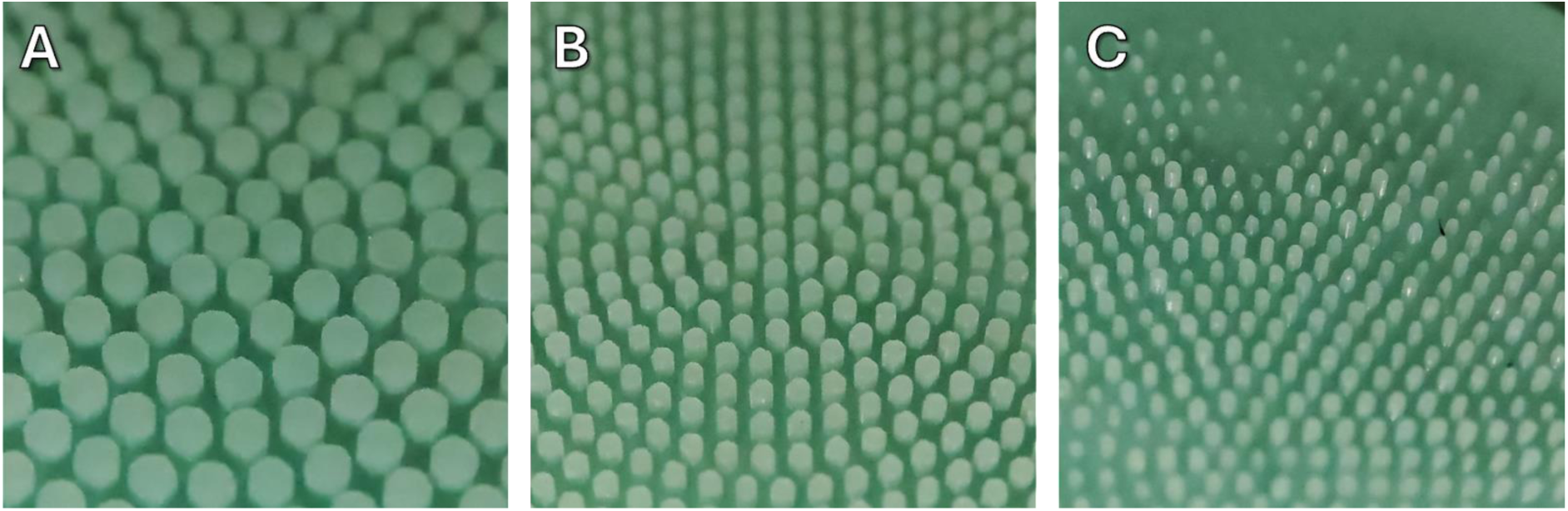
Macrophoto of silicone molds with cylinders of different diameters. A - 600 µm, B - 300 µm, C - 200 µm.

It should also be noted that in spite of the shape distortions due to the approximation of objects by voxels, the imprint area of objects is reproduced relatively accurately. To determine this, microphotographs of cylinders of different diameters were processed. For a 200 µm diameter cylinder the measured equivalent diameter was 223±11 µm, for a 300 µm diameter cylinder it was 318±10 µm, and for a 400 µm cylinder it was 446±17 µm. Details are given in the supporting materials (Section S8).

In general, we have seen that on the MSLA 3d printer it is possible to reliably print even quite small objects, which can then be used directly as stamps for creating agarose molds or for making silicone molds, in which agarose molds will subsequently be made.

### Stamp and mold design

The design of molds and stamps was created by iterative prototyping. Schemes of different versions with explanations are given in the supplementary materials, here the final designs for stamps and silicone molds are presented and described.

#### Stamp for glass bottom dishes

For confocal microscopy, specialized glass-bottomed petri dishes are often used to minimize the distance from the objective lens to the sample. Accordingly, the agarose layer should also be minimized so as not to increase the distance even further. Agarose of such thickness becomes surprisingly delicate and can be damaged by any careless movement, so the variant with the use of silicone molds for making such constructions was discarded at once and a stamp was designed, allowing to make a mold directly in a glass bottom dish or in a usual petri dish, but with an appropriate change of the stamp vertical dimensions. After a number of iterations, a stamp of the following design was developed (Figure 3).

**Figure 3.**
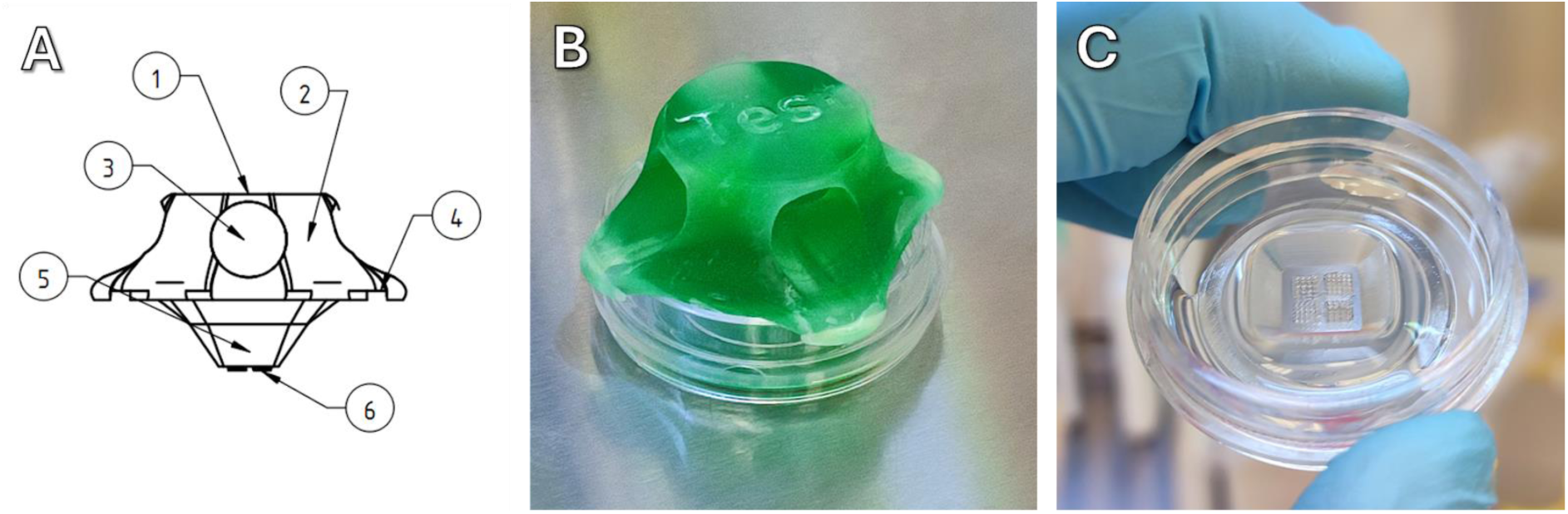
A - sketch of a stamp for a dish. 1 - flat part for marking, 2 - holes for easy grip, 3 - recesses for centering the stamp on the walls of the dish, 4 - part to be immersed in the well, 5 - working part for creating microwells. B - photo of the stamp installed in a glass bottom dish with poured agarose. C - microwells created in agarose

The stamp was designed to be printed flat on the printer’s table, without supports and easily detachable after printing. On the upper part of the stamp (in working position) there is a plane (1) on which an inscription can be embedded in the design. In the main part of the stamp (2) there are holes (3), by which the stamp can be conveniently held and, importantly, removed from the solidified agarose. The micropins (6) are located on the part (5), which is immersed in the agarose. And it is these micropins that leave deepenings in the agarose after removing the stamp.

Above we have already shown in what limits it is possible to vary the geometry of wells and now it is time to check how spheroids are formed in microwells of different geometry. All shapes and sizes can be examined, but we stopped at a few of the most representative ones. Test stamps for glass bottom dishes (Figure 4) containing micropins of different shapes were designed. These shapes are cylinders with a diameter of 300 μm (1), truncated pyramids with an upper square edge of 600 μm and a lower one of 100 μm (2), truncated cones with an upper diameter of 500 μm and a lower one of 200 μm, cylinders with a diameter of 300 μm (3), truncated pyramids with an upper square edge of 600 μm and a lower one of 300 μm (4).

**Figure 4.**
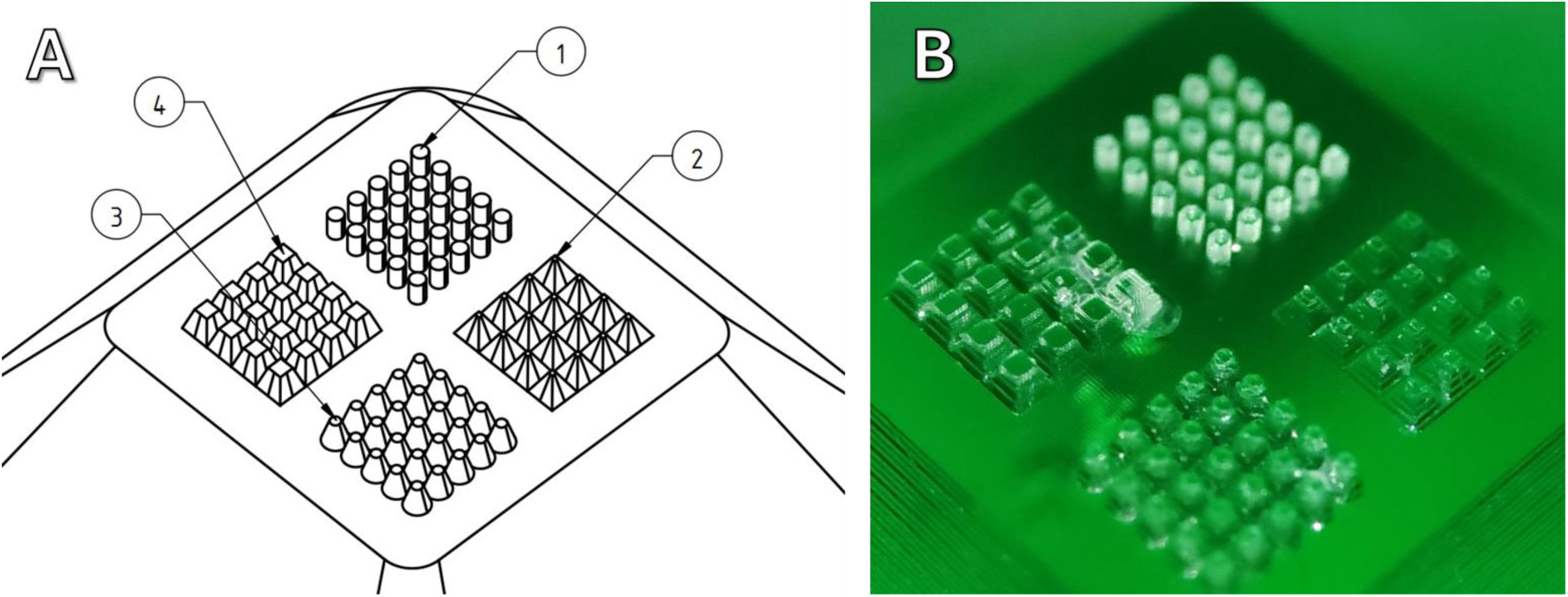
Schematic diagram of the working part of the stamp (A) for testing micropins of different shapes and its macrophoto (B). Explanations are given in the text.

As discussed above, when printing on an MSLA printer, the CAD program shapes are approximated by voxels, which leads to a noticeable distortion of the shape. It is the formation of “steps” on non-vertical walls that is most sensitive. These steps cause cells added to the microwells to get stuck.

With the help of a stamp with different shapes of micropins, the corresponding agarose molds were made in a glass bottom dish. Here it is necessary to make an important statement - agarose should be poured into the dish in a relatively little volume, for our dishes it is about 1 ml. If there is too much agarose, it sticks to the stamp too firmly after solidifying and during stamp removal agarose can be torn or removed from the dish entirely.

MSC ASC52telo hTERT culture cells with labelled nuclei were added to the prepared agarose molds, and images were then taken using a confocal microscope in z-stack mode, which allows a 3D image to be obtained (Figure 5).

**Figure 5.**
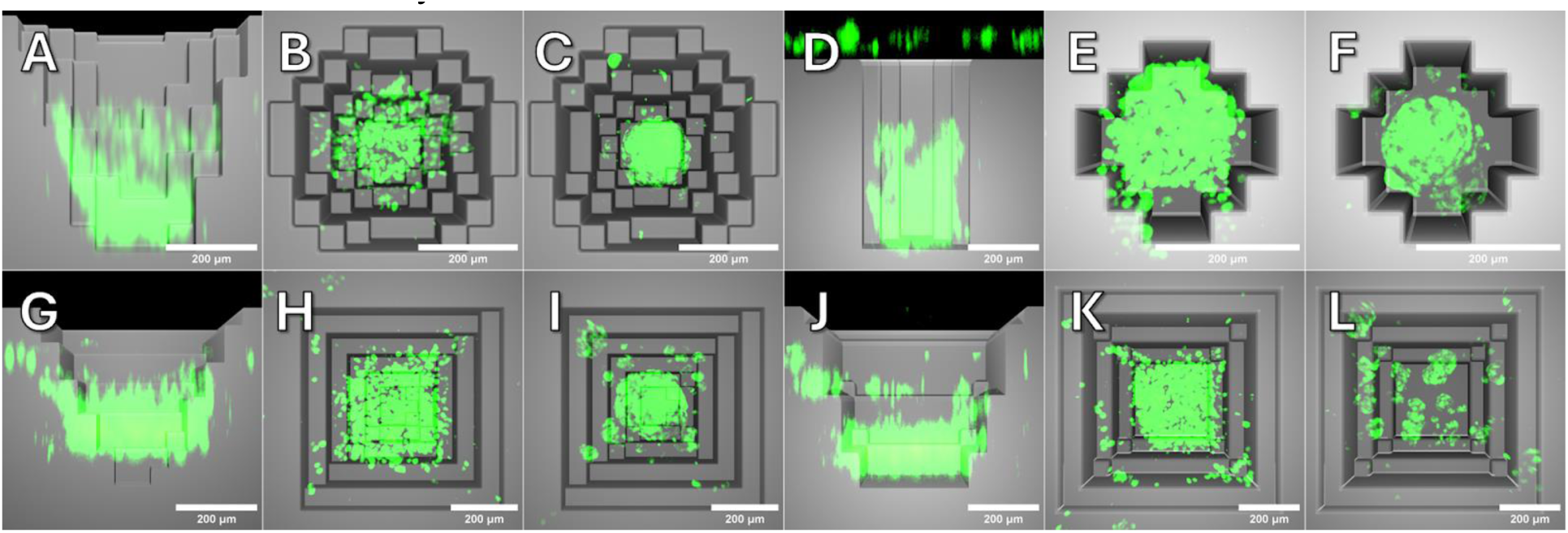
Combination of confocal microscopy (MSC ASC52telo hTERT cells, cell nuclei stained with Hoechst 33258) and 3d rendering of differently shaped microwells. The scale bar is 200 μm. All wells are 500 μm deep. Side and top views are provided In the case of the side view, half of the microwell model is cropped, for illustrative purposes. A, B and C are truncated cones with upper diameter 500 μm and lower 200 μm, D,E,F are cylinders with diameter 300 μm, G, H, I are truncated pyramids with upper square edge 600 μm and lower 100 μm, J, K, L are truncated pyramids with upper square edge 600 μm and lower 300 μm. A, D, G, J - side view of cells in a microwell shortly after seeding. B, E, H, K - top view of cells in a microwell shortly after seeding. C, F, I, L - top view of spheroids formed a day after seeding.

We were unable to get a high-quality confocal 3D image of the agarose mold itself, so the illustration below shows a 3d rendering of the microwells of the corresponding shapes overlaid with confocal microscope images. The figure shows the cells immediately after addition to the wells after the procedure of washing off excess cells and spheroids formed in the same wells after a day of incubation.

In cone-shaped microwells, the cells are partly stuck on the steps, but in a day all the cells gather at the bottom forming one spheroid. In cylindrical wells the picture is similar, although it should be noted that some cells remain on the surface formed by the flat part of the stamp. In pyramidal wells the picture is different, quite a lot of cells get stuck on the steps and not all of them fall to the bottom. In a pyramid with a bottom square edge of 100 μm, besides one large spheroid, several small spheroids were formed, and some cells did not become part of the spheroid at all. In the pyramid with the bottom cube edge of 300 μm several small spheroids were formed.

For reliability and reproducibility of biological studies, it is optimal to have one spheroid of known size per well. Accordingly, we subsequently used cone- and cylinder-shaped microwells.

Being placed in microwells, other cell cultures formed spheroids in a similar manner. Some, such as Hacat keratinocyte culture or HEK-293t kidney epithelium culture, formed dense spheroids in which individual cells were almost invisible in a normal microscope. Others, such as HeLa or A-549, formed much looser spheroids without such dense intercellular contacts, which is generally consistent with literature descriptions (Figure 6).

**Figure 6.**
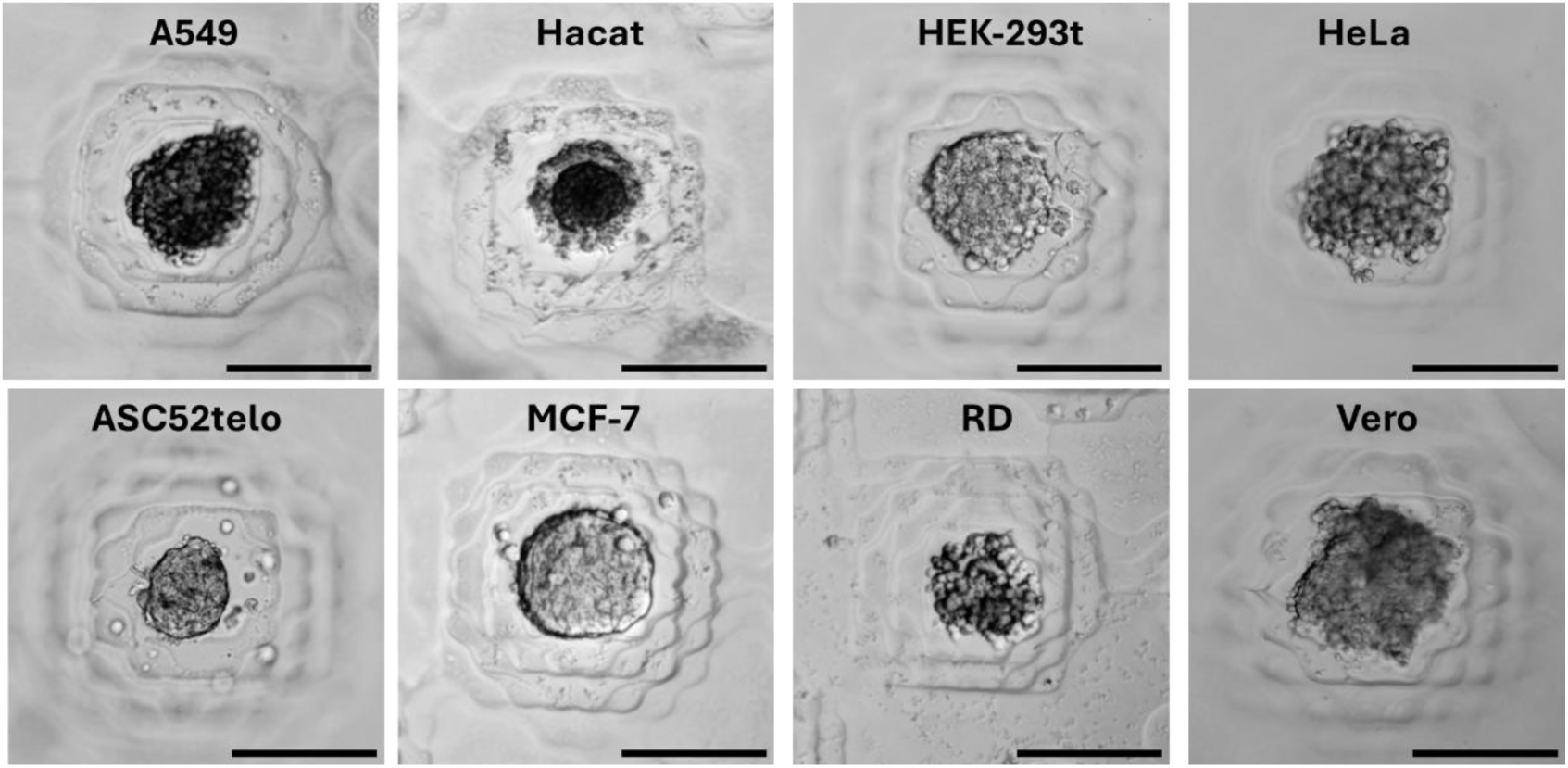
Spheroids from different cultures formed in conical microwells. Scale bar - 200 µm.

The main advantage of forming spheroids directly in the dish is the possibility to observe the process of their formation, which may also be important in some studies. The corresponding experiment was performed. We seeded MSC ASC52telo hTERT culture cells in microwells made in the glass bottom dish and observed spheroid formation for several hours (Figure 7), the video of the process is given in the supplementary materials.

**Figure 7.**
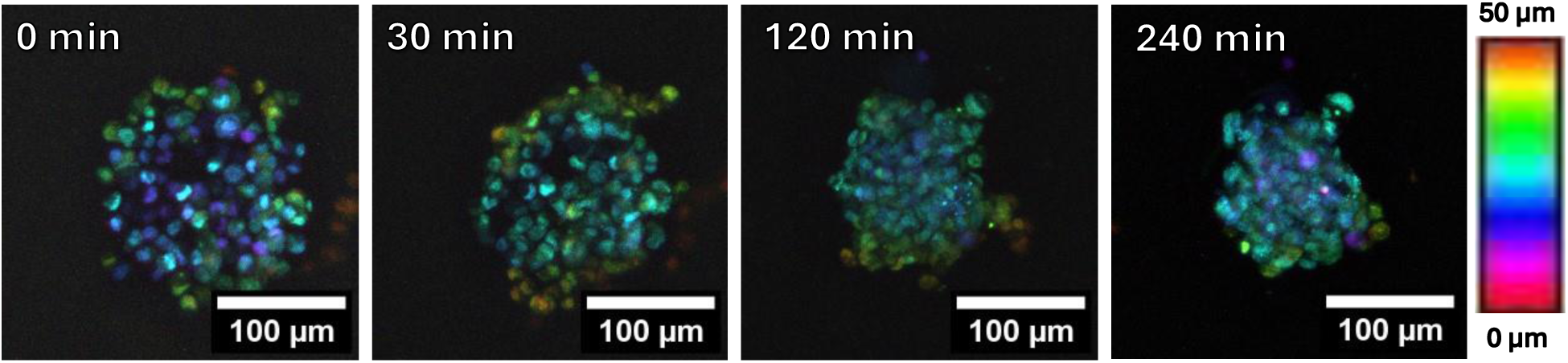
Spheroid formation in a microwell, z-stack. The color-bar on the right allows one to get an idea of the height distribution of cell nuclei.

Of course, observation of the dynamics of spheroid formation and development can be carried out for a longer period. For example, the figure below shows the complete life cycle of a spheroid. First, individual cells seeded in a microwell, then their consolidation into a spheroid and some increase in size due to cell division. About the third day, a necrotic core [31] begins to form inside the spheroid, arising from the deficiency of nutrients in the depth. During this experiment, the nutrient medium was not consciously changed to a fresher one, and accordingly, on the seventh day, when all nutrients are exhausted, the spheroid decomposes into separate non-viable cells (Figure 8).

**Figure 8.**
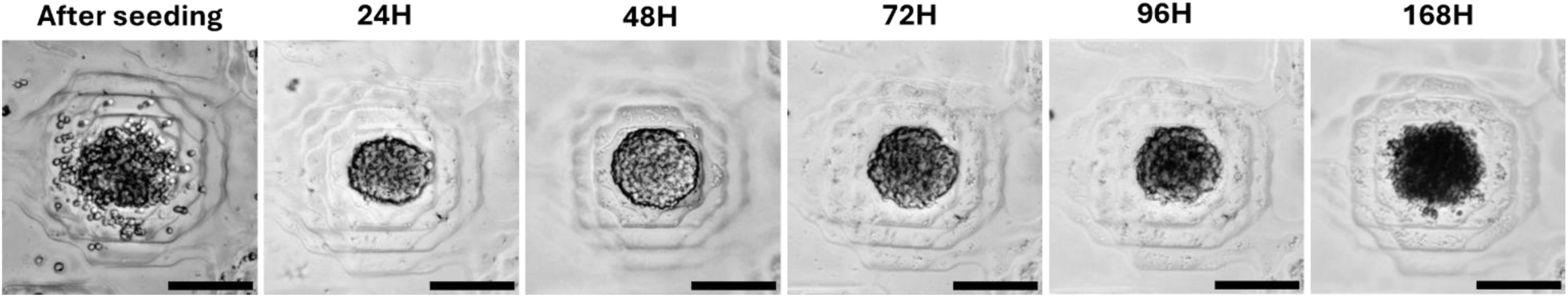
Spheroid development from HEK-293t culture cells in a microwell for a week. Scale bar – 100 um.

#### Spheroids optical clearing

The main problem with studying spheroids using optical microscopy is their optical properties: they are not very transparent. One way to overcome this limitation is optical clearing - a process in which an object is treated with different agents to remove the colored substances and to make its RI (refractive index) equal to the RI of the medium that is optimal for the optical system.

Of course, many such methods have been developed, but we want to simplify and speed up high-throughput analysis on spheroids. We have demonstrated the ability to clear the formed spheroids directly into gel molds formed in a dish.

We tested 5 different optical clearing methods on spheroids of human skin fibroblast and rhabdomyosarcoma cell cultures (Figure 9). In both cases, there were control groups of spheroids that were stained but not fixed. We evaluated not only how the optical properties of the spheroids changed, but also how the clearing process affected their size, but these results are too extensive and have been moved to the supplementary materials (Section S9).

**Figure 9.**
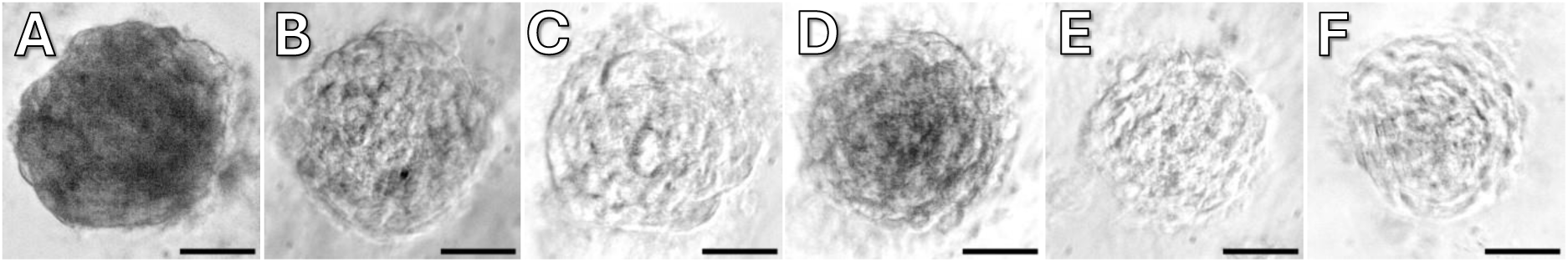
Spheroids of human skin fibroblasts cleared by various methods: A - Control uncleared spheroid, B - Spheroid after clearing with glycerol, C - Spheroid after ClearT, D - Spheroid after ClearT2, E - Spheroid after ScaleA2, F - Spheroid after ScaleS4. Scale bar - 100 μm

The CLSM images of selected spheroids were also further analysed. It can be noticed that a spheroid with a diameter of more than 200 μm forms a central dark region from which it is impossible to detect the dye fluorescence signal. This can theoretically lead to incorrect conclusions about the processes occurring inside the spheroid (Figure 6, A). In spheroids treated with clearing solutions, internal optical sections are uniformly visible (Figure 10, B–F).

**Figure 10.**
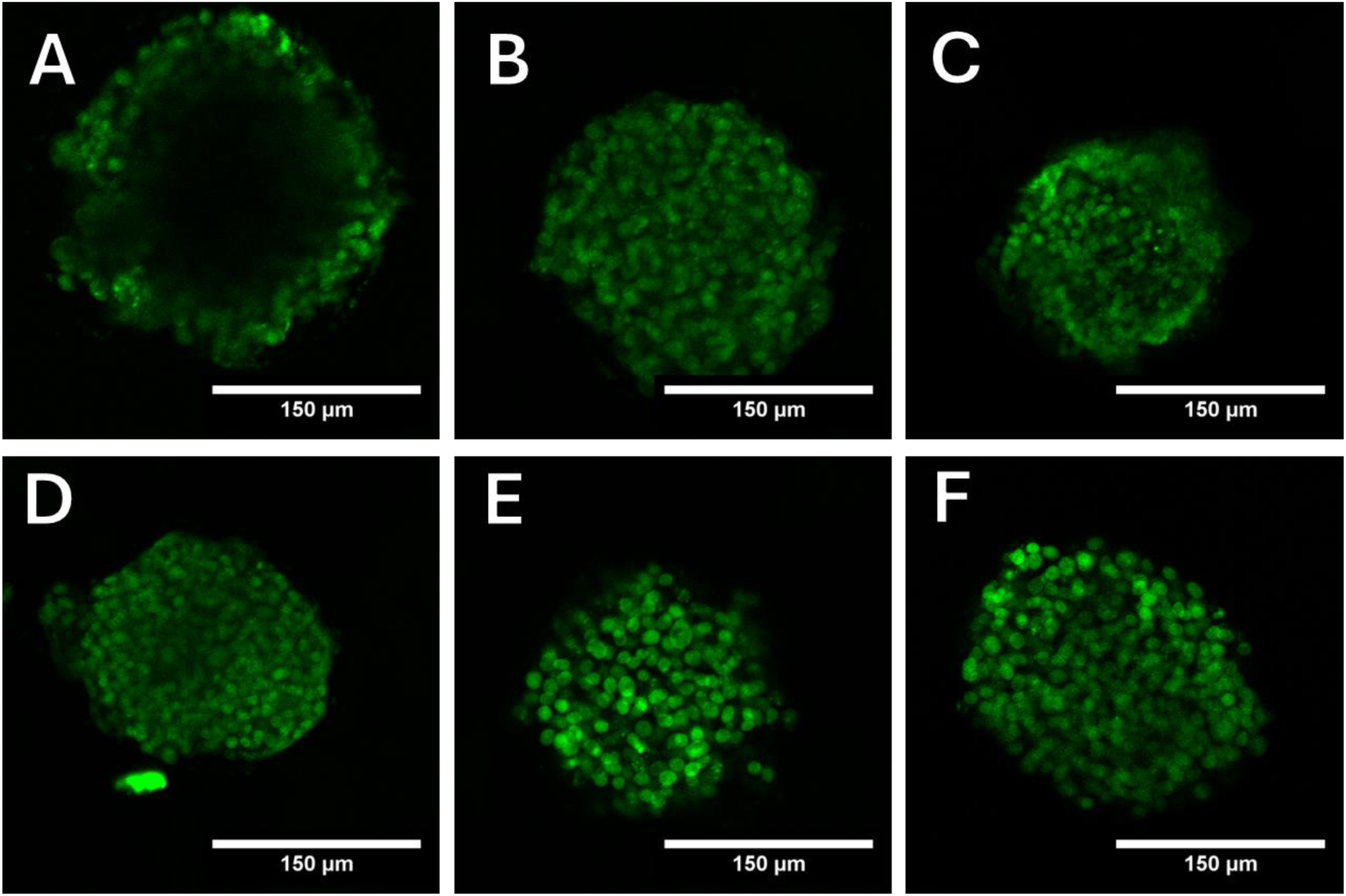
Fluorescent visualization of spheroids of rhabdomyosarcoma of different optical clearing methods: A - control, B - glycerol, C - ClearT, D - ClearT2, E - ScaleA2, F - ScaleS4.

A similar pattern is observed in spheroids of human fibroblasts. Additionally, it is noted that the spheroids from ScaleA2 and ScaleS4 exhibit more distinctly outlined nuclei that stand out against the background. It is suggested that this may allow counting them even in dense fibroblast spheroids.

A characteristic directly reflecting the efficiency of optical clearing of biological tissue is the thickness of the outer layer that can be cleared (clearing depth). Using ScaleA2 and ScaleS4 techniques, it was possible to increase the depth of the fluorescent dye signal by 85 μm. Overall, the average clearing depth of rhabdomyosarcoma spheroids was 57 μm. With each of the five techniques used, it is possible to significantly increase the thickness of the spheroid’s outer layer on which the dye fluorescence can be detected.

It was previously reported that optical clearing can cause a significant reduction in the fluorescence intensity of stained tissues, which can lead to incorrect results, for example, when using fluorescent proteins. For instance, formamide in ClearT destabilizes protein conformation. Displacement of water during clearing may also negatively affect fluorescence [23]. Polyethylene glycol in ClearT2 should preserve and increase fluorescence intensity by stabilizing protein structures [32].

Overall, we have shown that it is possible to optically illuminate spheroids directly in agarose molds, thus speeding up and simplifying their study.

### Molds for mass production of spheroids

Petri dishes, including glass bottom ones, are well suited for observing and experimenting on a relatively small number of spheroids. For some studies, however, it is necessary to work with a large number of spheroids and accordingly a substantially larger vessel area is needed. In this case, we are not limited by the optical features of confocal microscopy and, consequently, the thickness of the agarose mold can be noticeably larger, so we used an approach with silicone molds into which agarose is poured and then transferred to a culture vessel for further manipulation. Molds were constructed for 6 well plates and for T-125 culture vials.

The MSLA 3d printer is used to print a master model (Figure 11A) where a set of microwells of the required size and number are arranged in concentric circles. Two-component injection molding silicone is poured into this mold and vacuumed so that the silicone completely fills all the holes and forms, respectively, the silicone micropins. Then melted agarose is poured into the prepared silicone molds (Figure 11B) under sterile conditions; after solidification, the agarose molds (with many microwells) are removed (Figure 11C) and transferred into the wells of a 6-well plate.

**Figure 11.**
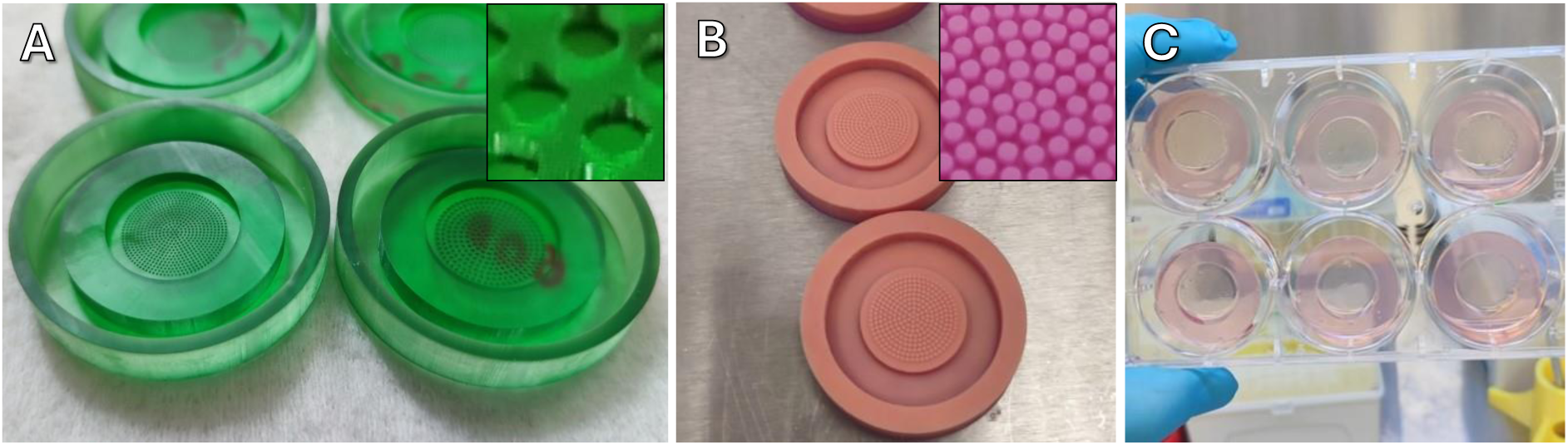
A - 3d-printed primary molds with different size and number of microwells (on the inset macrophotograph of microwells), B - silicone molds (on the inset macro image of mold’s micropins), C - agarose molds in a 6-well plate.

To demonstrate the ease and convenience of customization of microwells using our approach, we produced primary molds with different diameters of cylindrical wells: 200 μm, 300 μm and 600 μm.

As mentioned above, the printing of recesses with diameters smaller than 300 μm on MSLA printer is not reliable, respectively, silicone molds using 200 μm were also obtained with many defective micropins, but we still conducted experiments with them, although the number of obtained spheroids was less than it could be (Figure 12).

**Figure 12.**
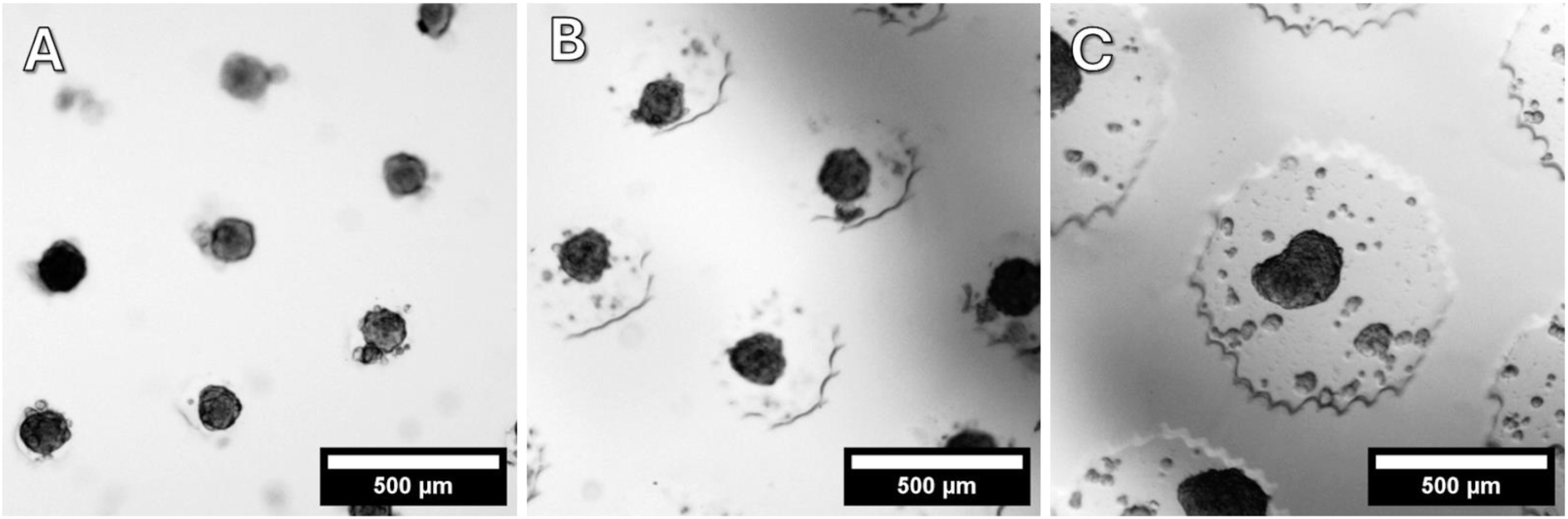
Spheroids in cylindrical wells of different sizes: A - 200 μm, B - 300 μm, C - 600 μm.

Spheroids from MSC ASC52telo hTERT cells were obtained in these agarose molds and photographed using an inverted optical microscope.

Using a combination of Ilastik and CellProfiler software, the size of spheroids and their roundness were determined (form factor - equal to 1 for a perfectly round spheroid). Evaluation of these parameters is a screening method for assessing the viability of spheroids, which was proposed in [33]. The results are presented in the figure 13.

**Figure 13.**
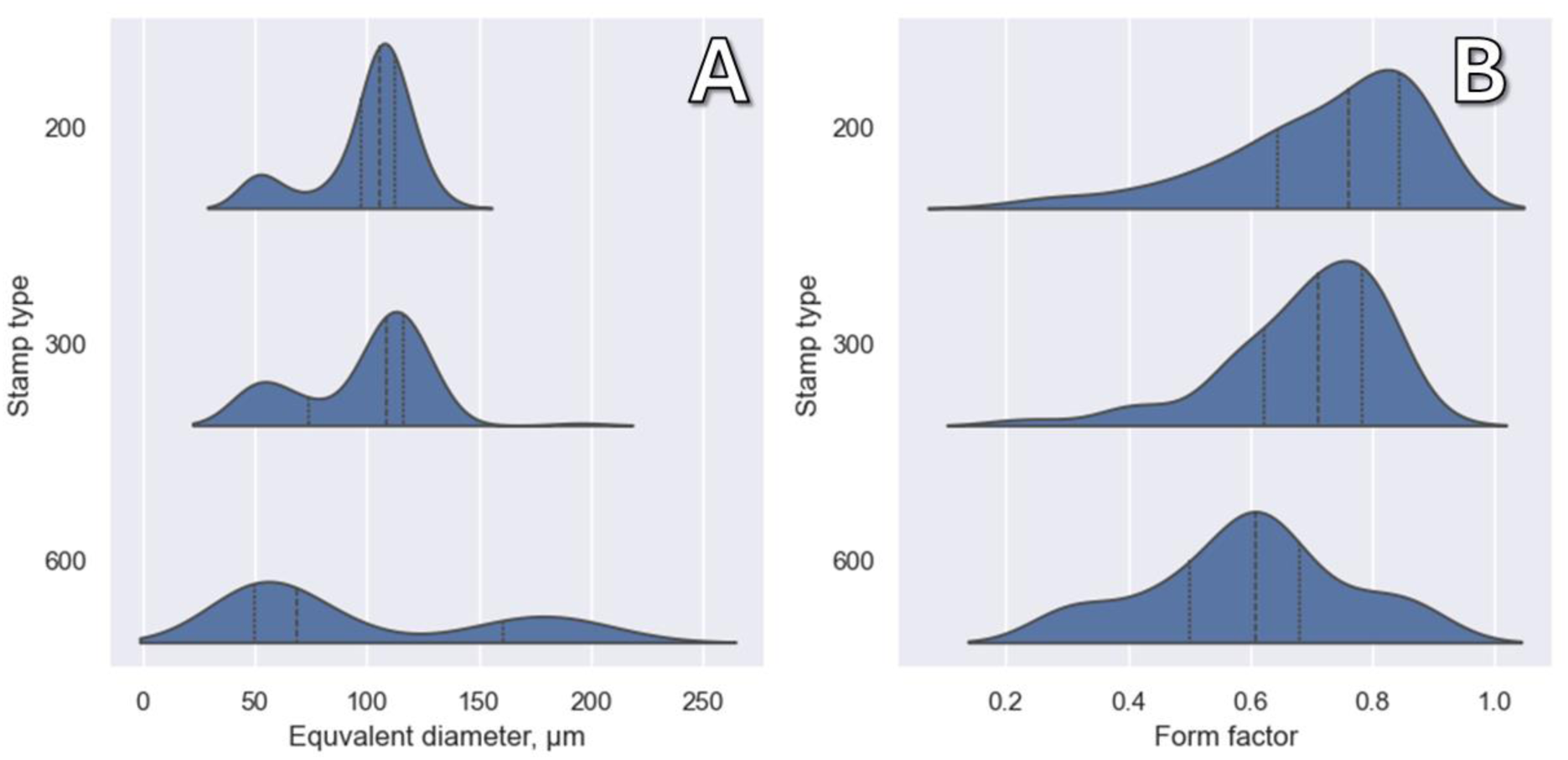
KDE-plots showing the distribution of spheroid sizes (A) and form factors (B) of spheroids in wells made with different stamps.

The size of the spheroids is always smaller than the well diameter, which was also seen in the images; at the same time, the size distribution is bimodal, with the presence of some smaller spheroids. This is especially pronounced in large (600 μm) microwells, where there are many small spheroids and fewer large ones (Figure 14). In general, large wells are not very suitable for spheroids fabrication, which can be seen from the shape factor distributions. Also a noticeable number of spheroids have an irregular shape, most likely this is because in the wells several small spheroids merge into a large one.

**Figure 14.**
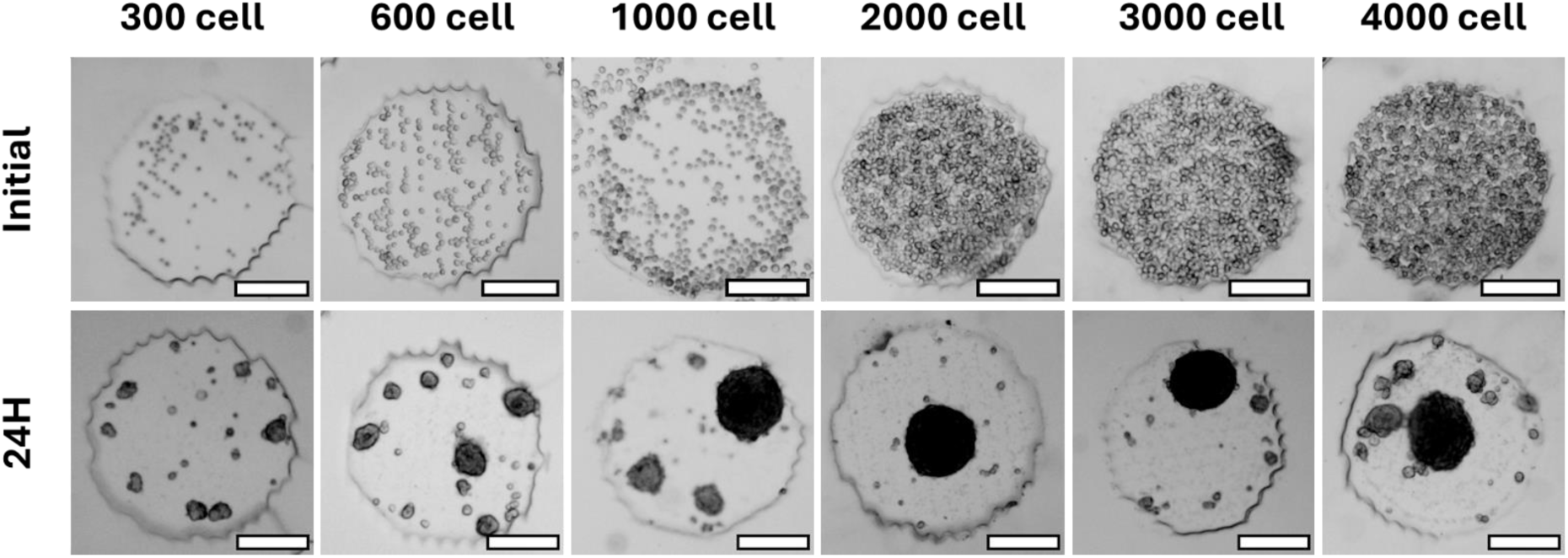
Different numbers of cells in a 600 μm diameter microwell planted in a microwell and the spheroids formed from these wells after 24 hours. Scale bar 200 μm

Another approach to adjusting the number of resulting spheroids is to vary the number of cells in the wells. We seeded different quantities of cells in a same microwells with a diameter of 600 µm (Figure).

When there are few cells, individual small spheroids are formed, consisting of tens of cells. If there are too many cells in a microwell, several spheroids or irregular spheroids are formed (resulting from the fusion of several smaller ones). And finally, there is a “goldilocks zone”, about 1000-3000 cells per well of such an area, when only one large spheroid is formed.

We also compared our system to a commercial system made by MicroTissue. These silicone molds are made using much more precise equipment than the MSLA printer and the shape of the holes is much smoother (Figure 15). However, is this a critical flaw in our approach?

**Figure 15.**
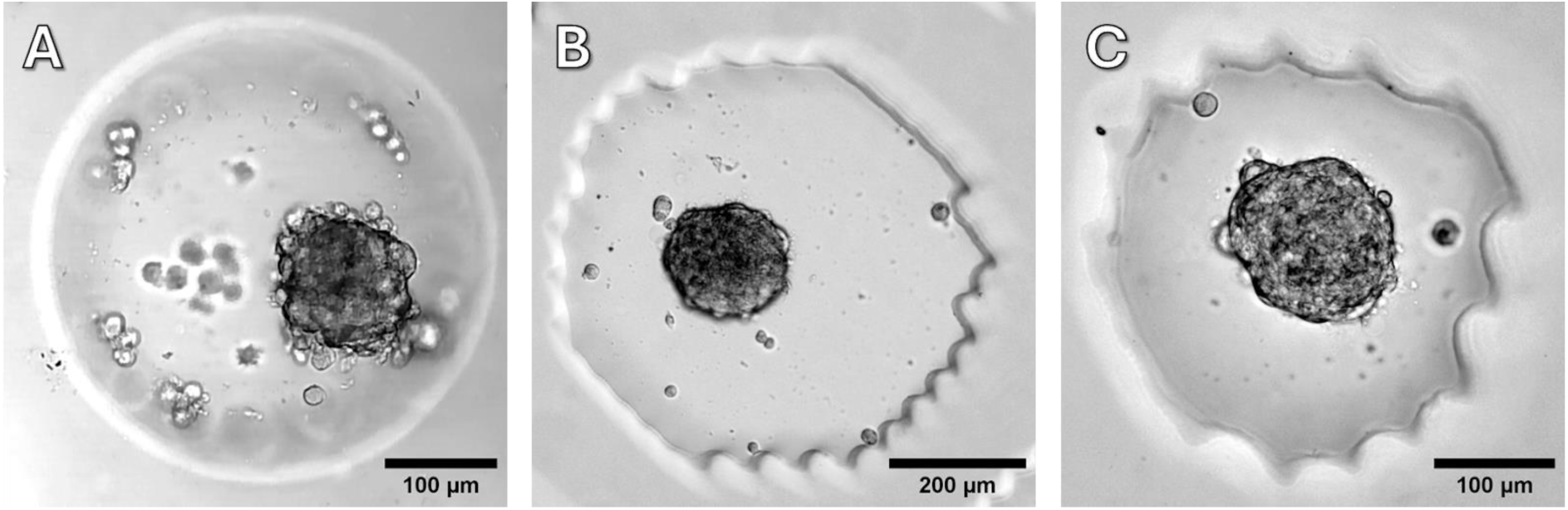
Spheroid formed in MicroTissue molds (A) with a bottom diameter of 300 μm, and in molds described in the article with diameters of 600 um (B) and 300 μm

Even though visually the wells in molds made with MicroTissue look perfectly even, the shape and size of spheroids are not significantly affected (Figure 16).

**Figure 16.**
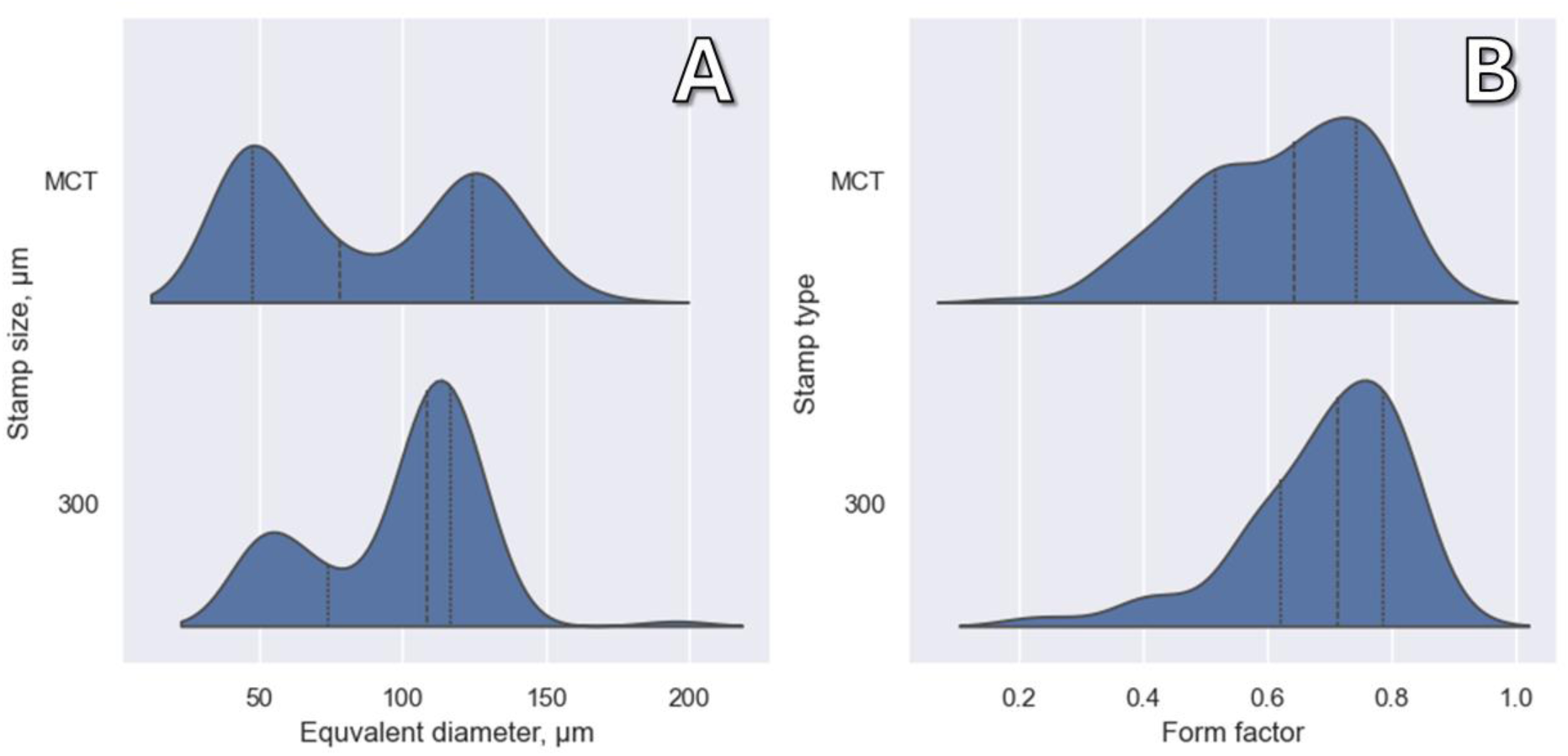
KDE-plots show the distribution of spheroid sizes (A) and form factors (B) of spheroids in wells made with MicroTissue system (denoted as MCT) and approximately similar mold of our construction with 300 μm microwells.

Another popular approach to creating spheroids is to use culture plastic with a specialized low-adhesive coating. For comparison, we used this method by adding a similar number of cells to a dish with a low-adhesive coating (in terms of dish area, as the dishes we used were noticeably larger than the molds). As can be seen from the Figure 17, under these conditions the spheroids formed irregularly shaped and chaotically distributed on the dish. When imaging and time lapse spheroids in wells, challenges can arise due to the lack of a fixed point of contact with the plastic bottom.

**Figure 17.**
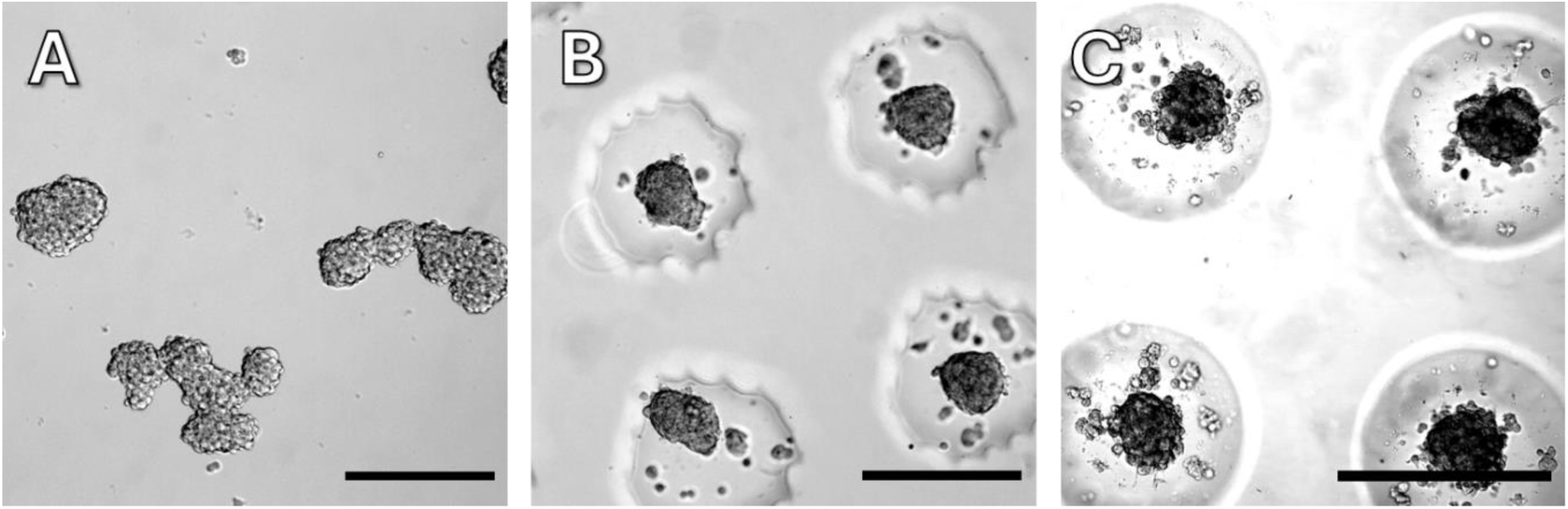
A - spheroids formed in 3DSphearo dishes, B - spheroids formed in the molds described in the article and C - in MicroTissue molds. Scale bar – 300 um.

Some studies may require even larger numbers of cell spheroids. To achieve this, spheroids can be grown in big culture vials instead of dishes or plates (the development of molds for the bioreactor [34] is definitely beyond the scope of this study). For example, cell culture flasks with a reclosable lid may be used. It is rather inconvenient to insert a stamp there due to the complex shape of the lid, in addition the printing of such a large object is also not a very easy task. It was much more reliable for us to print a mold, make a silicone cast of it and use it to make agarose molds with 14282 wells. After that two molds were inserted in the flask, allowing 28564 spheroids to be formed. Such a mold is shown in the picture below. Working with such a form like this requires a large number of cells: with a quantity of 1000 cells per microwell, 2.8E+07 cells taken from two or three T-225 flasks are required (Figure 18).

**Figure 18.**
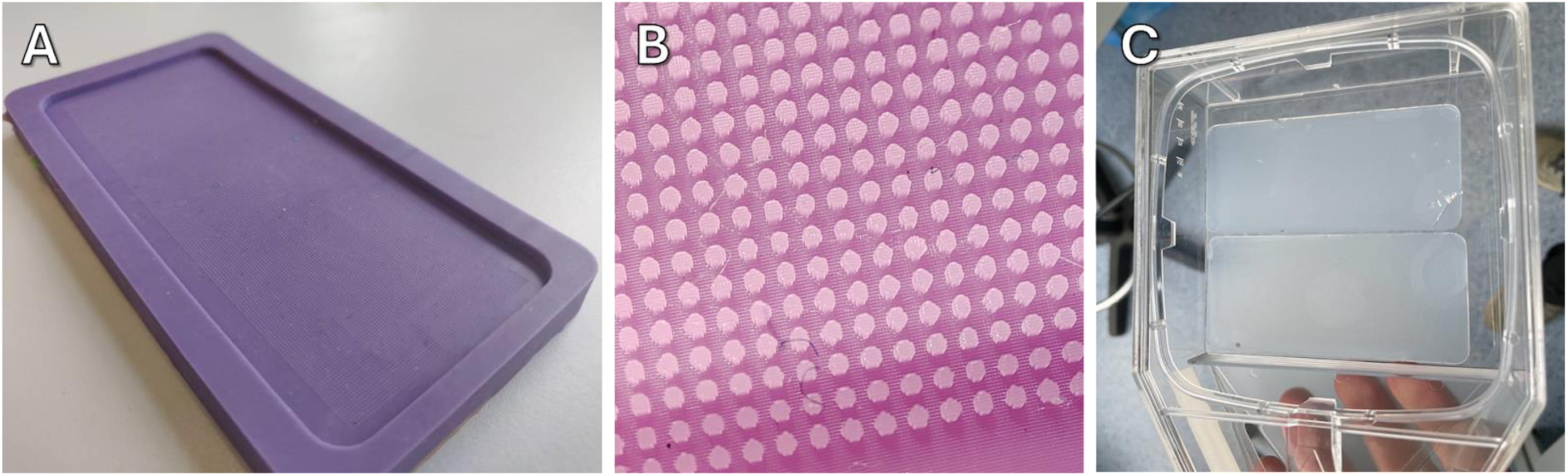
Photograph (A) and macrophotograph (B) of a silicone mold with 14282 cylindrical micropins and agarose molds made with it placed in flask with re-closable lid.

### Stamp for 96-well plates and cytotoxicity study

The 96-well plate is the workhorse of cell biology, where a wide variety of tests and assays can be performed on cell cultures, while remaining relatively affordable and not requiring the use of robots, as is sometimes necessary for plates with a large amount of wells [35]. We have tried to show that it is quite easy to adapt the wealth of available screening assays to spheroids created by our approach as well.

Agarose thickness is not a problem here as with dish stamp, but the challenge is quantity. It would not be so difficult to make a silicone mold in which agarose would be poured and after solidification would be transferred to the wells of the plate, but to do it 96 times for each plate would be quite time-consuming and tedious procedure. Consequently, we went back to the stamp approach. We tried to make the process as fast and convenient as possible, since we are talking about screening studies, which are often quite large and even the small-time costs start to add up. In addition, the whole approach is designed for the use of a multichannel pipette, which allows efficient dosing of liquids including pouring molten agarose.

Making a stamp for a 96-well plate also took several iterations and we finally settled on the following design (figure 19). Like the dish stamp, this one is printed directly on the printer’s bed, without supports. In the upper part there is a flat plane for labels (1), in the main part (2) there are holes for better handling (3). Centering rings (4) allow to locate the stamp in the wells of the plate when the rods (5) are immersed in on the bottom of the rods there are micropins (6), which form microwells. The label (7) indicates the ventilation channels inside the stamp, they have no effect on the microwell manufacturing process itself, but they simplify the stamp printing process by reducing the suction effect from the flat surface. In addition to printed stamps, we decided to try stamps made of silicone to compare the usability (Figure B), which are completely identical in the lower part of the design, but due to the difficulties of working with silicone do not have a handle on top.

**Figure 19.**
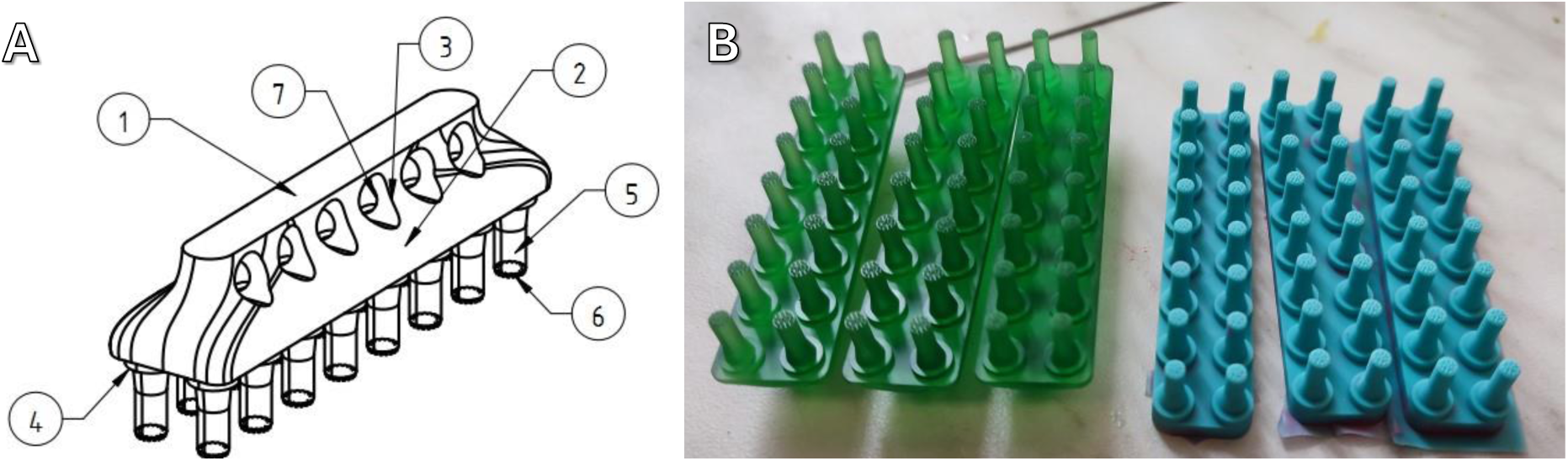
A - sketch of a stamp for a 96-well plate. B - photos of stamps printed on MSLA printer (left) and silicone stamps similar to them in design of the working part (right).

What you should pay attention to at once - each stamp is designed for two columns - 16 wells of a 96-well plate. Larger stamps are rather inconvenient both for printing and for handling, as well as they are extremely difficult to remove due to adhesion of the stamp in agarose in each well.

Accordingly, agarose is poured into two columns of wells with a pipette, and after that the stamp is placed there (Figure 20A). As we mentioned above, the most homogeneous distribution of spheroids was obtained in cone-shaped and cylindrical microwells, but after a number of pilot experiments with 96-well plates, we noticed that cylinders are much more likely to cause damage to the agarose mold, probably due to the larger surface and, as a consequence, greater adhesion to agarose, respectively, we finally settled on the cone shape of microwells (Figure 20B), which allows to achieve the formation of 1 spheroid per microwell (Figure 20C).

**Figure 20.**
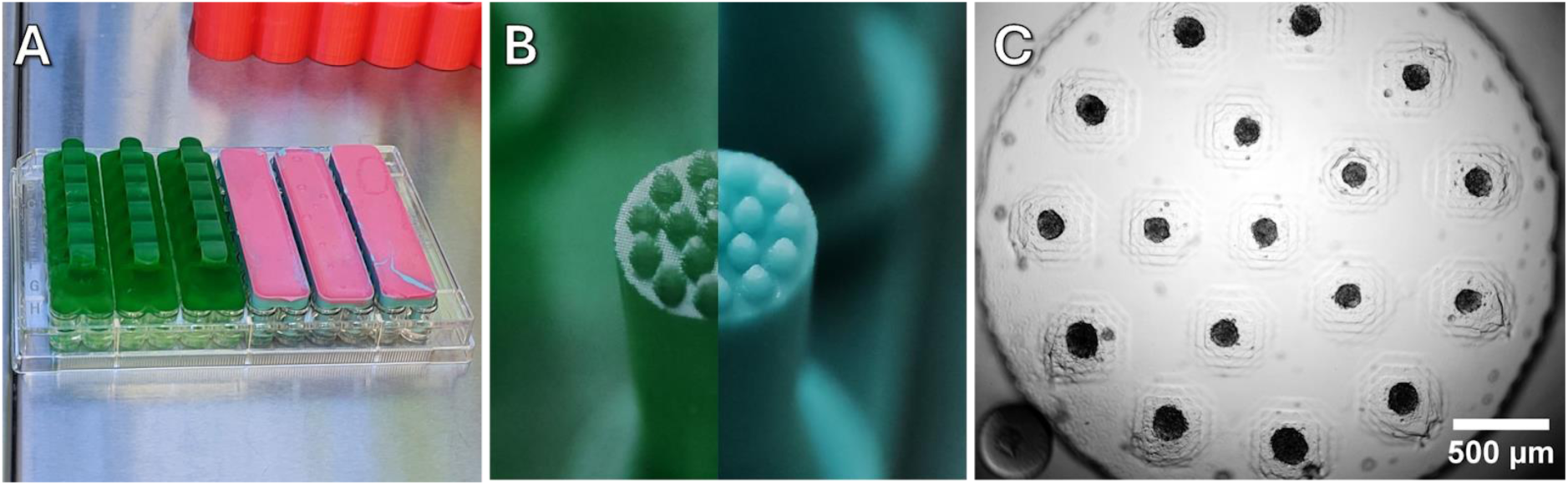
A - printed (left) and silicone (right) stamps installed in the plate. B - working part of the printed and silicone (left and right, respectively) stamps for 96-well plates for the production of cone microwells. C - spheroids formed in microwells

From several variants of configuration of microwells arrangement in the mold we decided on 34 micropins located in concentric circles. The area of the mold itself is limited by the size of the field of view in the microscope - we wanted the whole mold to fit into it, this greatly simplifies screening observation, and the location and number of pins is somewhat arbitrary. Microwells should not be placed too close (as mentioned in the section “Diameters, depth and height capabilities”), as for the rest there can be many variants of their configuration, which we have not found significant pros and cons.

As in the case of glass bottom dishes, the volume of agarose in the well is important here. The volume of gel must be sufficient: in a small volume, microwells are not formed; in a large volume, the adhesion of agarose to the stamp is too strong and the agarose mold can tear when stamp removed. For our stamps and the plates used in this work, the optimal volume was 50±10 µl. If there is too much agarose, it will adhere more strongly to the stamp and when the stamp is removed, the microwell will tear or be completely removed from the plate. If there is too little agarose, then, logically, the stamp does not reach it at all. However, even with the optimal amount of agarose, defects may occur during the production of microwells. The most common ones are shown in the figure below. Sometimes an air bubble is trapped under the agarose and does not come out of it (Figure 21A), such defects do not affect the formation of spheroids but can complicate computer quantification. If the bubble gets trapped in the gap between the agarose and the stamp (Figure 21B and 21C), some of the wells may be damaged, resulting in spheroids not forming there.

**Figure 21.**
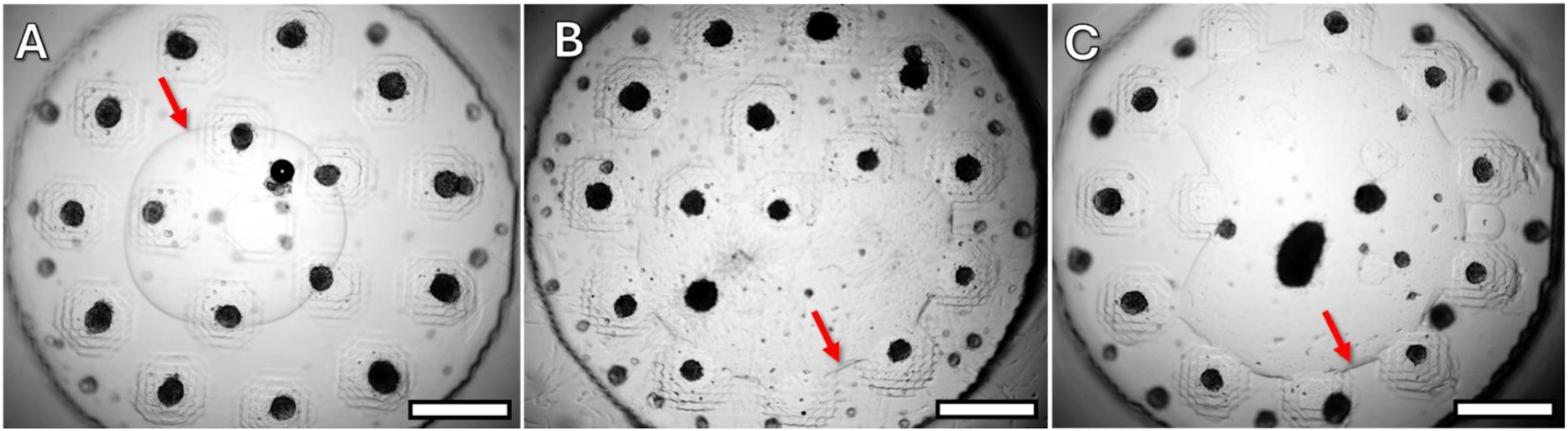
Various defects formed during the manufacture of stamps. A - bubble located under the agarose mold. B - absence of cells in the mold. C - bubble in the area of the mold where microwells are located. The arrows indicate bubbles in different parts of the mold. Scale bar - 300 μm.

Stamps printed on MSLA printer and stamps made of silicone with identical shapes were created. Immediately at microscopic inspection it was found out that the defects described above occur in molds made with silicone stamps very often, somewhere in one third of wells. Probably the adhesion of agarose to silicone is higher than to photopolymer. Nevertheless, the spheroids were photographed, and their size and shape were measured (graphs in the figure 22).

**Figure 22.**
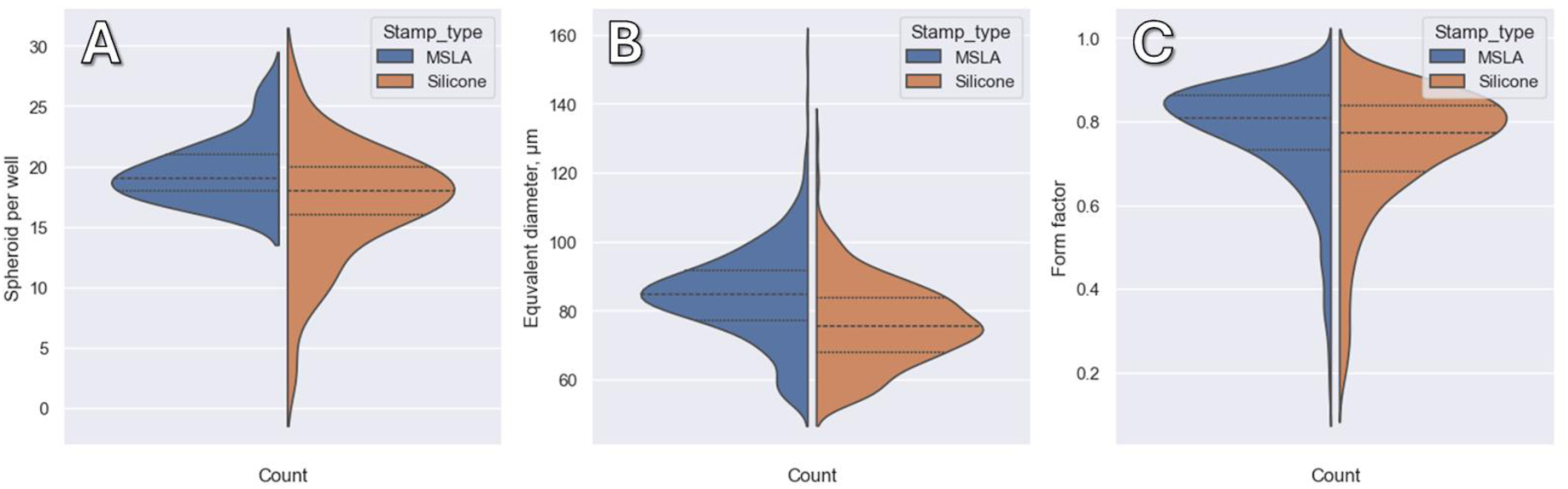
Violin-plot showing the number of spheroids per cell (A), size distribution (B) and form factors (C) of spheroids in microwells made with printed stamps and their silicone counterparts

The average size of the spheroids made by MSLA stamps was 84±12 μm and in the ones made with silicone 76±12 μm, the average form factor was 0.77 and 0.76, respectively. Overall, when looking only at the numerical values MSLA and silicone stamps are very close to each other. Although if we evaluate the distribution (Figure 22), it is obvious that the number of spheroids in molds made with silicone stamps is much less reliable. In addition, making silicone stamps is a noticeably more tedious process, hence in the future we will use exclusively printed ones.

Of course, different numbers of cell spheroids is required for different tasks. And, for example, to study their metabolic activity during cytotoxicity studies of various drugs, the more spheroids the greater the change in the analytical signal in each well of the plate will be achieved. Accordingly, we made a series of stamps with different numbers of micropins: 7, 17, 25, 34, 49 and 57. The micropins are distributed differently, 7, 17 and 34 as concentric circles, 25 and 49 as a square grid. An image of the spheroids formed in the respective microwells is shown in the figure 23.

**Figure 23.**
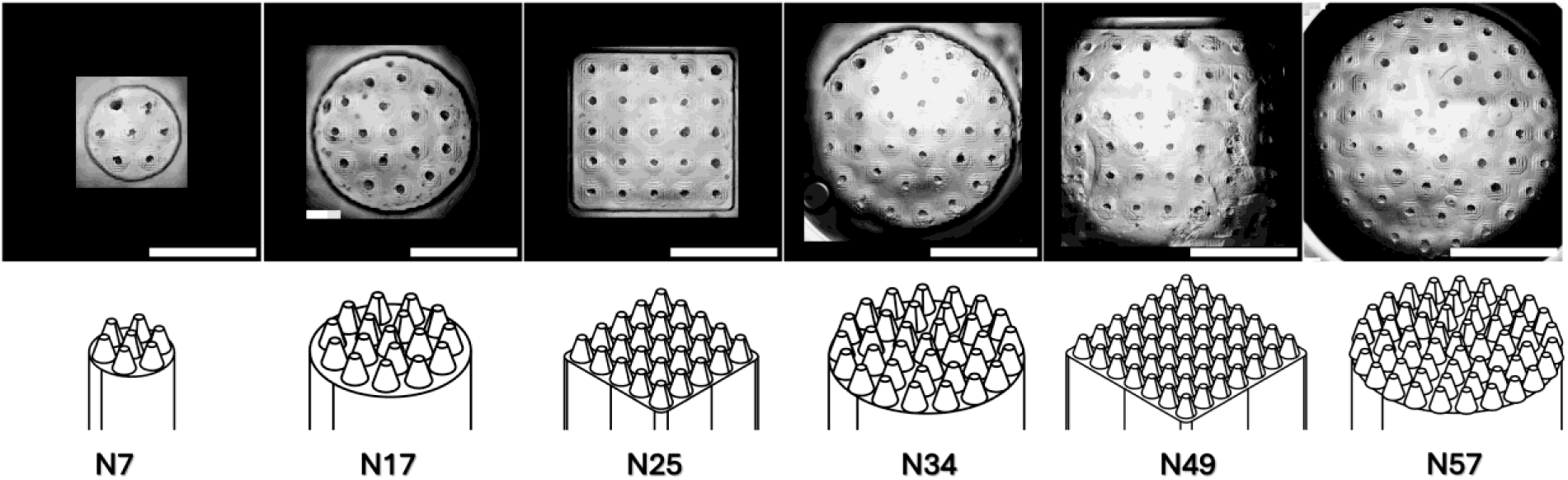
Spheroids from HEK-293t culture cells grown in cells of a 96-well plate with different numbers of microwells. Images were obtained by stitching individual frames together. Scale bar - 1000 µm.

The necessity for a large stamp area is a consequence of the requirement for a large number of microwells, as previously discussed. So, it is not possible to place micropins too close to each other. Nevertheless, a larger stamp area enhances the adhesion of the gel to the stamp, which results in the rupture of the gel layer in three-quarters of wells when using stamps with 49 and 57 micropins. The figure above illustrates the rare instances where the process was successful. Accordingly, the stamp with 34 microwells is the largest one that can be used effectively, and further work was carried out using such stamps. Its only disadvantage is that the whole image does not fit in the field of view of the microscope, however, it is not so essential for a high-throughput metabolic assay.

As mentioned above, an important task is also the determination of the optimal number of cells for the production of cell spheroids. In 96-well plates, this task is technically easier and more convenient than in the large molds described above. For this purpose, it is enough to seed the cells in a series of wells, diluting them twice in each subsequent well. Below is a figure 24 showing the result of such an experiment.

**Figure 24.**
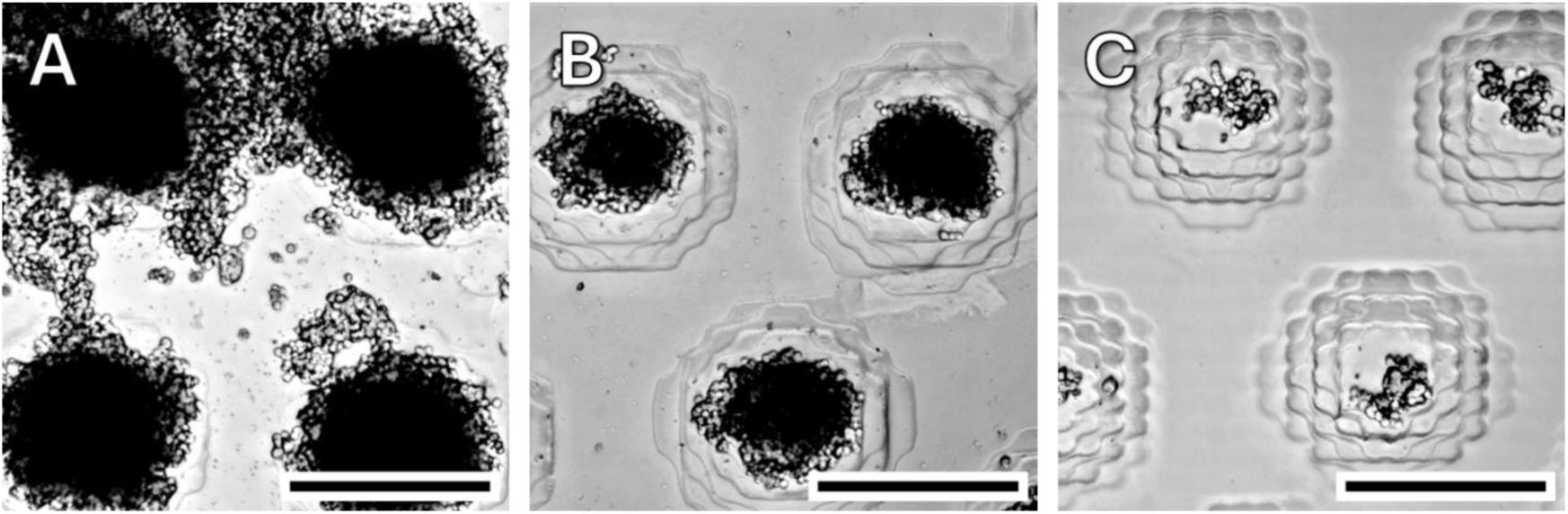
Formation of spheroids from different numbers of HeLa cells in microwells in a 96-well plate. A - 4000 cells per microwell, B - 1000 cells, C - 100 cells.

### Cell viability testing

To demonstrate the applicability of agarose microwell plates made by our method, we chose the MTT and a resazurin assays, both well-known and incredibly popular techniques for screening the effect of various substances on cell viability. These analyses are based on common principles and are not without their drawbacks. However, if done correctly, they are surprisingly cheap, simple and relatively reliable [36].

The MTT and the resazurin assays are analyses based on the assessment of the level of cellular respiration [37]. In both cases, cellular oxidoreductases metabolize the respective substances into products with different optical characteristics. The reaction products are colored variously, and resafurin (resazurin reduction product) in addition fluorescents in a different range. Measurement of optical absorbance (in the case of the MTT assay) or fluorescence (in the case of the resazurin) provides an indication of cell activity and viability.

However, their essential difference lies in the methodology of the experiment. The MTT reduction product is insoluble in water and to measure its concentration (in a typical setup) it is necessary to remove the medium, add the medium containing MTT to the wells, after incubation remove the medium and replace it with a solubilization solution, usually DMSO, detergent solution or acidified alcohol and only then measure the optical density of the liquid. These manipulations are not difficult to carry out with a multichannel pipette, but when there is a gel insert in the plate well, everything becomes much more complicated. It is easy to damage the insert and when removing the liquid, a piece of gel will get stuck in the tip, resulting in an incorrectly sampled volume.

The resazurin assay, on the other hand, requires only the addition of a resazurin solution and, after a certain amount of time, a measurement of optical density or fluorescence intensity.

Before conducting more serious experiments it was decided to compare the applicability of both methods. For this purpose, HEK-293t cell spheroids were formed in 96- well plates and the corresponding viability assessment method was applied to each plate. Of course, the MTT method was much more labor intensive and the extraction of the MTT derivative from the spheroids with DMSO took considerable time (and there is no guarantee that it was extracted completely). The numerical results are shown in Figure 25.

**Figure 25.**
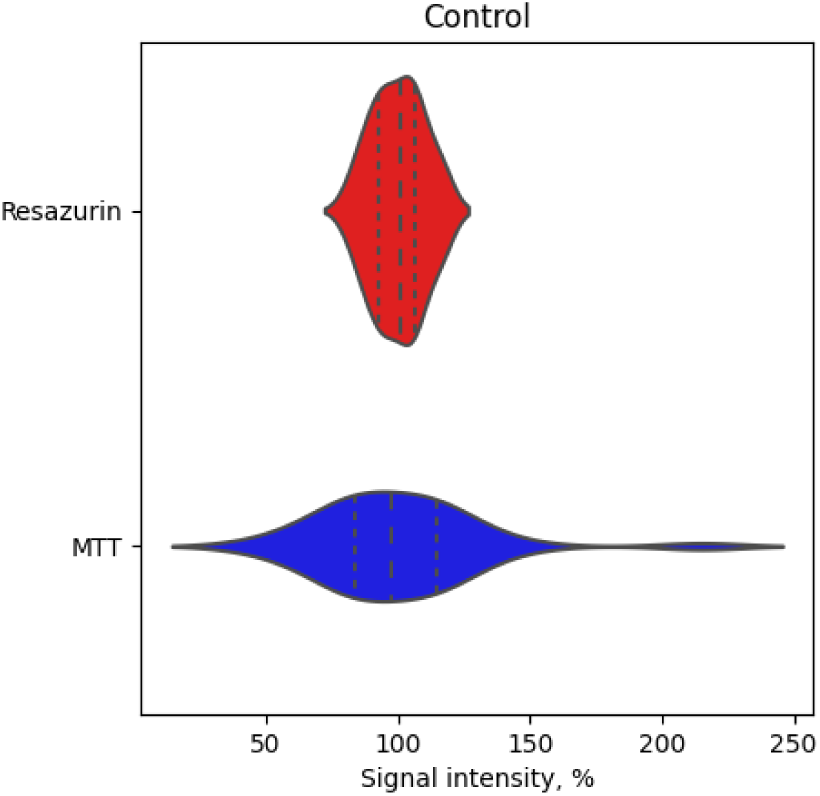
Violin-plot showing the intensity distribution of the corresponding signals for spheroids formed from HEK-293t to which no additional substances were added.

Even visually, one can see how much less reproducible the MTT test is compared to the resazurin assay when applied to spheroids in microwells. The standard deviation for the resazurin test is about 9%, whereas for the MTT test it is about 30% which makes it practically pointless. Here it should be added that besides MTT there are other similar tests (XTT, MTS and so on) that do not require so many manipulations, but MTT has been and remains the most popular one, probably due to its low cost.

Similar results were obtained when these tests were performed on spheroids grown from cells to which magnetic nanoparticles with an iron core-carbon shell structure were added. We knew from earlier studies that these nanoparticles do not significantly affect the metabolic activity of cells [38], and now we have tested this on spheroids, with the same outcome. However, a comparison of the data obtained using the MTT and resazurin assays revealed that the MTT assays performed on spheroids in agarose microwells is considerably less reliable than the resazurin test, as was anticipated. The figure 26 below shows the results of metabolic tests, where we can see a significant difference in confidence intervals for MTT and resazurin, as well as illustrative images of spheroids formed from normal cells and from cells saturated with magnetic nanoparticles.

**Figure 26.**
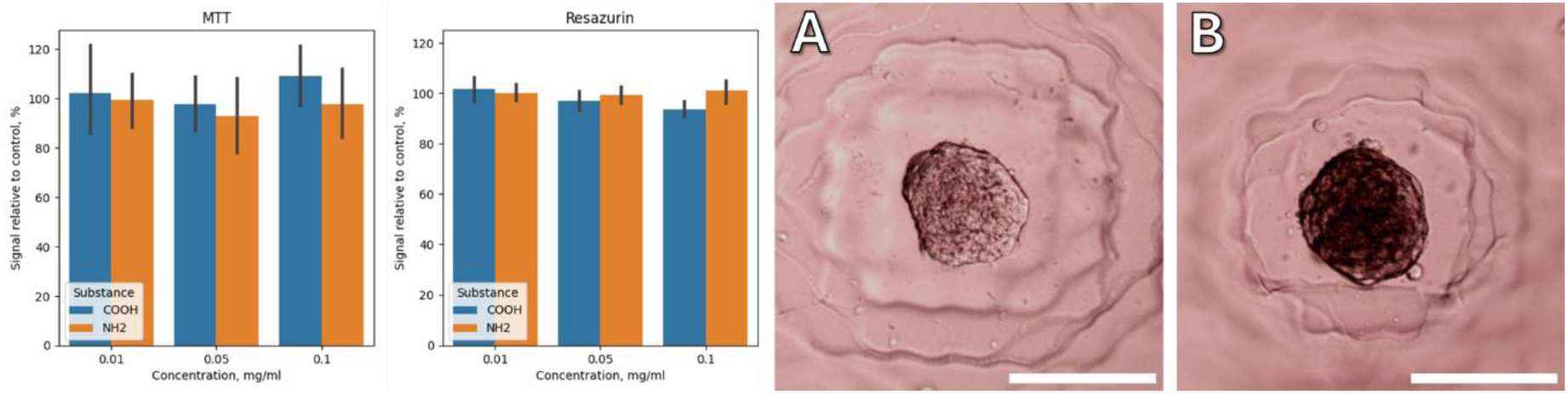
MTT and resazurin assay results performed on ACS52 telo spheroids incubated with Fe@C magnetic nanoparticles. A and B – images of spheroids without (A) and with (B) nanoparticles. Scale bar – 150 um.

Despite the fact that there is no need to use DMSO to perform the resazurin test, it is still a very common solvent in biology. Many substances, including cytotoxic and cytostatic drugs, are dissolved in DMSO when added to cell culture. To estimate its effect, we cultured spheroids in a medium with different amounts of DMSO and assessed their respiratory activity using the resazurin test. (Figure 27) .

**Figure 27.**
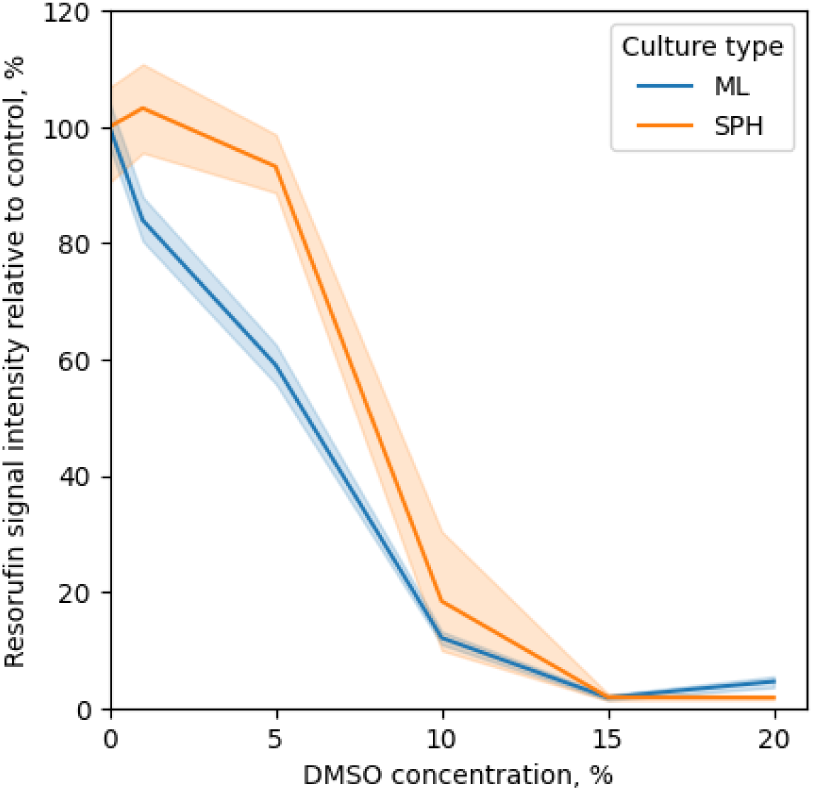
Effect of varying concentrations of DMSO in media on the metabolic activity of cells (HEK-293t) in spheroids (SPH) and in normal adherent cell culture (ML).

There are publications describing that cells in a spheroid are often relatively more resistant to various influences than when they grow as a monolayer on plastic, due to the hindered diffusion of the toxic substance inside [39]. The data obtained on spheroids formed from HEK-293t confirm this; a decrease in the viability of a monolayer culture is noticeable already in the presence of 2% DMSO in the medium, while for spheroids a significant effect is recorded only in the presence of 5-7%. Naturally, such high concentrations of this solvent are rarely used (apart from the cell freezing procedure, where other goals are pursued). Nevertheless, these results support the possibility of using the resazurin test to assess the viability of spheroids in microwells.

Next, we conducted a screening study, taking a set of substances used in cell biology and medicine as cytostatic drugs. The set of substances included the mTOR inhibitor Everolimus, the intercalating drug Doxorubicin, the alkylating agent Oxaliplatin, the microtubule stabilizers Paclitaxel and Docetaxel, and the topoisomerase inhibitor Camptothecin. All drugs were added to the medium at a concentration of 20 μM, which significantly exceeds the IC50 determined for these substances for conventional cell cultures. However, in this case, our goal was to see the effect, and not to conduct a systematic study, which is outside the scope of this particular work.

As a positive control, we took a 30% DMSO solution in the nutrient medium, which reliably kills cells, including those in spheroids, as demonstrated above. The data obtained during the resazurin test is shown below, along with images of spheroids after 72 hours of incubation in the corresponding substances.

The obtained data shows (Figure 28) that different drugs have quite different effects on the metabolic activity of cells in the spheroid, which is not unexpected in general. Everolimus and Doxorubicin have proven to be the most toxic in this series.

**Figure 28.**
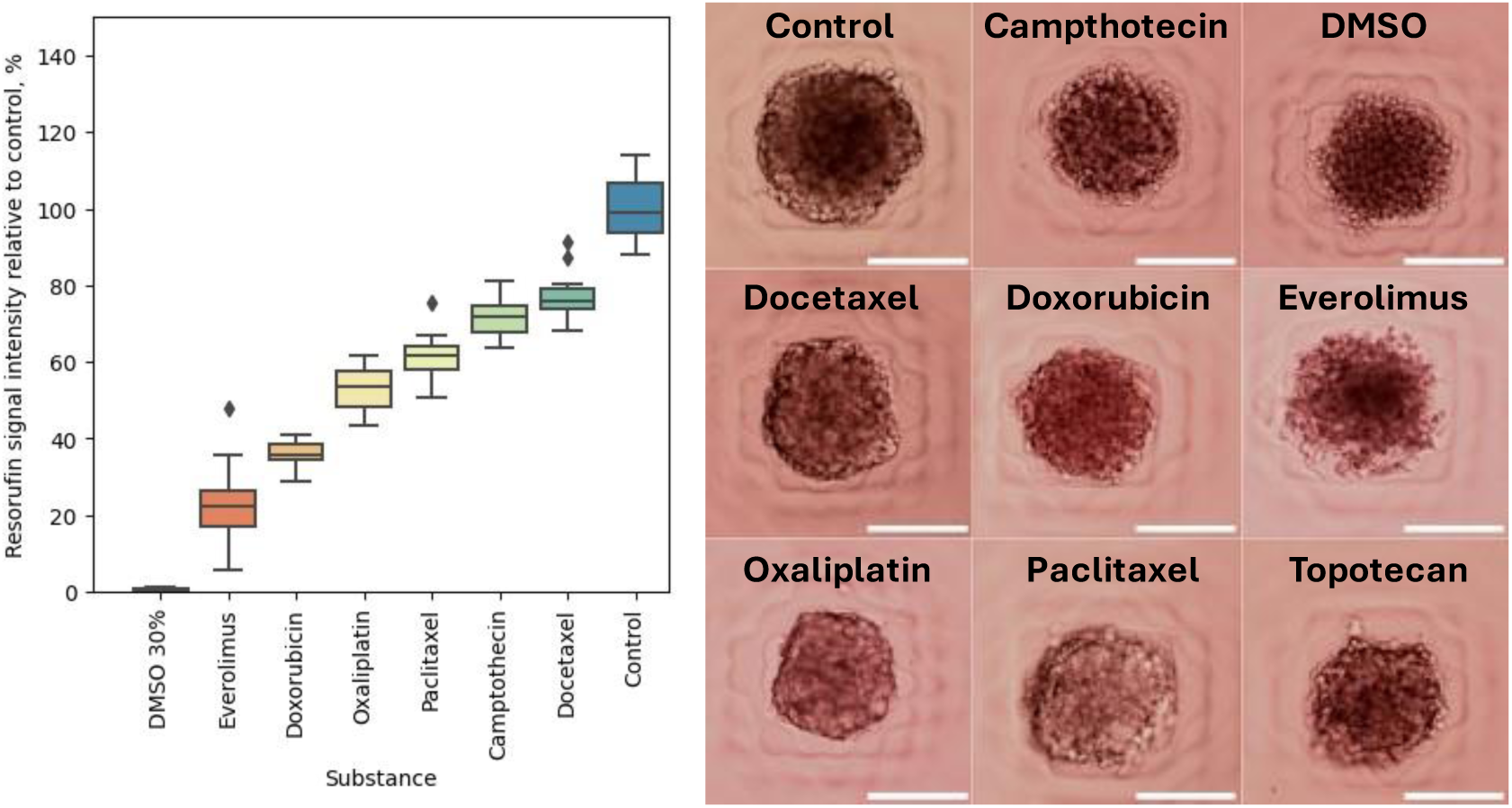
Effect of various cytostatic drugs (each was added at a concentration of 50 μM) on the metabolism of spheroids formed from the HEC-293t culture (left) and images of spheroids obtained after 72 hours of incubation in a medium containing the corresponding drugs (righs). Scale bar - 150 μm.

For a more in-depth study, we chose doxorubicin as a common substance often used as a positive control in cytotoxicity studies on cell cultures. HEK-293t cells growing as a monolayer or as a spheroid in gel microwells were incubated with different concentrations of doxorubicin in the medium. The obtained data was processed using the Hill equation, which is usually used to determine the dose-response relationship in various biological experiments, and in cytotoxic studies allows to determine the IC50 value. The obtained dependency graphs fitted by the Hill curve are shown in the figure 29.

**Figure 29.**
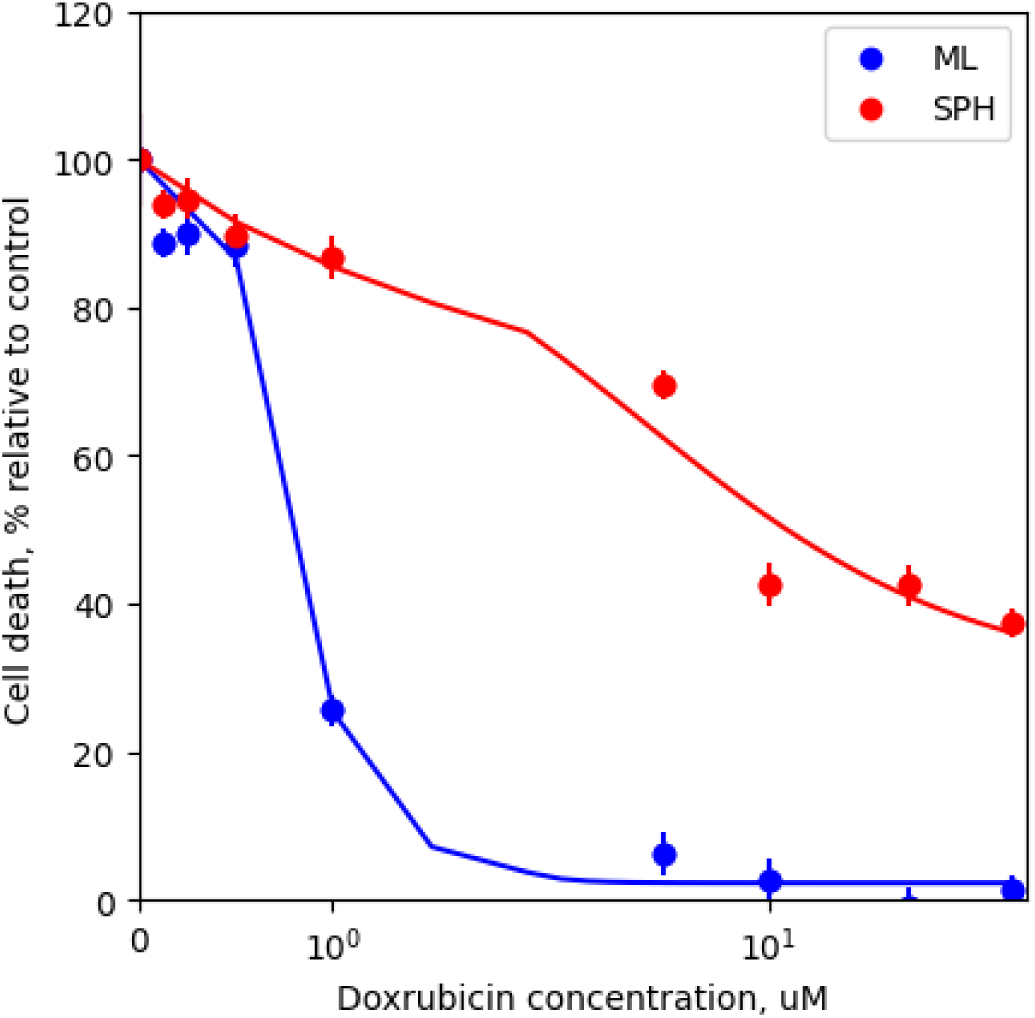
Response of HEK-293t cells (adherent culture and spheroids in microwells) to different concentrations of doxorubicin, determined using the resazurin assay

It should be noted that for spheroids the dependence has a different form from that usually obtained on adhesive cultures. Nevertheless, IC50 was determined, was 0.77 μM for adhesive culture and 4.3 μM for spheroids. And, as we wanted to show, in general, the process for spheroids and conventional culture was not different, except for the process of spheroid production. Thus, we demonstrate that it is possible to perform routine cell biology tasks on spheroids using our approach without much difficulty or expense.

### Image processing

At the same time, the effect of substances on spheroids is noticeable not only in the metabolic test, but also visually. In the control sample, the formation of a necrotic core is visible, in the sample with DMSO and Everolimus, disaggregation of the spheroid into individual cells is observed. Methods for processing images of spheroids as a way to assess their vital activity are also described in the literature [33,40,41], and gel microwells are ideal for such application. The spheroids are fixed in the microwell and will not move anywhere when manipulating the plate, which allows you to observe them for several days.

To analyze the morphology of spheroids we used the open source and free software CellProfiler. Of course, nowadays there is quite a lot of software for processing microscopic images and, image processing in the era of neural networks has reached new heights, but here we concentrated our efforts on a simple tool available to any biologist, including those without deep knowledge in programming. Despite its simplicity, CellProfiler has quite a few tools for evaluating different properties of objects in an image. We used the built-in modules of the software that allow us to estimate the size and shape of objects (MeasureObjectSizeShape), the heterogeneity of its texture at the level of individual pixels (MeasureTexture) and on a larger scale (MeasureGranularity), as well as the distribution of colour intensity on the object (MeasureObjectIntensityDistribution). Each of the modules generates an array of metrics and we used ML techniques to highlight the most significant factors, which may not be obvious to the human eye at first.

Preliminary tests were performed with synthetic images, where it was shown that the use of our pipeline does produce the expected results. It was shown that it is possible to detect the change of radial distribution in the image, as well as local inhomogeneity in contrast (granularity). Images and results of their processing are given in the supplementary materials.

Further, the real images of spheroids were analyzed. Of course, the natural heterogeneity of biological samples makes its own adjustments, and the results are not as obvious as when analyzing synthetic data. One of the key results: there is no single metric for comparing the effect of different drugs on cell spheroids. This has already been partly discussed in the paper [42], where the authors drew attention to the marked differences in the effect of drugs, some leading to a change in size, others affecting morphology, others causing separation of the spheroid into isolated cells.

Tables with highlighted metrics for each individual substance are given in the supporting materials, individual examples are given below. For instance, the effect of topotecan, its effect is to slow the growth of spheroids and slightly reduce their size, whereas in the control, spheroids grow without restriction, reaching their maximum size determined by nutrient diffusion inside (Figure 30).

**Figure 30.**
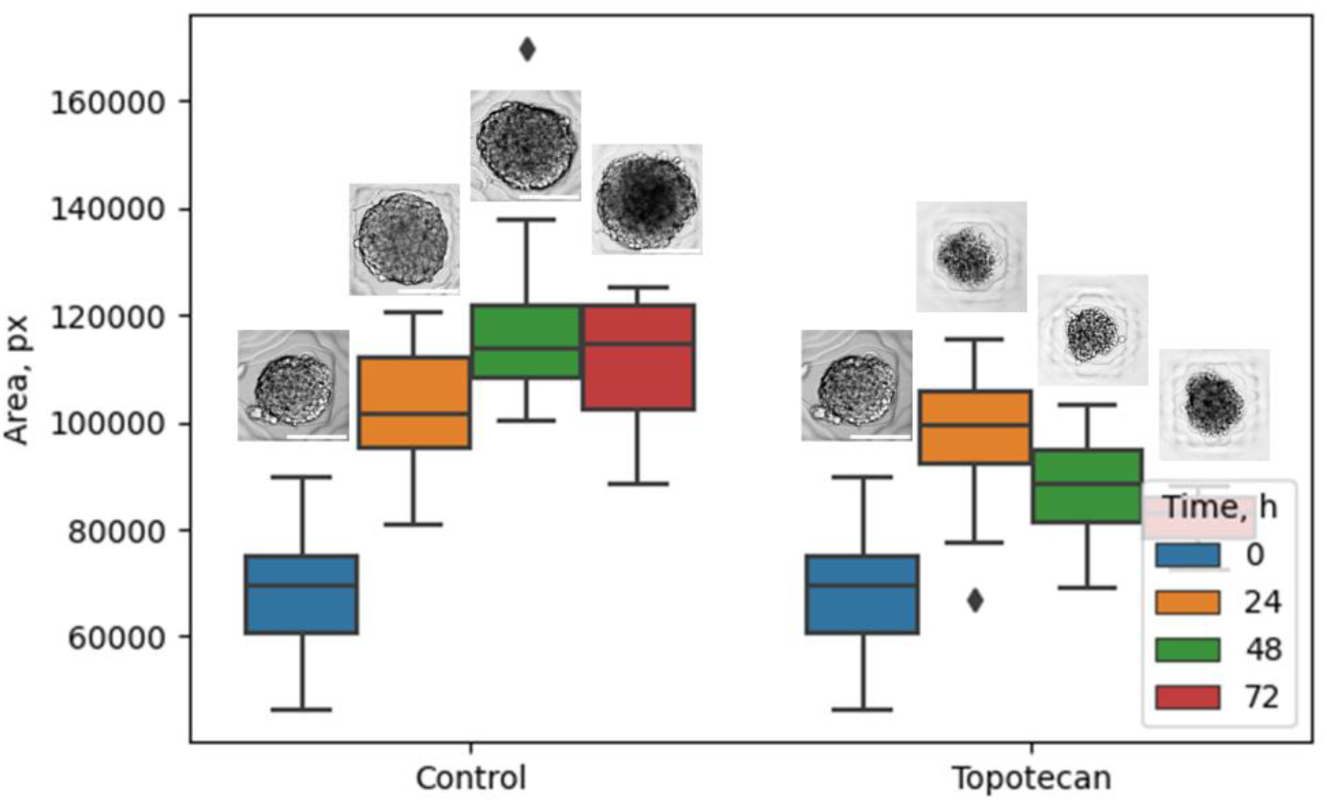
Change in the size of spheroids formed from HEK-293t cells exposed to topotecan over time. Representative images of these spheroids are shown for illustration.

In spheroids under the influence of e.g. doxorubicin, cell division and metabolic activity are suppressed and hence no darkening in the center is formed (Figure 31).

**Figure 31.**
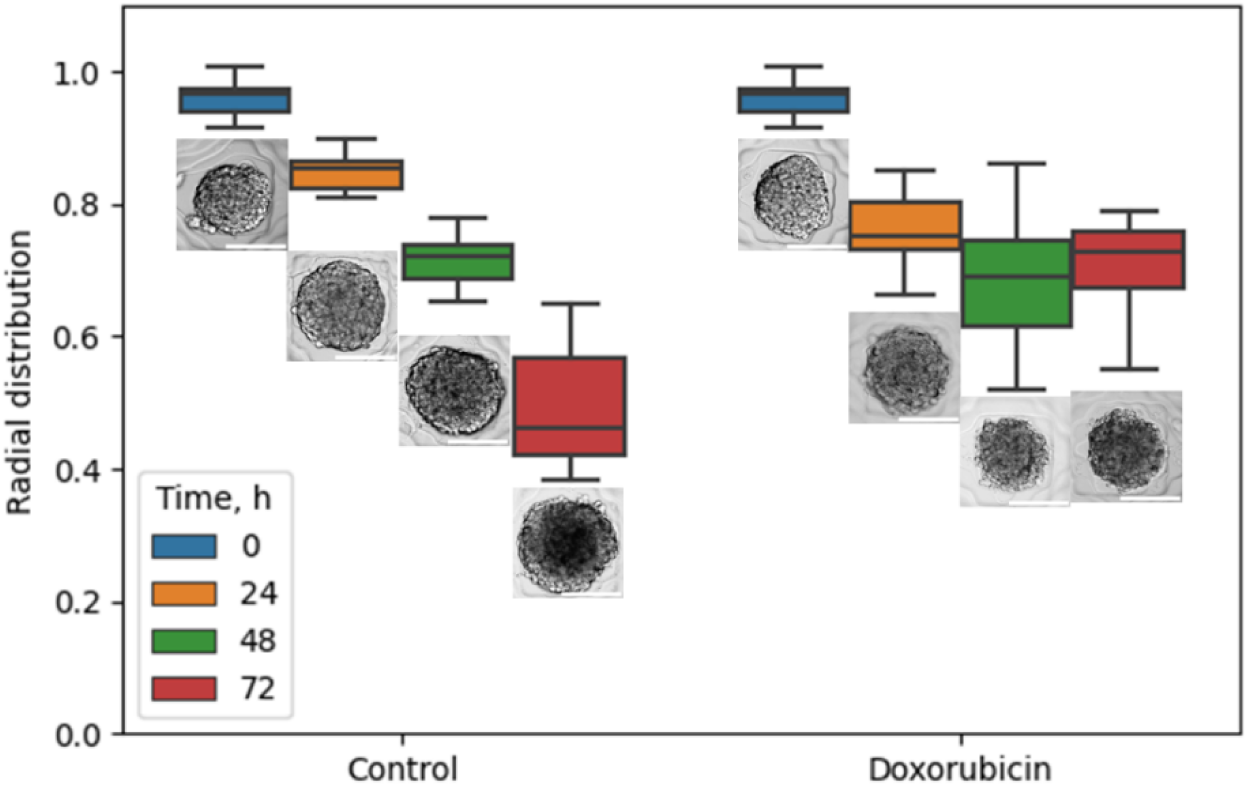
Changes in radial brightness distribution in spheroids formed from HEK- 293t cells exposed to doxorubicin over time. Representative images of these spheroids are shown for illustration.

When a spheroid is disaggregated, there is an increase in its optical heterogeneity (measured using the MeasureTexture and MeasureGranularity functions). The difference between control spheroids and spheroids affected by everolimus is not so large, but still noticeable (Figure 34).

**Figure 34.**
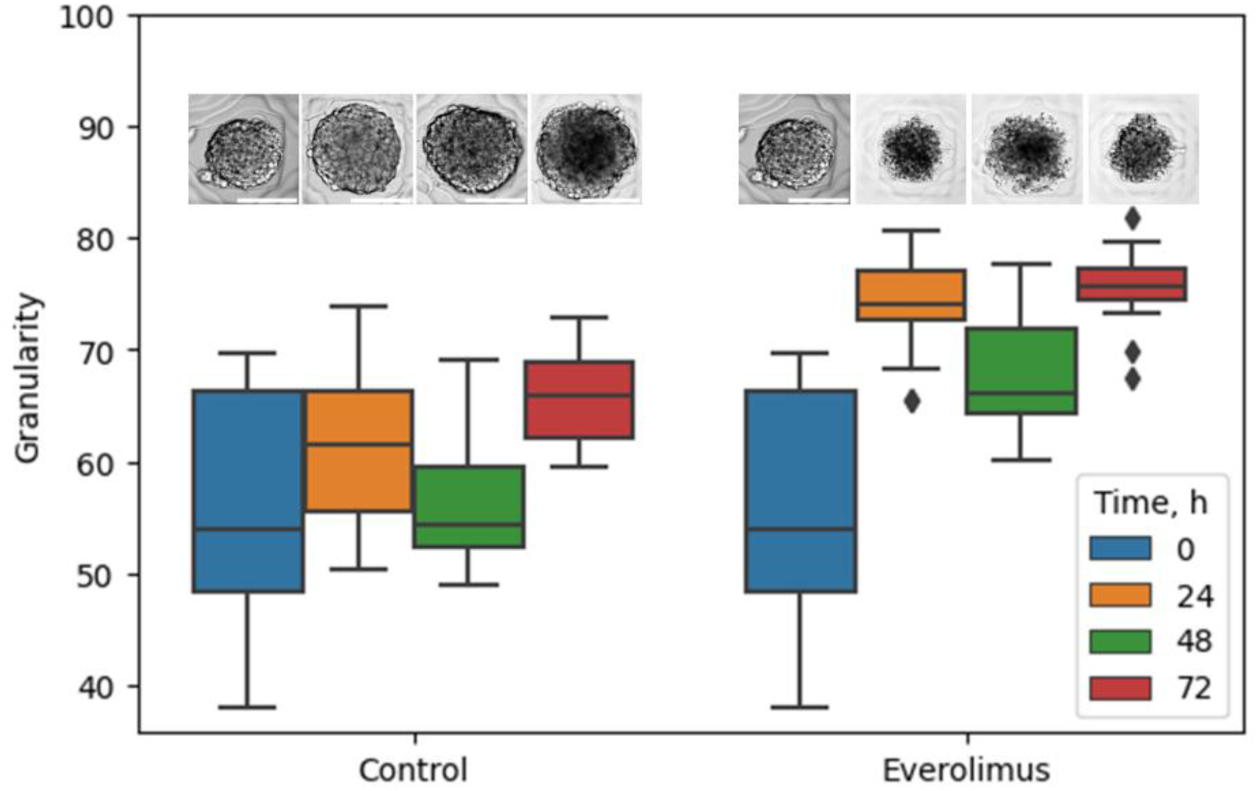
Change in the heterogeneity of brightness (granularity) in spheroids formed from HEK-293t cells exposed to everolimus over time. Representative images of these spheroids are shown for illustration.

Finally, we have shown that spheroids grown in microwells which were made according to our proposed method, can be studied not only by metabolic tests, but also by analyzing their images. And, of course, this can be carried out in convenient and familiar for biologists 96-well plates.

## Repeatability

Scientists in other laboratories (not affiliated with the authors of the study), having received stamps, silicone molds and instructions, were able to reproduce the process of stamp printing and spheroids manufacturing (Figure 35). They used other cell cultures, laboratory plastic from other manufacturers, as well as agar and agarose of other brands. Instructions as well as 3d models are available on the project page on GitHub and in the supplement materials (section S5).

**Figure 35.**
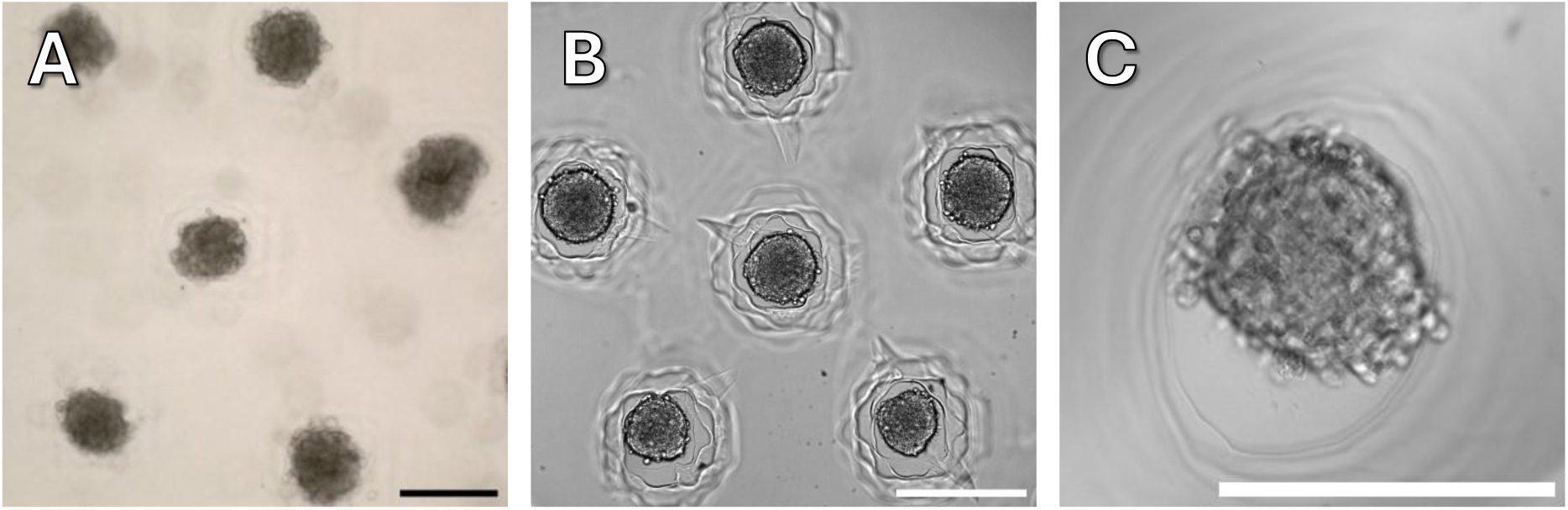
A – cell spheroids from HT-29 cell culture made in Pacific Institute of Bioorganic Chemistry RAS, B – spheroids from HCT116, made in Engelhardt Institute of Molecular Biology, C – spheroid from human dermal fibroblasts made in FRC ‘Fundamentals for Biotechnology’ RAS. Scale bar 300 µm.

In general, scientists from independent laboratories have been able to replicate the spheroid fabrication process described in this article without difficulty. Necessary clarifications and additions that arose during the discussions were added to the protocols. However, in one case, our colleague encountered the problem mentioned above, the size of the culture plate is not standard. He used our stamps for 96-well plates with costar 3599 plates (Corning, USA) and found that they are a fraction of a millimeter shallower, which causes the stamp to pierce the gel to the plastic. This leads to migration of cells under the agarose layer instead of forming a spheroid. After the necessary correction of the model and printing of a new stamp, the experiment was completed successfully.

## Pricing

Speaking about the possibility of adapting our approach, its cost should be mentioned. In our opinion, it turned out to be quite affordable by the standards of any scientific organization.

The prices given below (table 1) are somewhat conditional, as the cost of different items may vary significantly depending on regional exchange rates, taxes and duties. Also, the prices do not take into account the cost of electricity and other household needs, as well as payment to the printer operator. However, we hope that this will give an idea at least to an order of magnitude. The main things you need to have (in addition to standard equipment and supplies for working with cell cultures) are a 3d printer, photopolymer resin and two-component silicone. For making silicone molds it is also necessary to have a vacuum chamber with an appropriate vacuum pump.

**Table 1.** Cost of the basic materials and equipment required for making stamps and molds.

| Equipment and Materials | Quantity | Price, USD |
| --- | --- | --- |
| MSLA 3d printer | 1 pcs | 400 |
| Vacuum chamber with vacuum pump | 1 pcs | 450 |
| MSLA resin | 1 liter | 50 |
| Two component silicone | 1 liter | 50 |

If only direct costs are taken into account, the cost of stamps and molds is determined by their volume. The stamp designs presented in this article were partially optimized in terms of reducing the cost of photopolymer and silicone, but this was not the main goal; the price, however, was already relatively low (Table 2).

**Table 1.** Cost of the materials required for making stamps and molds per stamp/mold.

| Pices | MSLA resin volume/ Silicone volume, ml | Cost, USD |
| --- | --- | --- |
| Petri dish stamp | 7/none | 0.4 |
| 6 well mold | 6/5 | 0.3/0.3 |
| T-150 flask mold | 44/39 | 2.5/2.1 |
| 96 well stamp | 10/none | 0.6 |

As you can see above, the cost of one mold is quite low, especially when compared to the commercial options (a set of Microtissue molds costs a couple hundred dollars) described above in the article. At the same time, printed stamps and silicone molds can be used many times. The master molds from which silicone replicas are made, however, gradually degrade and you can make 4-5 casts from one before the quality starts to drop and the micropins are destroyed when removing the silicone mold.

In addition to the relatively low price, the DIY approach allows you to customize stamps and molds to suit your specific needs. It takes two to three hours to print several stamps (depending on the specific printer), it takes about a day to make a silicone mold from a printed stamp (time for final polymerization of silicone).

## Conclusion

We have developed approaches to reliable and reproducible agarose microwell fabrication with rich customization possibilities in order to democratize, simplify and make this process accessible to every laboratory. Using MSLA 3d printing, we have produced a series of stamps as well as master molds for making silicone molds that allow us to make agarose microwells at a wide variety of: from a few cell spheroids in a well of a 96-well plate to tens of thousands in T-150 culture flasks.

We have shown how the fundamental limitations of this 3d printing method determine the size and shape of microwells and, using confocal microscopy, demonstrated how this affects the process of spheroid formation. We compared our approach to the mass production of spheroids with its commercial counterpart and showed that, despite the limitations of our technique, spheroids are generally similar. Finally, we have shown the prospects of growing spheroids in 96-well plates. This method opens up the possibility of high throughput screening of various substances on spheroids, using standard approaches utilized in the cell culture laboratory, by modifying the routine resazurin assay on cellular spheroids with a number of model objects.

## CRediT author statement

Minin, A.: Conceptualization, Methodology, Software, Validation, Investigation, Writing - Original Draft, Visualization. Semerikova, T.: Investigation, Resources. Belousova A.V. - Investigation. Karavashkova, O.: Investigation; Pozdina, V.: Investigation. Tomilina M.: Investigation, Zubarev, I Supervision, Funding acquisition.

## Acknowledgments

The reported study was funded by the Russian Science Foundation Grant #22-74- 10041. Confocal microscopic examination was performed using the equipment of the Shared Research Center of Scientific Equipment SRC IIP Ural Branch of RAS.

The authors are enormously grateful to Vladimir Tatarsky for providing resazurin as well as cytotoxic drugs: paclitaxel, everolimus, doxorubicin, topotecan, docetaxel, camptothecin and oxaliplatin.

For testing our methodology and proving its reproducibility, we are grateful to Anatoliy Zubritskiy from FRC “Fundamentals for Biotechnology” RAS (Moscow, Russia), Kuzmich Alexandra and Yurchenko Ekaterina from Pacific Institute of Bioorganic Chemistry RAS (Vladivostok, Russia), Alexandra Dalina from Engelhardt Institute of Molecular Biology RAS (Moscow, Russia) and Saida Karshieva from National University of Science and Technology MISIS (Moscow, Russia)

