## Supplement for "MSLASpheroidStamp: 3d cell spheroids for everyone"

### S1. Print settings

Print settings used in the work, matched to the printer and the photopolymer we used. Once again we remind you that for another printer and another resin the settings **will be different**! And even for the same printer and the same resin, but under different conditions (e.g. room temperature can play a big role). It is recommended to use a test object such as [Cone of Calibration] to adjust shutter speeds and other printing parameters.

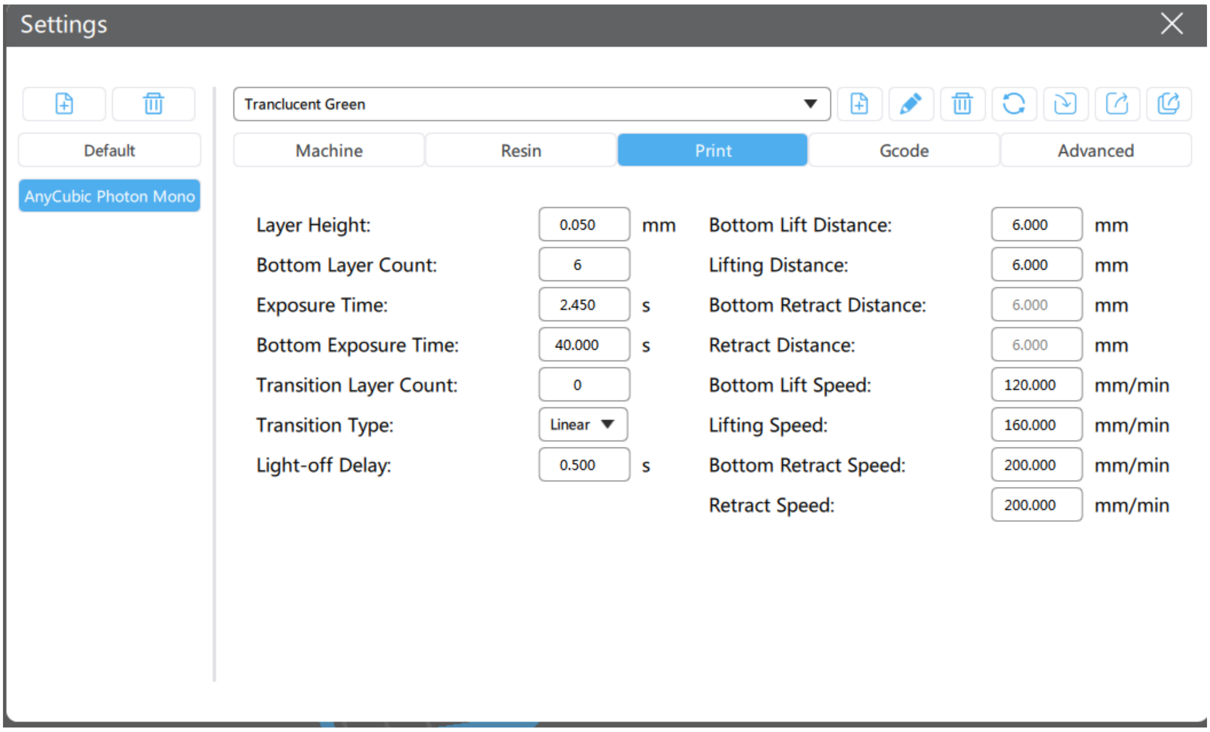


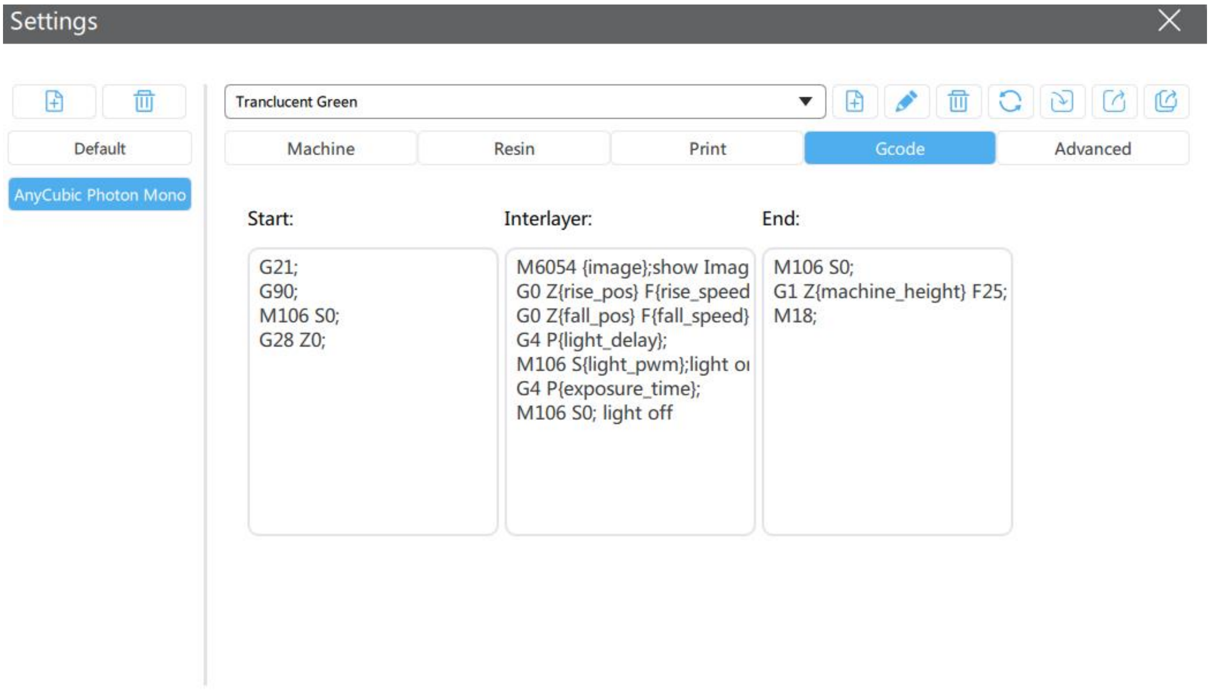

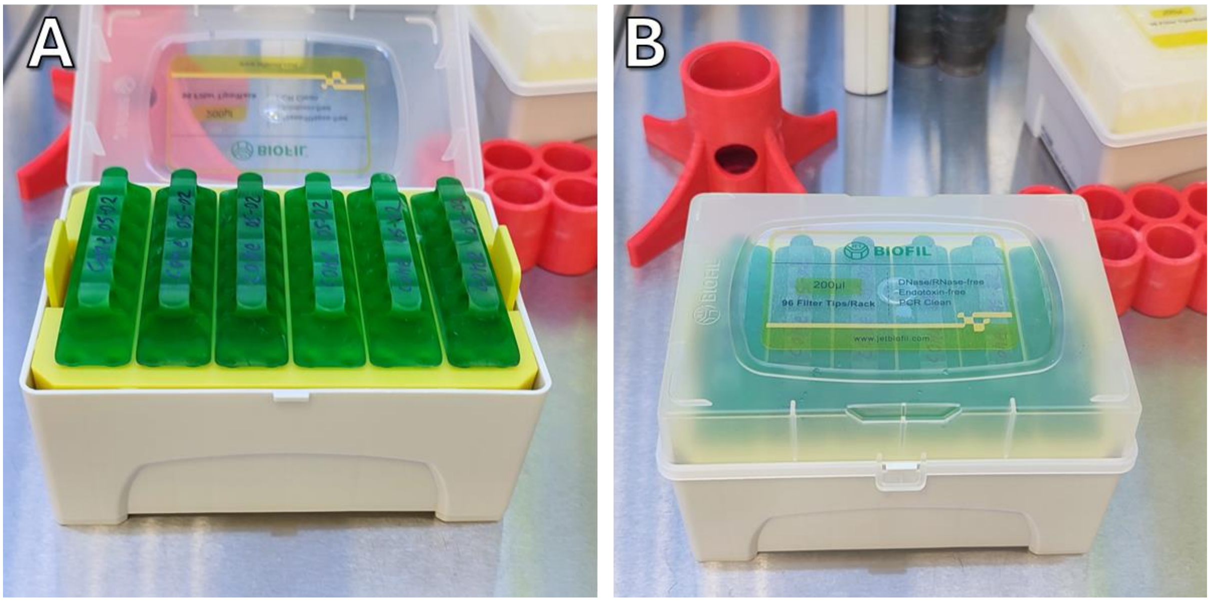


*Figure S1. Storing stamps for 96-well plates in a tips box*

### S2. Key dimensions

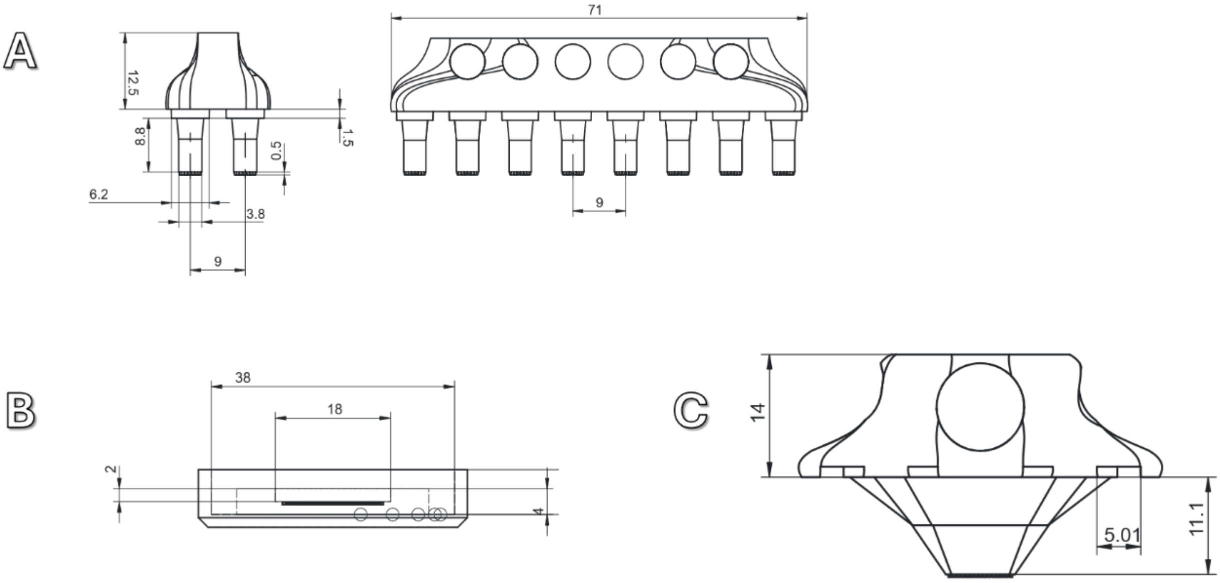


*S2. Key dimensions of the stamps and mold*

### S4. Bubbles in an agarose mold

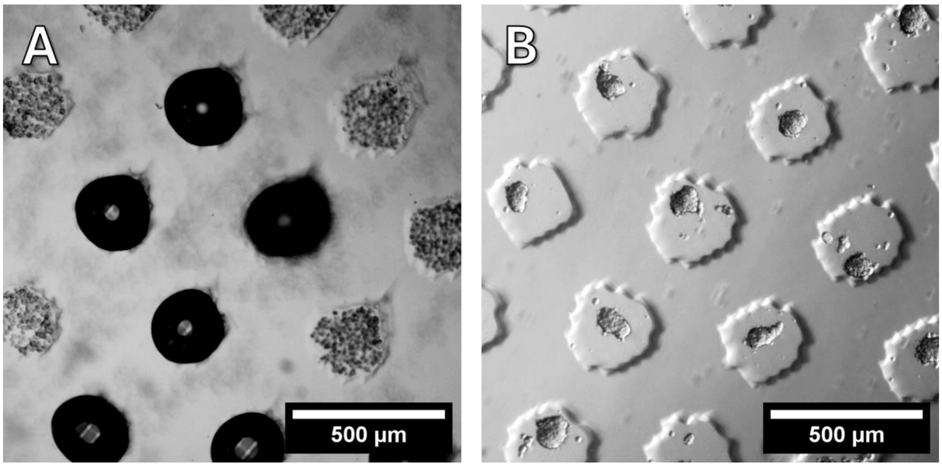


Agarose molds with a microwell diameter of 300 µm filled with nutrient medium with cells. Without (A) and with treatment (B) with isopropyl alcohol. A shows air bubbles filling the microcells.

### S5. Protocols

**The choice of silicone**

This is the tricky part. Staying in the ideology of accessibility and DIY, I used the usual model two-component silicone available in the hobby store. Two-component platinum-based silicones definitely do not work - there is some kind of reaction between the silicone and the printed part, and the silicone does not polymerize. Tin-based silicones remain, they generally work well. However, not all, I have met at least one that sticks to MSLA printed parts tightly and then it is impossible to separate the finished molds. Accordingly, if someone decides to make such molds, silicone will have to be selected empirically. And I didn't use any chemical separators because I didn't want to add their remnants to the biological medium.

#### 6-plate/dish microwell production

**Tools and equipment**

- Vacuum chamber
- Two-component silicone (that you are sure about)
- Dye for silicone (with it is better to see how well mixed silicone, as well as you can use different colours for colour coding
- Disposable wooden or plastic spatulas.
- Wipes and gloves (this process will inevitably be messy)

**Process**

The total volume of silicone depends on how many molds you make at a time. Also silicone is quite hard to transfer quantitatively, it's a skill. Therefore, it is better to take with reserve. You will need two portions of silicone. The first portion is about ten grams per mold, the second twenty.

1. Mix the first (smaller) portion of silicone with the hardener and dye

2. Transfer to the molds and spread evenly with the spatula, making sure that the silicone covers the entire area evenly.
3. Transfer the molds to the vacuum chamber and evacuate, then let air in and evacuate again. Leave under vacuum for 3-5 minutes
4. Mix the second batch of silicone with hardener and dye (another colour can be used)
5. Release air, take the molds out of the vacuum chamber and pour the second portion of silicone into them
6. Remove the excess with a spatula
7. Leave for 24 hours (or as recommended by the silicone manufacturer) until curing

**Gel mold productuion**

**Before starting work**

Nothing is sterile by default or even intended to be. In the laminar box, it is highly recommended to treat the molds and stamps with 70% isopropyl or ethyl alcohol before the first use. You can use a spray bottle, but it is better to treat them by immersing them in a alcohol for a few minutes. I would not advise autoclaving, and molds and stamps, in principle, should withstand such treatment, but without guarantees. It is most convenient to store it in the box of tips, this box can also be treated with alcohol and stored inside the laminar box while working.

**Tools and equipment**

- Laminar flow cabinet
- Microwave or other way to melt the gel
- Molds
- Plastic pasteur pipettes (2-5 ml)
- Tips for 10-200 ul pipette
- Agar/agarose solution with a concentration of about 1-3% mass. The minimum volume required per plate is about 5 ml, but it is better to melt 10-20 so that the melted gel in the bath does not solidify so quickly.

**Process**

1. Melt the agar/agarose solution, preferably bring to a boil
2. Using a paster pipette, transfer the gel to the silicone molds, spreading it gently over the surface, large bubbles can be sucked up with the pipette
3. If you use a silicone mold with a small size of micropins (300 um), air bubbles often get stuck on them, which spoils the agarose mold. To get rid of them mold should be ‘ironed’ for example with a 10-200 tip (Watch the video below).
4. Agarose hardens for 10-20 minutes, after which the mold can be carefully removed from the mold and transferred to a plate cell 6-well plate cell or a suitable dish.
5. Agarose molds are ready for cells

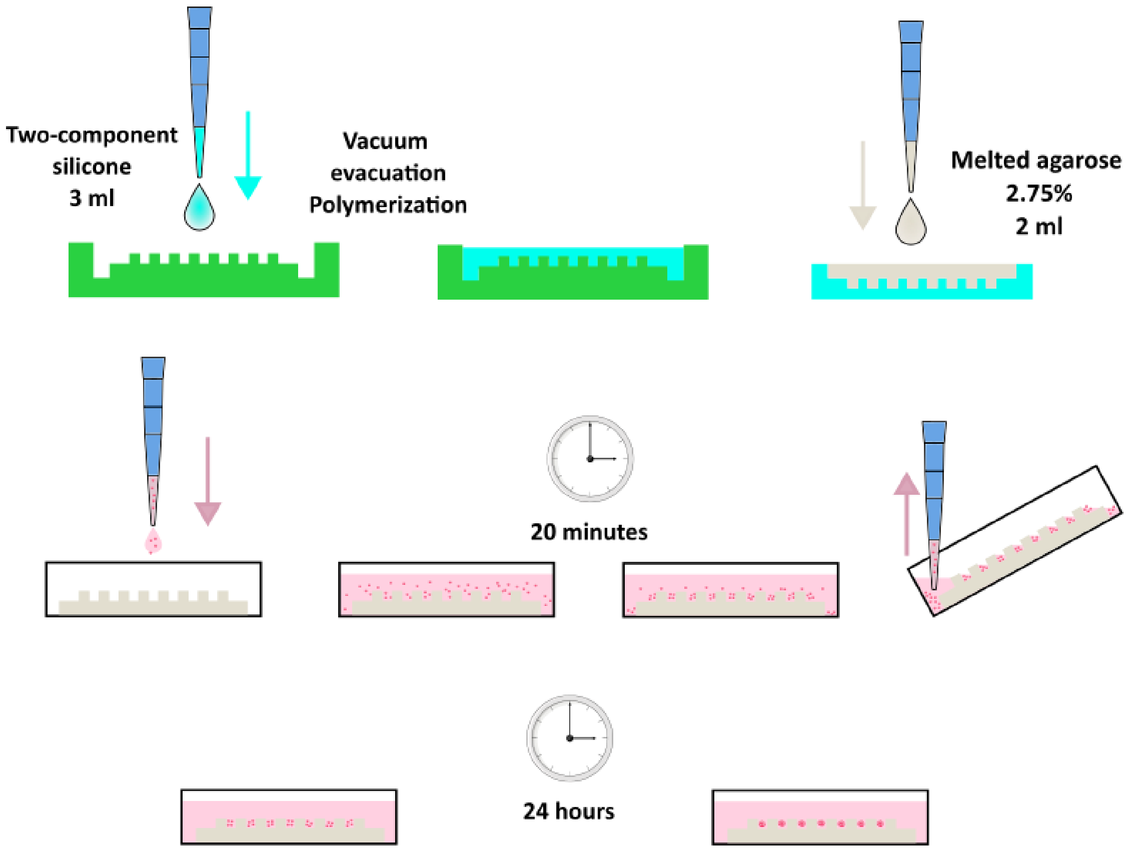

*Fig. Scheme illustrating the process of spheroids manufacturing in molds for 6-well plates*

**Cell seeding**

1. Detach the cells using the laboratory procedure.
2. Count the cells in suspension and dilute to the required number per microwell. About 300 µl can be poured into the recess of the agarose mold.
3. Transfer the cell suspension into the recess of the mold
4. Allow 15-30 minutes for the cells to settle under gravity.
5. Tilt the plate with molds and carefully remove the medium with excess cells that did not fall into the wells
6. Add fresh medium, including the empty space around the mold.
7. Place the plates in the incubator until the spheroids consolidate for 24 hours

### 96-plate microwell production

**Before starting work**

Nothing is sterile by default or even intended to be. In the laminar box, it is highly recommended to treat the molds and stamps with 70% isopropyl or ethyl alcohol before the first use. You can use a spray bottle, but it is better to treat them by immersing them in a alcohol for a few minutes. I would not advise autoclaving, and molds and stamps, in principle, should withstand such treatment, but without guarantees. It is most convenient to store it in the box of tips, this box can also be treated with alcohol and stored inside the laminar box while working.

**Tools and equipment**

- Laminar flow cabinet

- Microwave or other way to melt the gel
- Stamps - 6 pieces
- Multichannel pipette (capable of dispensing 50 µl of liquid) - 1 piece
- Tub for multichannel pipette
- Tips for pipette
- 96-well plate
- Agar/agarose solution with a concentration of about 1-3% mass. The minimum volume required per plate is about 5 ml, but it is better to melt 10-20 so that the melted gel in the bath does not solidify so quickly.

**Process**

Working with molten agar/agarose requires care and, more importantly, speed, so it is better to prepare everything within reach in advance

1. Set the pipette to 50 µl
2. Melt the agar/agarose solution, preferably bring it to a boil
3. In a laminar flow cabinet, pour the agarose into the tub for the multichannel pipette
4. Use the pipette to transfer 50 µl into the two columns of the plate and immediately place a stamp in these columns. Make sure that the stamp is fully inserted (there is a tab on it centring it in the well, make sure it is inserted). It is necessary to do everything as quickly as possible, so that the gel does not freeze in the spouts of the dispenser
5. Pour two more columns, set the stamp, repeat until all the stamps are installed
6. Wait for the gel to solidify (10-15 minutes, depending on agar/agarose and room temperature)
7. Remove the stamps. This is the most critical step. It is most convenient to turn the plates with the short part towards yourself and remove the stamps with two hands from the sides. Gently rock the stamp back and forth before removal. If the gel concentration is optimal and the gel is fully cured, the stamp should come out clean and free of gel pieces and the microwells should remain intact inside the tablet. A few wells per plate usually come with bubbles, it's a matter of luck and it's hard to do anything about it. A video of the extraction process is shown below
8. The microwells are ready for seeding of cells

**Cell seeding**

Before conducting a serious experiment, it is best to determine the optimal number of cells needed to form spheroids. They should not be too many or the extra cells in the well will form spheroids of uncontrolled size outside the microwells. The figure below shows spheroids formed from HeLa with different numbers of cells per cell. In this case about 1000-500 cells per microwell is optimal, when dense spheroids are formed after a day and there are practically no cells outside the cells.
You will need standard equipment and supplies for working with cell cultures, as well as a microwell plate made in the previous step, and:

- Centrifuge or vortex with a plate rotor (optional)

1. Detach the cells using the laboratory procedure.
2. Count the cells in suspension and dilute to the required number per microwell. One plate well can be filled with 50 - 100 µl of medium.

3. Transfer the cell suspension to the tub for the multichannel pipette
4. Using the multi-channel pipette, transfer the cell suspension into the plate
5. Either centrifuge at minimum speed to allow the cells to settle to the bottom of the microwells or hold the plates for 15-30 minutes to allow the cells to settle by gravity.
6. Some cells are deposited between the cells or above or at the top of the plate cell where the stamp entered the agarose. To remove excess cells the plate should be tilted at 45 degrees and carefully remove the medium with excess cells with a multichannel pipette, pipetting several times.
7. 50-100 µl of new medium can be poured in place of the removed medium
8. Place the plates in the incubator until the spheroids consolidate for 24 hours

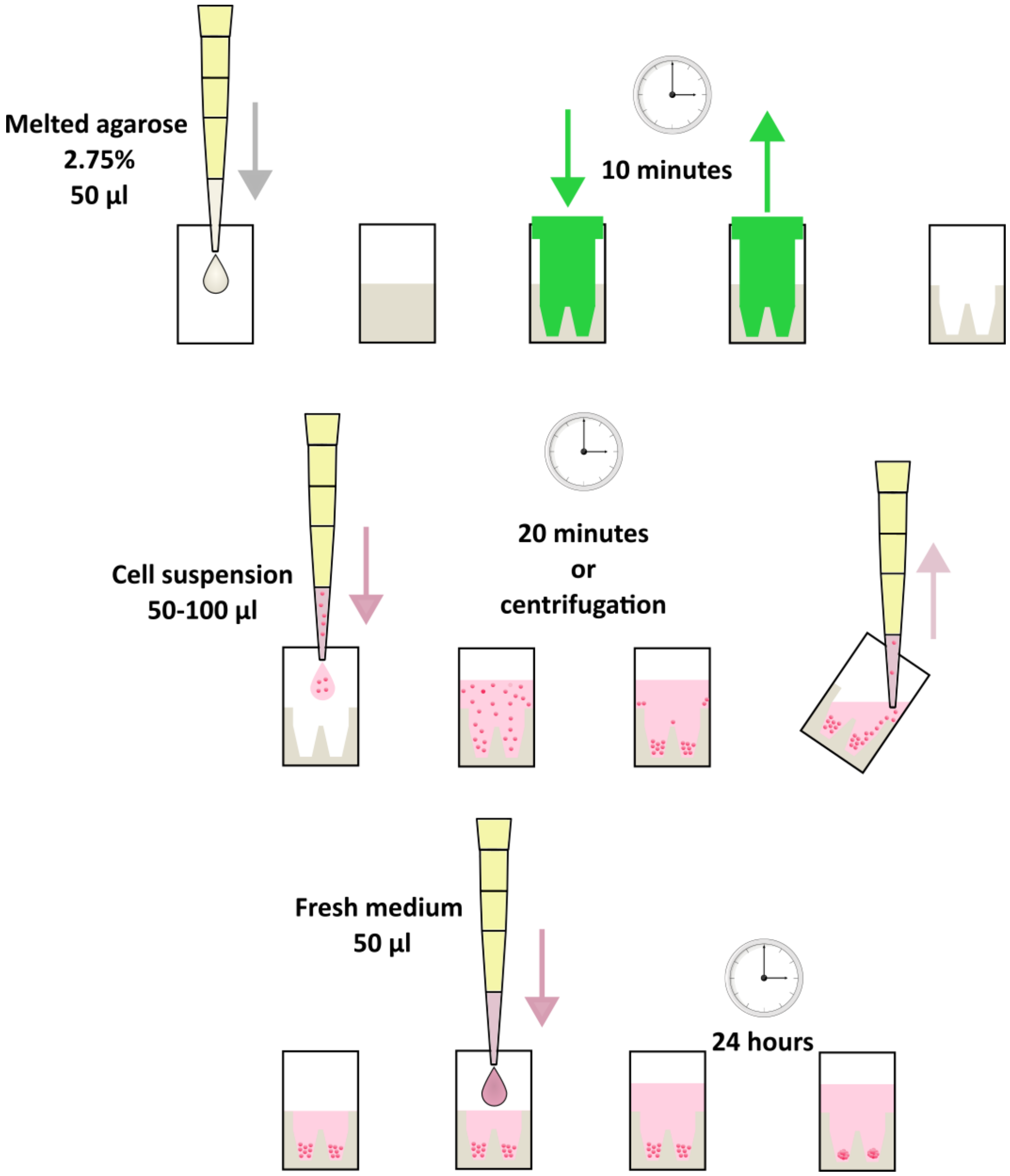


*Fig. Scheme illustrating the process of spheroids manufacturing with stamps for 96-well plates*

### Glass bottom dish microwell production

**Before starting work**

Nothing is sterile by default or even intended to be. In the laminar box, it is highly recommended to treat the molds and stamps with 70% isopropyl or ethyl alcohol before the first use. You can use a spray bottle, but it is better to treat them by immersing them in a alcohol for a few minutes. I would not advise autoclaving, and moulds and stamps, in principle, should withstand such treatment, but without guarantees. It is most convenient to store it in the box of tips, this box can also be treated with alcohol and stored inside the laminar box while working.

**Tools and equipment**

- Laminar flow cabinet
- Microwave or other way to melt the gel
- Stamps
- Plastic pasteur pipettes (2-5 ml)
- Tips for pipette
- Glass bottom dishes plate
- Agar/agarose solution with a concentration of about 1-3% mass. The minimum volume required per dish is about 1 ml, but it is better to melt 10-20 so that the melted gel in the bath does not solidify so quickly.

**Process**

Working with molten agar/agarose requires care and, more importantly, speed, so it is better to prepare everything within reach in advance

1. Melt the agar/agarose solution, preferably bring it to a boil
2. In a laminar flow cabinet, pour the agarose into the tub for the multichannel pipette
3. Use the pipette to transfer 1-2 ml into the dish of the plate and immediately insert a stamp. Make sure that the stamp is fully inserted (there is a tab on it centring it in the well, make sure it is inserted)
4. Wait for the gel to solidify (10-15 minutes, depending on agar/agarose and room temperature)
5. Remove the stamp.
6. The microwells are ready for seeding of cells

**Cell seeding**

Before conducting a serious experiment, it is best to determine the optimal number of cells needed to form spheroids. They should not be too many or the extra cells in the well will form spheroids of uncontrolled size outside the microwells. The figure below shows spheroids formed from HeLa with different numbers of cells per cell. In this case about 1000-500 cells per microwell is optimal, when dense spheroids are formed after a day and there are practically no cells outside the cells.

You will need standard equipment and supplies for working with cell cultures, as well as a microwell plate made in the previous step, and:

- Centrifuge or vortex with a plate rotor (optional)

1. Detach the cells using the laboratory procedure.
2. Count the cells in suspension and dilute to the required number per microwell. One dish can be filled with 200 - 400 µl of medium.
3. Using the pipette, transfer the cell suspension into the plate
4. Place the dishes in the incubator until the spheroids consolidate for 24 hours

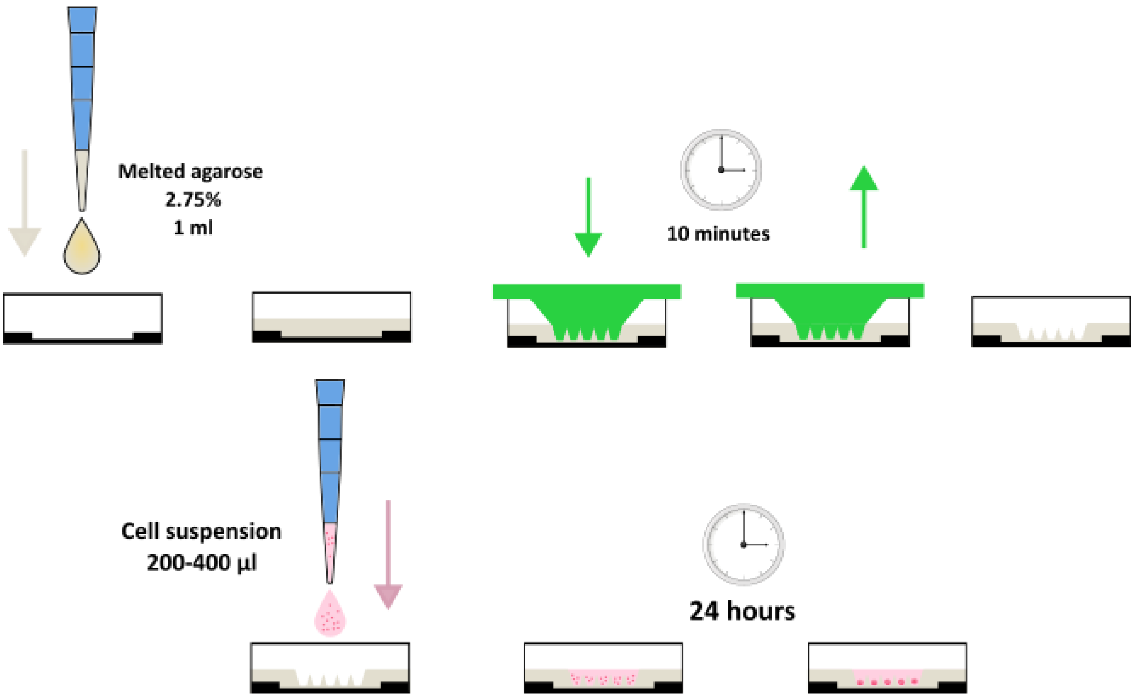

*Scheme illustrating the process of spheroids manufacturing with stamps for glass-bottom dishes*

### S6. Basic of MSLA 3d printing

#### Principles of MSLA 3d printing

MSLA (masked stereolithography) printer is based on stereolithography to produce three-dimensional models. The process is carried out through the polymerization of photosensitive compounds under the influence of light, and the mask controls which part of the composition will be illuminated.

There are several main components in a typical MSLA 3D printer. The liquid photopolymer is poured into a vat, the bottom of which is made of a transparent film made from fluorinated polymers (FEP film), which ensures minimal adhesion. The bath is installed in the printer on top of an LCD screen, under which an array of LEDs (with a wavelength of 405 nm) is installed.

While the printer is in use, a platform is automatically lowered into the resin-filled vat. Light from photodiodes is then projected through an LCD screen onto the platform. The screen's individual pixels control the passage of light, functioning as a mask (hence the "M" - in "MSLA") in the printing process.

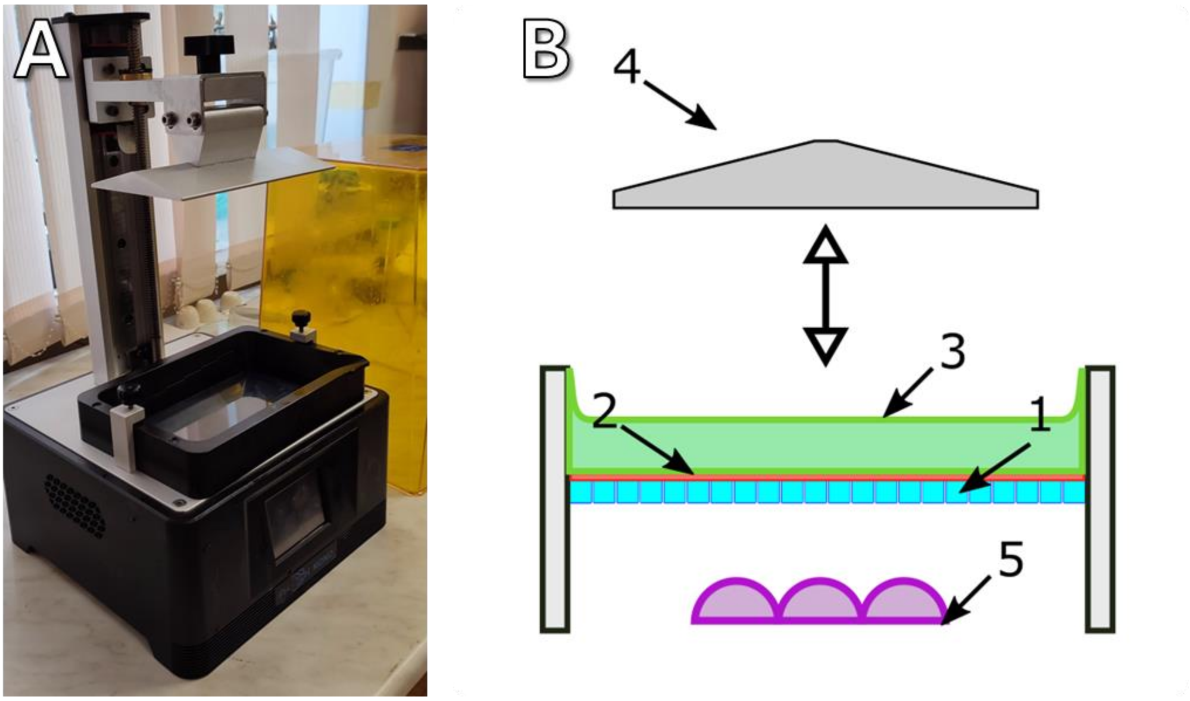


*Figure S1. Anycubic Photon Mono 3d printer (A) with the protective cap removed and an MSLA diagram of the printer showing the main construction components (B).*

At the first printing cycle the stage is lowered almost close to the FEP-film, at a distance of one layer (usually in 3d printers of hobby class it is 50-25 μm), after which the irradiation is switched on. The layer of photopolymer between the film and the stage polymerizes and adheres to the stage, and the process is selective, only in those areas where the light passes through the LCD screen.

When one layer of the part is illuminated and stuck to the stage, the stage is raised to a certain height, necessary for fresh photopolymer to flow under it, then lowered back down, with the new distance from the FEP to the stage being greater by the height of one layer, after which a new layer is illuminated (polymerized), sticking this time to the already solid first layer. The process is repeated the necessary number of times to print the entire part.

#### Angle of printing

In this whole process of MSLA 3d printing, there are a number of non-obvious factors that need to be considered for successful printing of various objects with small surface features [https://doi.org/10.1002/bit.28031].

- FEP film has minimal adhesion to the photopolymer, but not zero, and, in addition, over time the film degrades and the printed objects begin to stick to it more and more, which eventually leads to the object coming off the bed. The adhesion is also aggravated by the hydrodynamic pressure generated by the upward movement of the stage.
- If the area of the object that is adhered to the bed is too small, it can also tear off.

A common way to eliminate such problems in stereolithographic printing (not only MSLA, but other types as well) is to place objects at an angle, which minimizes the contact area with the film. At the same time, to increase adhesion of the object to the stage, supports are added to the model, which ensure successful printing and are separated from the object after printing.

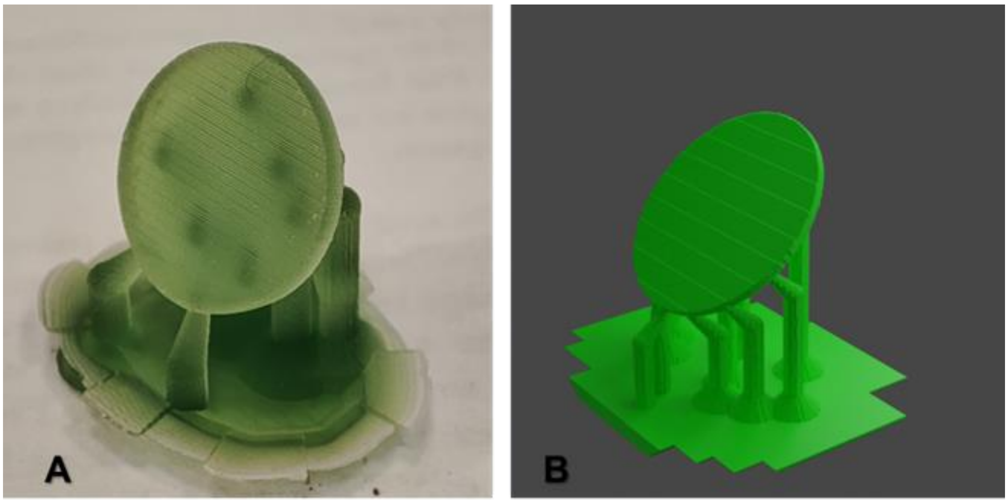


*Figure. Macrophotograph (A) and (B) 3d rendering of a test object printed at a 60 degree angle to a table with supports*

However, the above-mentioned ways to achieve successful printing have their significant disadvantages. The fact is that the MSLA 3d printer has limited resolution in all three axes. On the xy axis the resolution is limited by the pixel size of the LCD screen (approximately 40x40 μm, slightly different for different printer models available on the market), and on the z axis the resolution is limited by the layer height (25-50 μm, depending on the settings).

This leads to the fact that the object printed by the printer is not a perfect reproduction of the digital model, the software (slicer) used for printing approximates this model with voxels, which leads to various artifacts of this transformation, as shown in the diagram below.

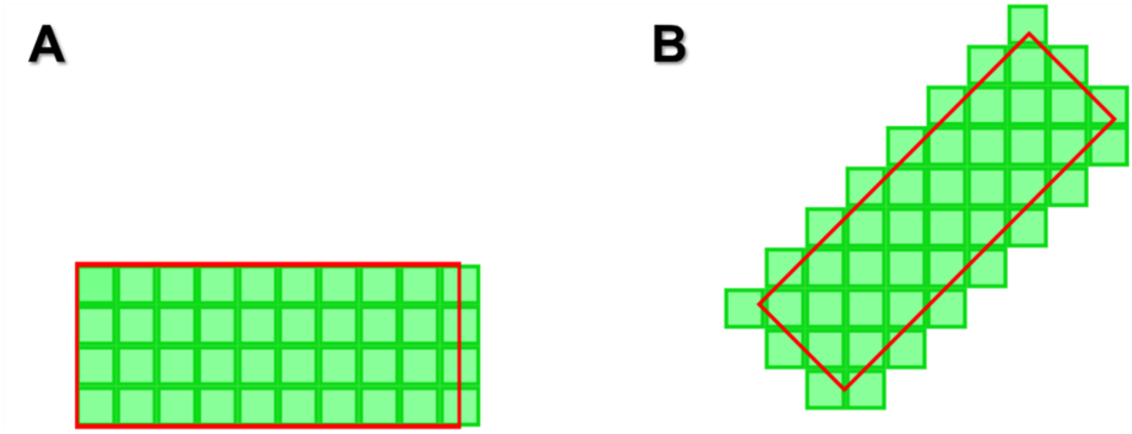


*Figure. Illustration of the process of approximation of a flat model by voxels*

The scheme is presented in 2d to simplify perception. Red denotes the original model embedded in the slicer, green - the resulting model, approximated by voxels (in case of 2d illustration - pixels). Object A is located parallel to the printing plane, object B is located at an angle of 45 degrees (the diagram does not indicate the support necessary for printing in this orientation).

It is shown how the conversion to voxels performed by the slicer can distort the geometric dimensions of the object if it is slightly smaller or larger than an integer number of pixels - this can be seen on the right side of the object A.

The example of object B demonstrates how flat faces are distorted, in case of printing at an angle - the slicer approximates them with steps, which leads to a significantly inhomogeneous surface of the plane.

Below are the images of the test object in the 3d software and how it will look when printing. Also, a clarification must be made that this is still an idealistic digital representation of the object. In reality, to the already distorted model will be added various mechanical and optical artifacts of the printing itself, which further affects its geometry.

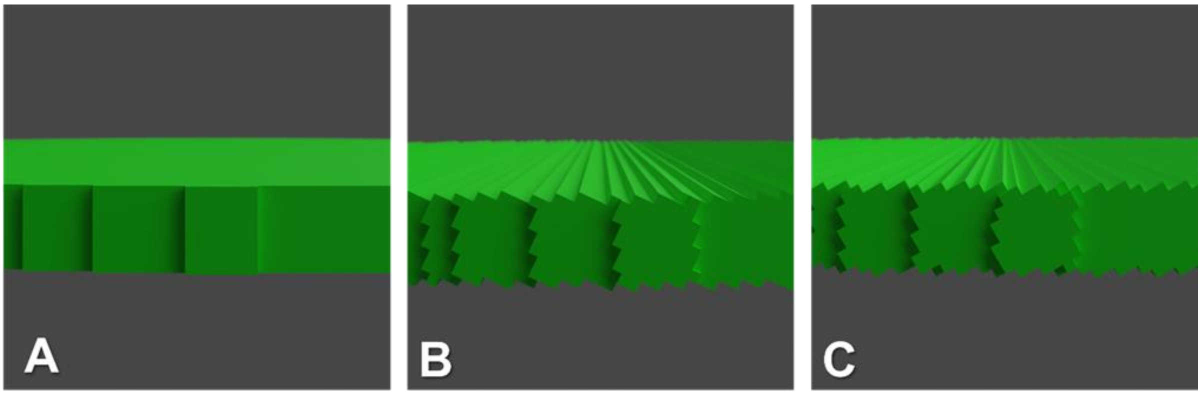


*Figure. 3d render test objects, profile view. A - the object is oriented in the print plane, B - at an angle of 25 degrees, C - at an angle of 60 degrees*

If you look at the printed test objects in a microscope, the picture looks a bit different. The objects printed at an angle to the print plane look similar to the way they looked on the rendering - there are noticeable clear steps. However, the test object printed parallel to the plane also looks inhomogeneous, a clear square grid is visible. This grid is the result of inhomogeneous illumination in the body of each individual pixel of the LCD matrix and is an unrecoverable artefact of this type of printing. The cause of these artefacts is illustrated in the figure below. The letter A on it denotes the ideal process of illumination of the sample, and the letter B denotes how the process goes in reality. Number 1 in the illustration denotes the illumination structure of the pixel area, number 2 the structure of the light beam and number 3 the single layer of the polymerising model.

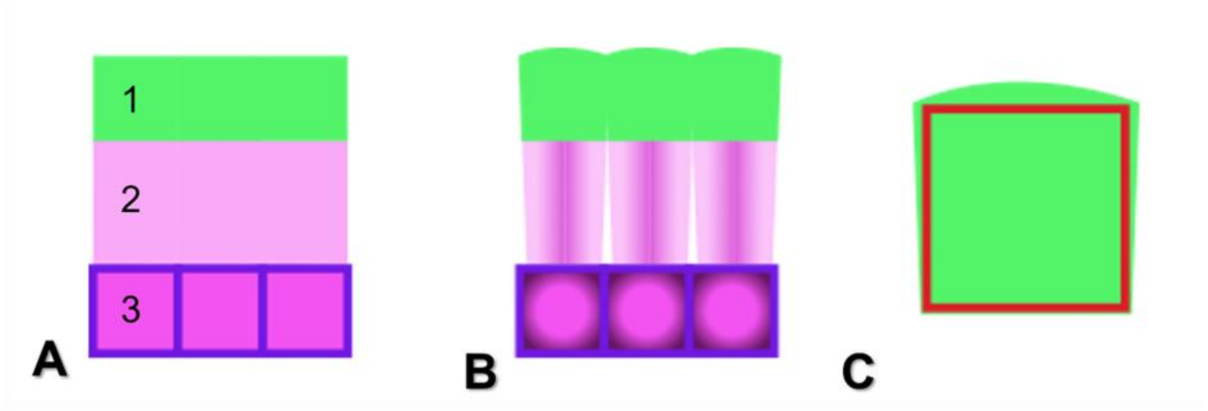


*Schematic illustration of the process of pixel shape distortion. A - unrealistic ideal representation of the process - 1 - photopolymer layer, 2 - light beam, 3 - LEDs. B - schematic illustration of how the process goes in reality, C - distorted shape of each particular voxel, red frame - theoretical voxel shape, green figure - how it will turn out in reality.*

Under ideal conditions (A) the pixel of LCD matrix transmits light absolutely uniformly, the light without distortion goes to the photopolymer and illuminates it uniformly. But in reality, the pixel transmits more light in the centre and less at the edges, which in turn leads to inhomogeneity of the light flux. In addition, light in the medium is subject to diffraction, resulting in beam broadening. The combination of these effects results in a pixel shape that differs from the ideal pixel shape, which is shown in Figure C. The ideal pixel shape is shown in red, the green shows how it is obtained in reality.
Measurement with a confocal microscope showed that the heterogeneity in the height of one pixel on the printed test object is of the order of 4-5 μm from the periphery to the centre,

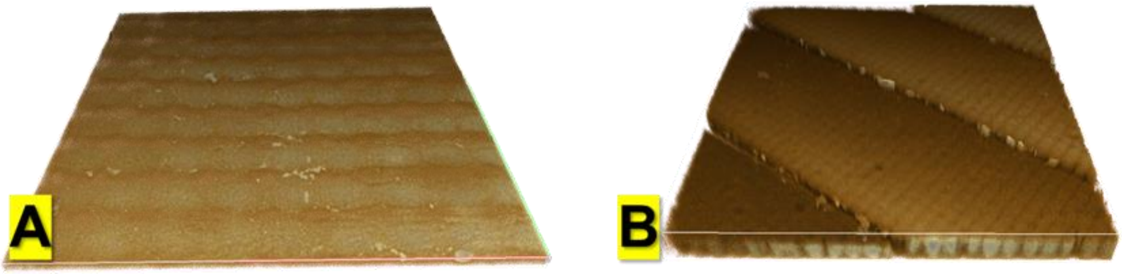


3D reconstruction of the surface of the test object printed parallel to the printing plane (A) and at an angle of 60 degrees (B). 3D reconstruction was made with the Drishti (2.7) software.

### S7. Test object

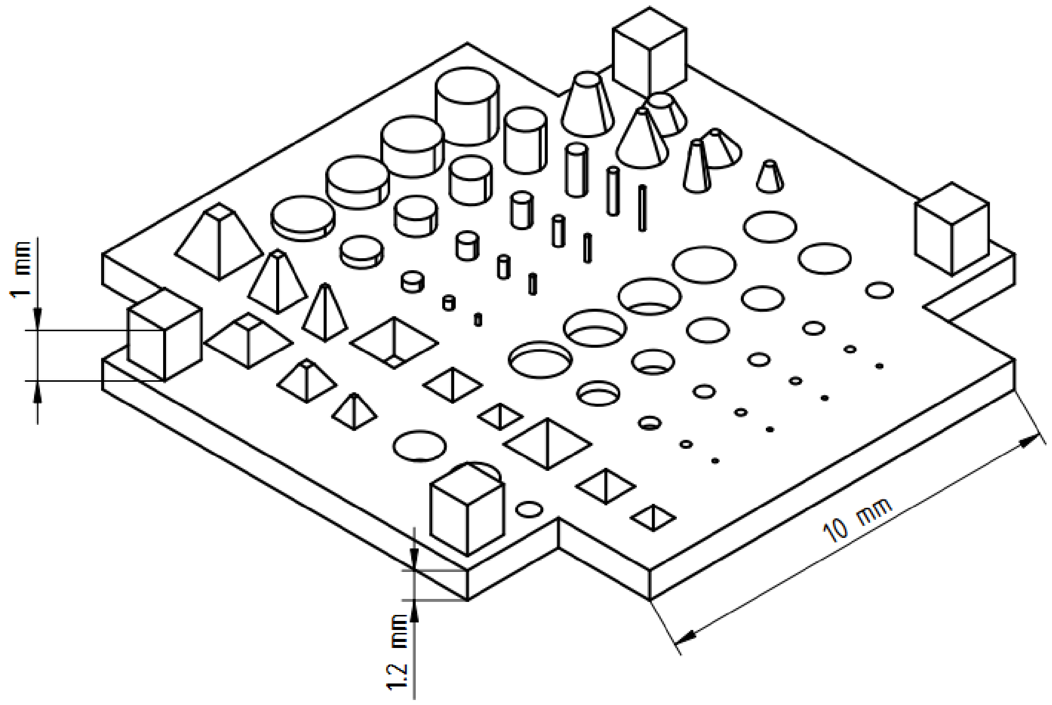

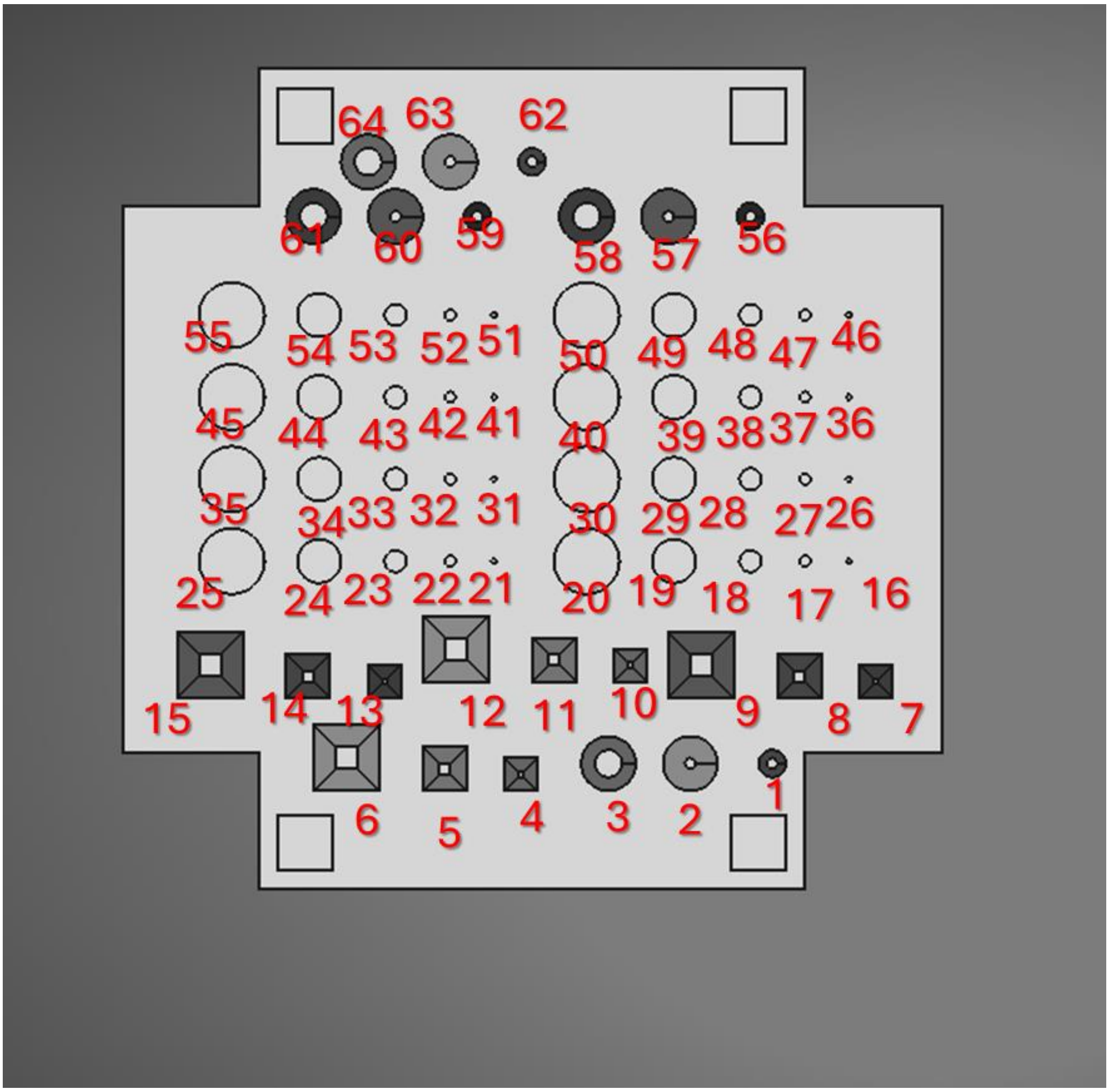


| N | Shape | Dimensions | |
|---|---|---|---|
| 1 | Cone well | 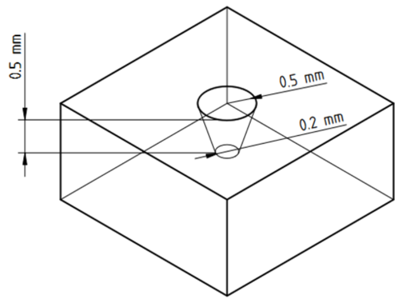 | 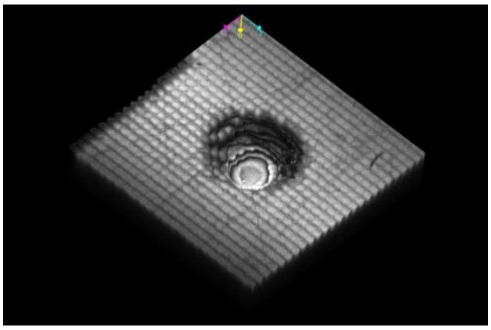 |

| | | | |
|---|---|---|---|
| 2 | Cone well | 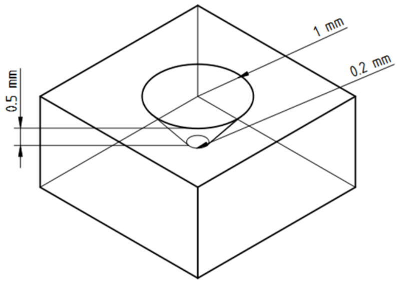  | 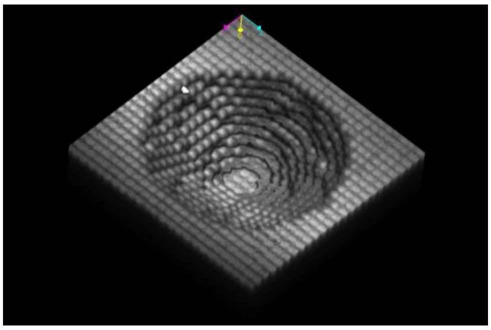 |
| 3 | Cone well | 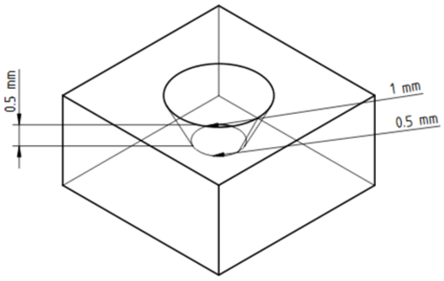  | 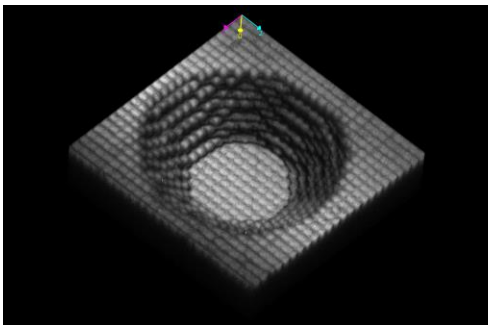 |
| 4 | Piramide | 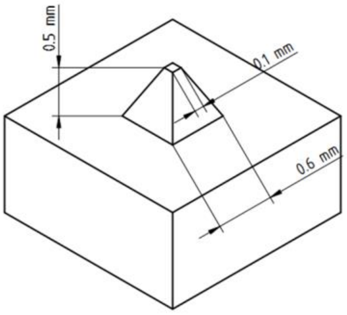  | 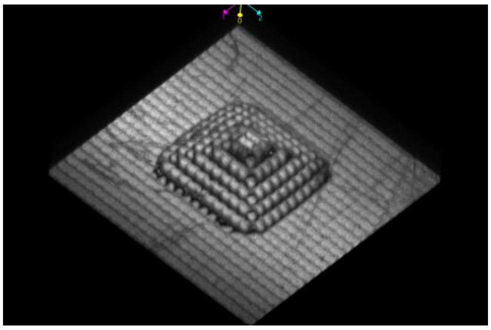 |
| 5 | Piramide | 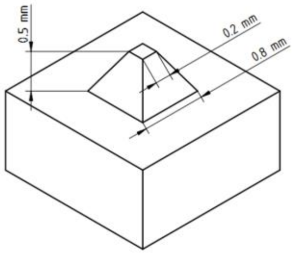  | 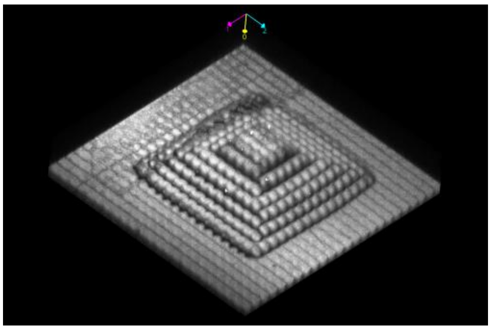 |

| | | | |
|---|---|---|---|
| 6 | Piramide | 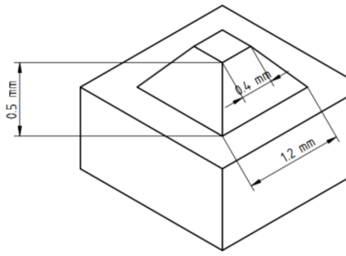  | 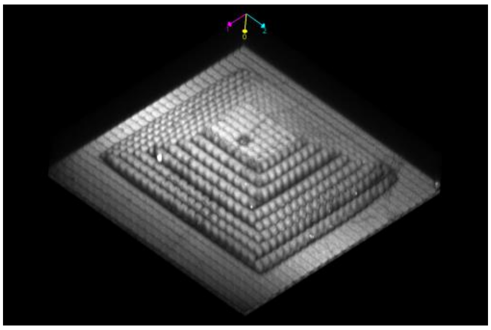 |
| 7 | Piramide well | 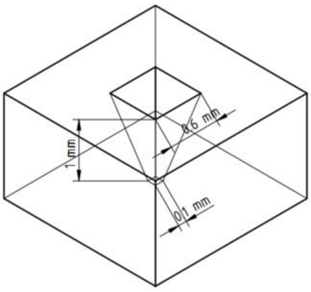  | 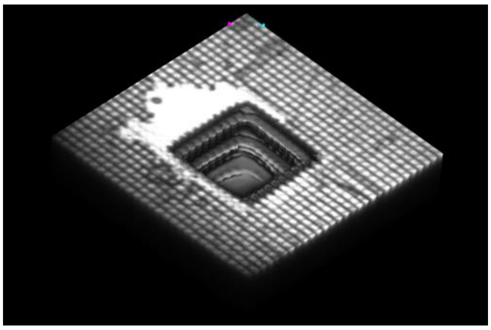 |
| 8 | Piramide well | 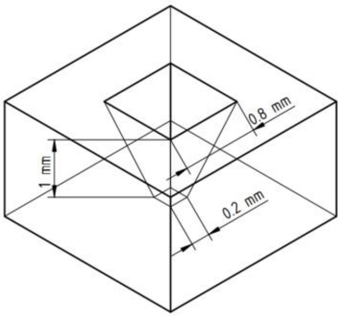  | 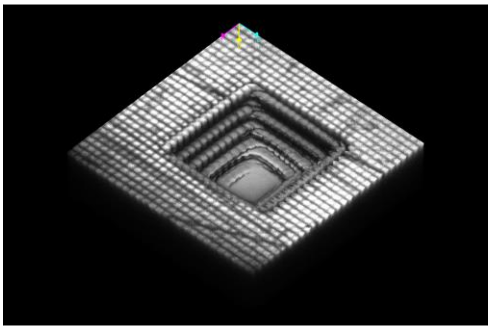 |
| 9 | Piramide well | 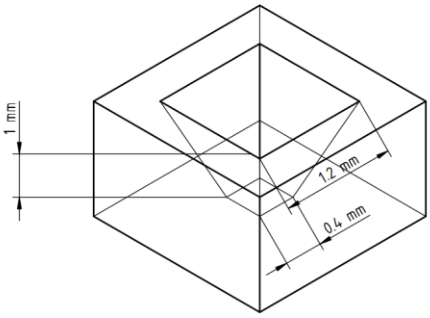  | 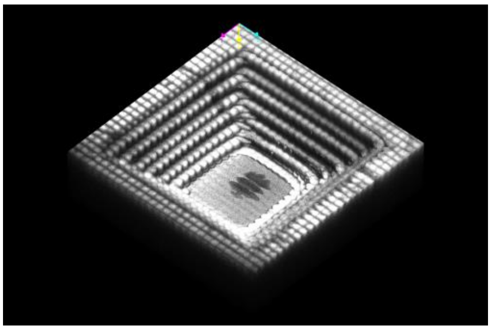 |

| | | | |
|---|---|---|---|
| 10 | Piramide well | 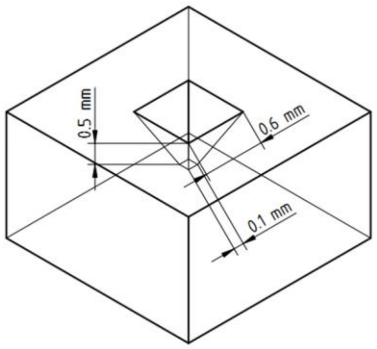  | 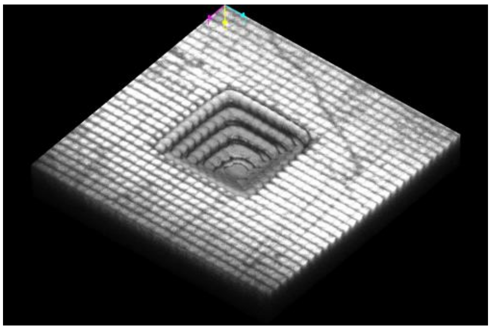 |
| 11 | Piramide well | 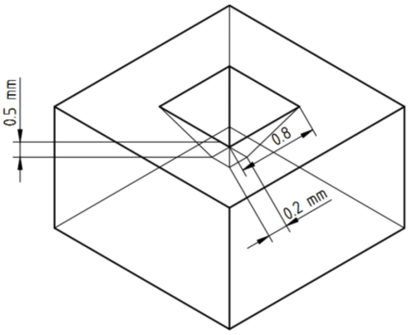  | 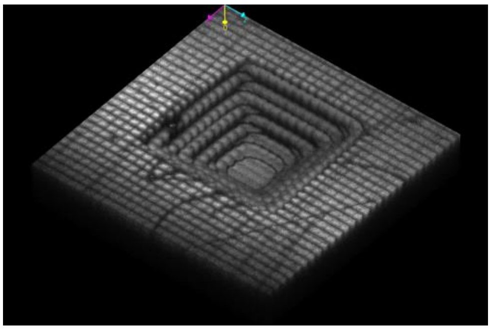 |
| 12 | Piramide well | 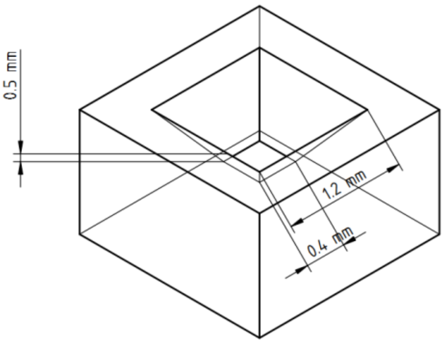  | 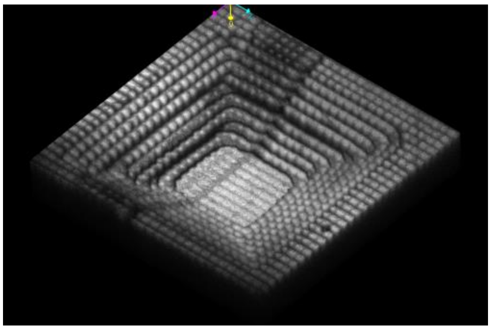 |
| 13 | Piramide | 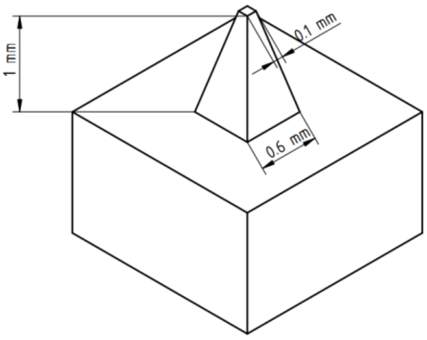  | 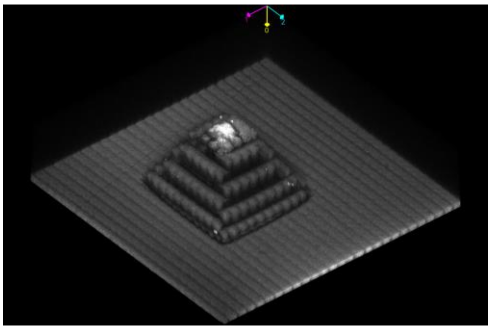 |

| | | | |
|---|---|---|---|
| 14 | Piramide | 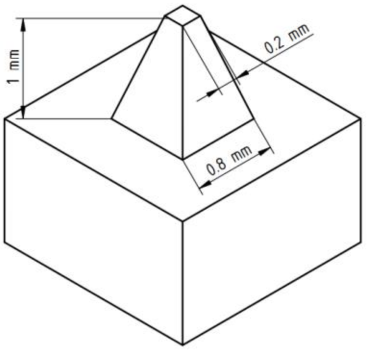  | 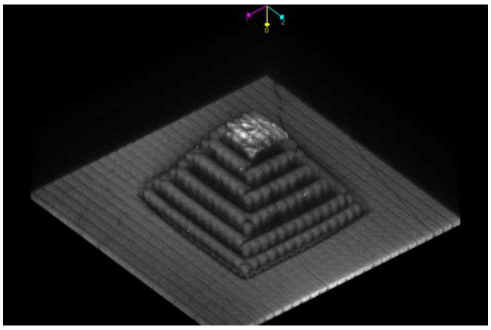 |
| 15 | Piramide | 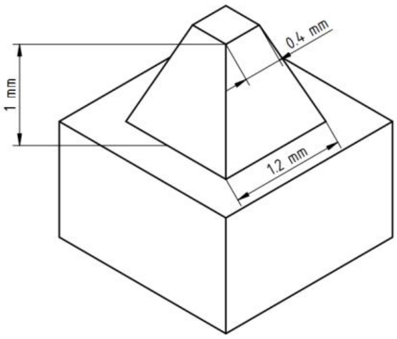  | 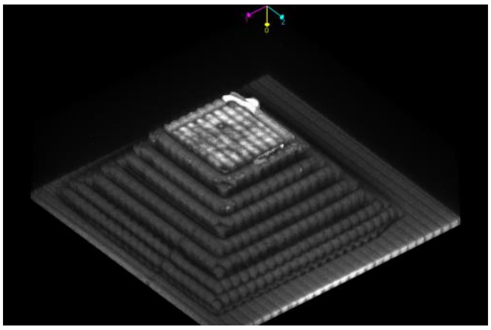 |
| 16 | Cylinder well | 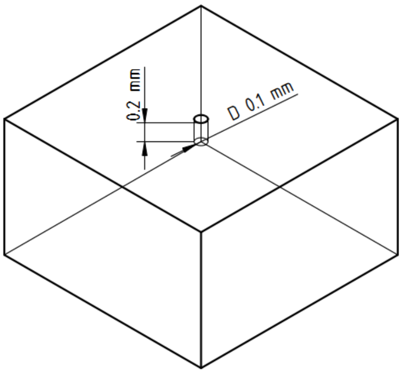  | Printing failure |
| 17 | Cylinder well | 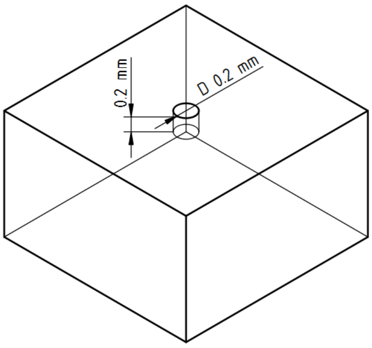  | 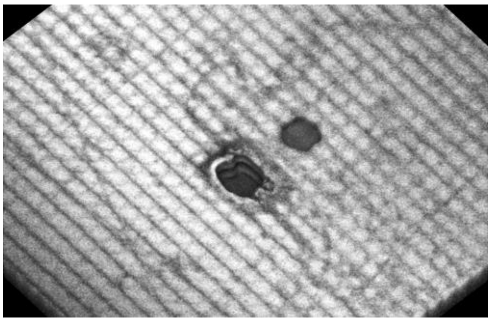 |

| | | | |
|---|---|---|---|
| 18 | Cylinder well | 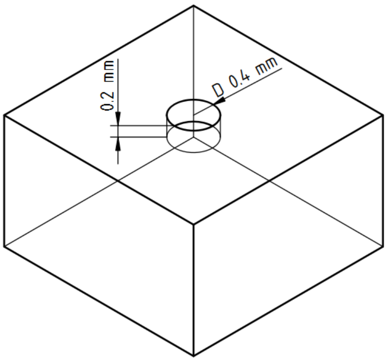 | 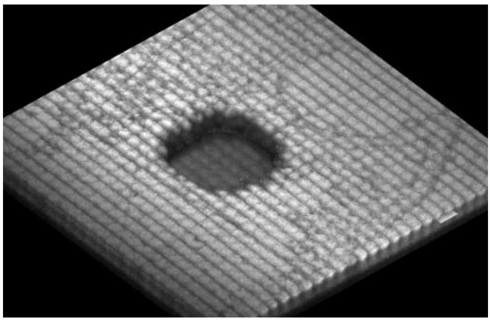 |
| 19 | Cylinder well | 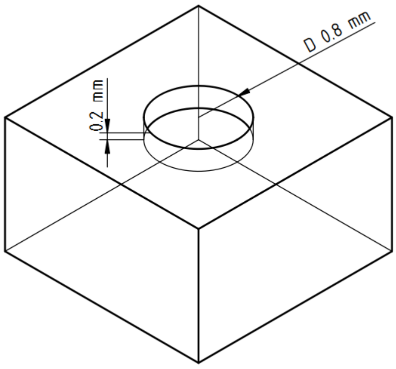 | 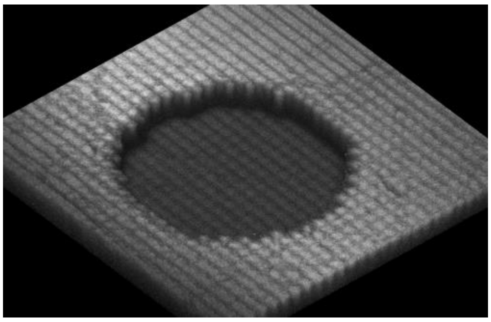 |
| 20 | Cylinder well | 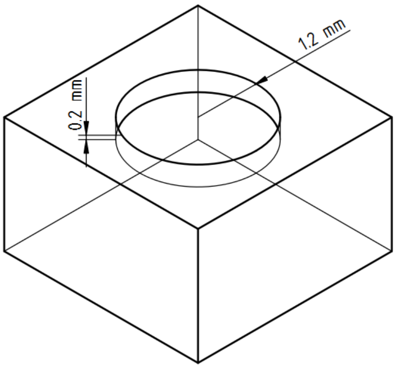 | 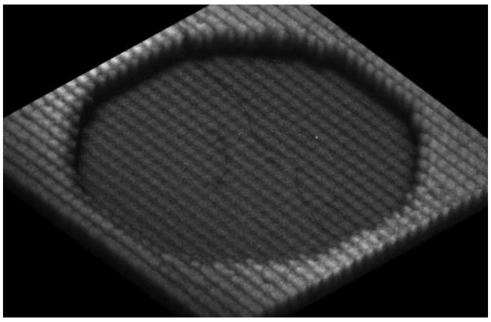 |
| 21 | Cylinder | 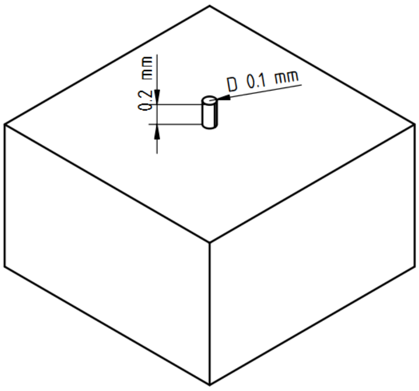 | Printing failure |

<table>
<tr><td>22</td><td>Cylinder</td><td>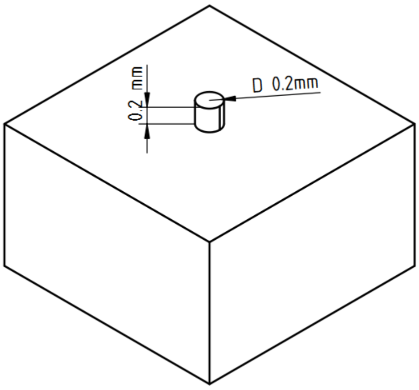</td><td>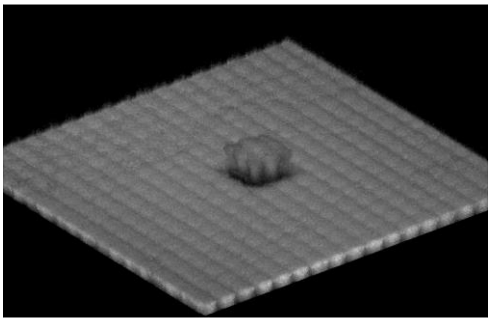</td></tr>
<tr><td>23</td><td>Cylinder</td><td>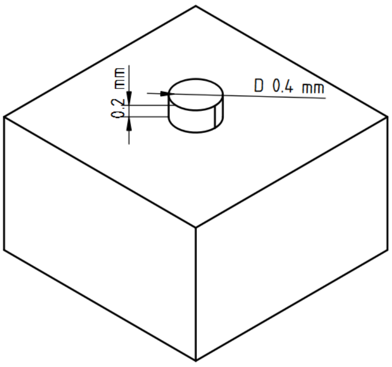</td><td>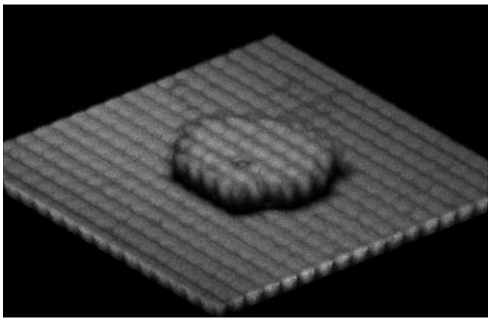</td></tr>
<tr><td>24</td><td>Cylinder</td><td>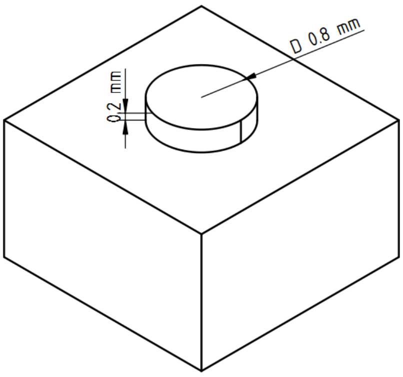</td><td>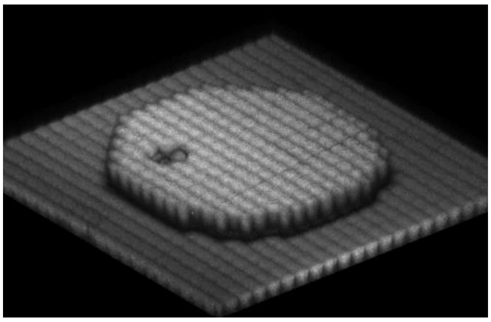</td></tr>
</table>

| | | | |
|---|---|---|---|
| 25 | Cylinder | 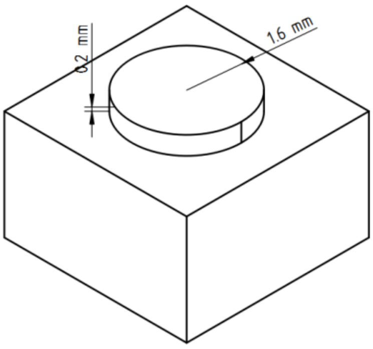  | 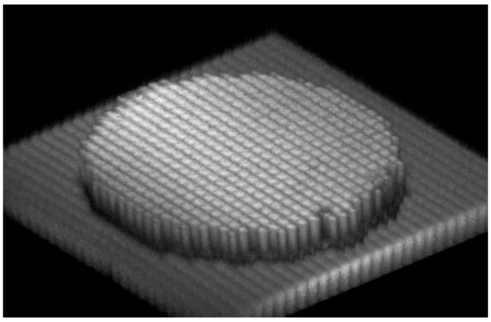 |
| 26 | Cylinder | 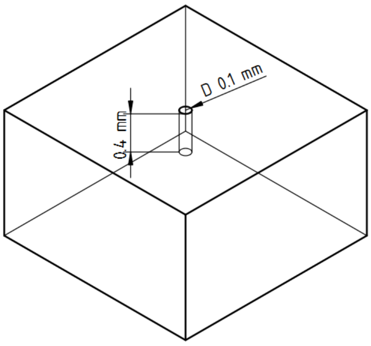  | Printing failure |
| 27 | Cylinder well | 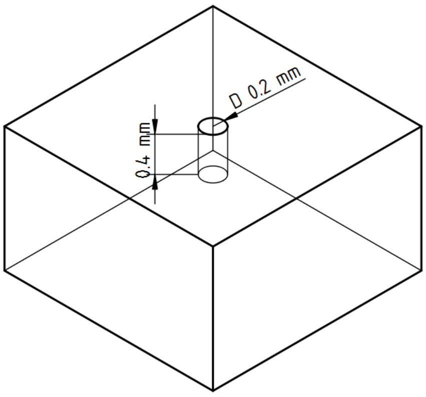  | 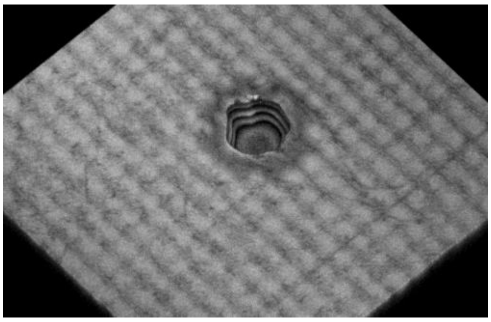 |
| 28 | Cylinder well | 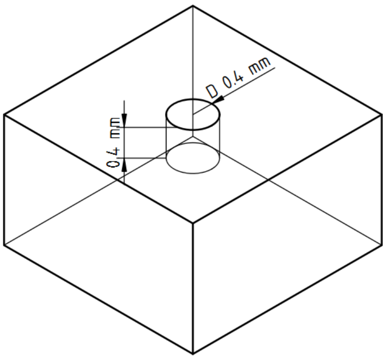  | 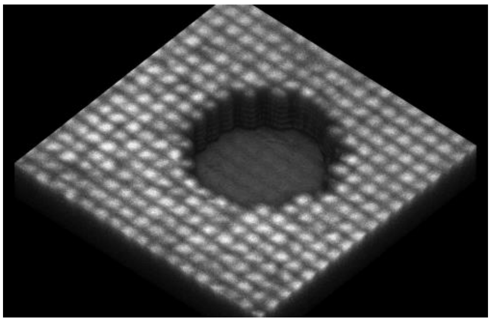 |

| | | | |
|---|---|---|---|
| 29 | Cylinder well | 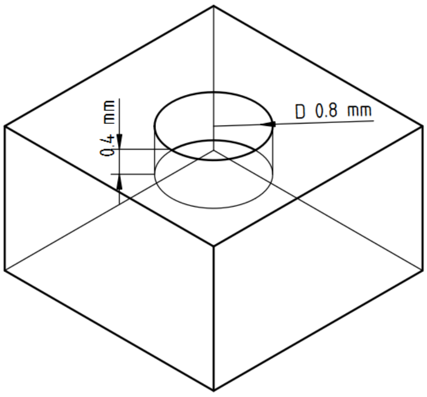  | 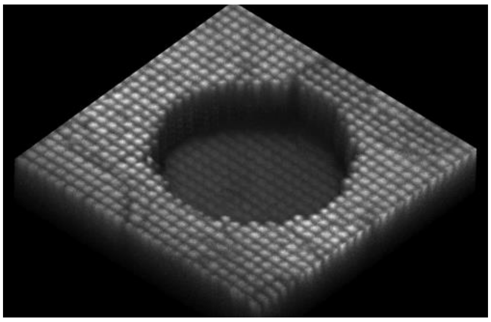 |
| 30 | Cylinder well | 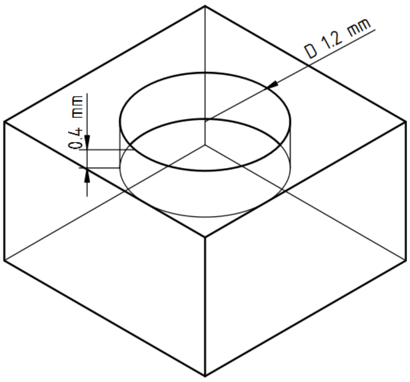  | 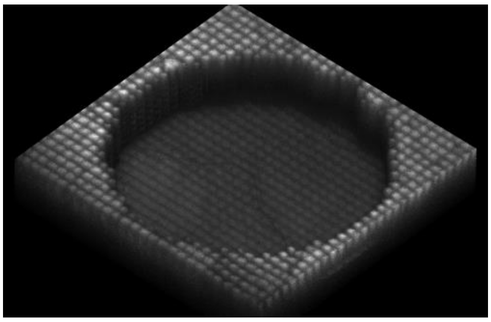 |
| 31 | Cylinder | 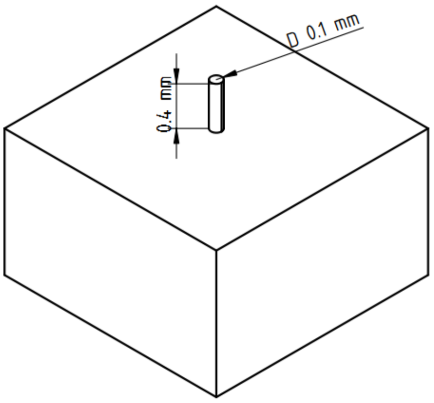  | Printing failure |

| | | | |
|---|---|---|---|
| 32 | Cylinder | 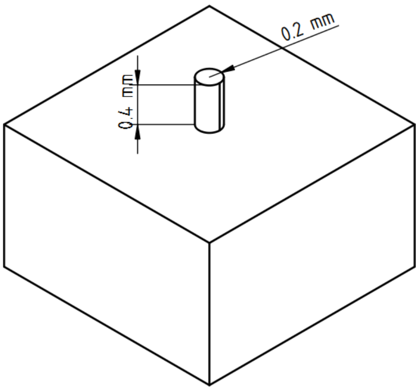  | 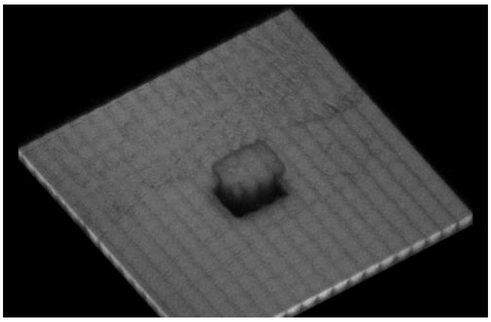 |
| 33 | Cylinder | 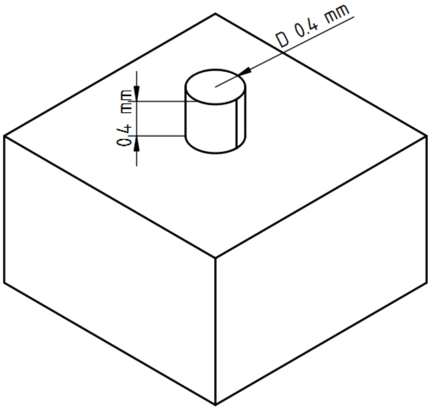  | 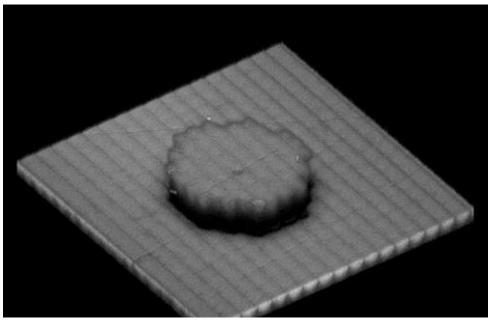 |
| 34 | Cylinder | 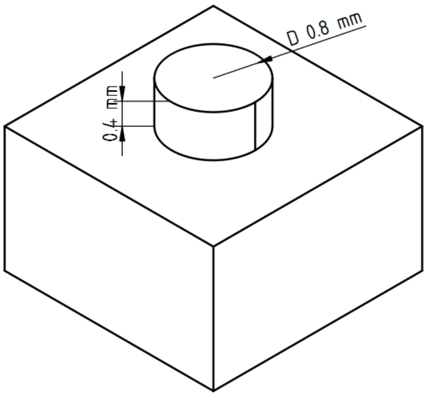  | 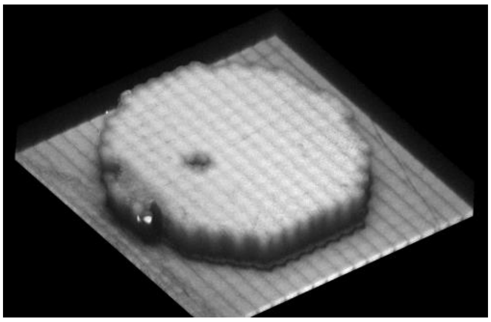 |

| | | | |
|---|---|---|---|
| 35 | Cylinder | 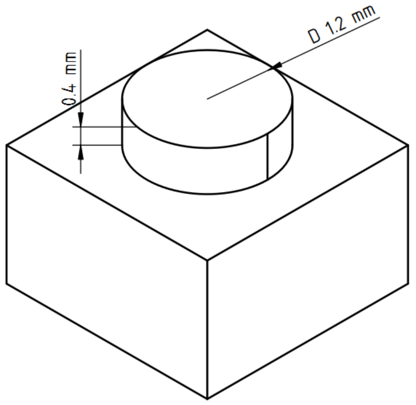 | 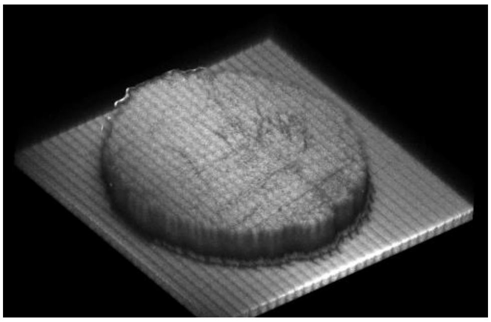 |
| 36 | Cylinder well | 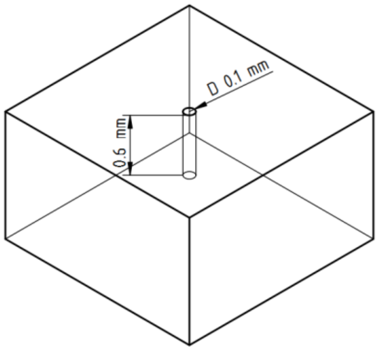 | Printing failure |
| 37 | Cylinder well | 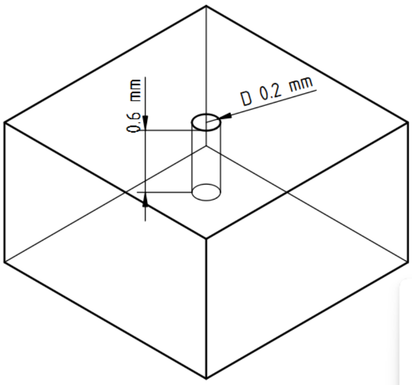 | 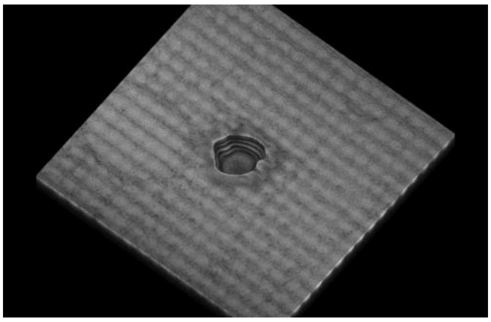 |
| 38 | Cylinder well | 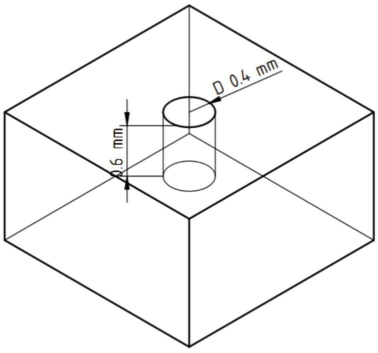 | 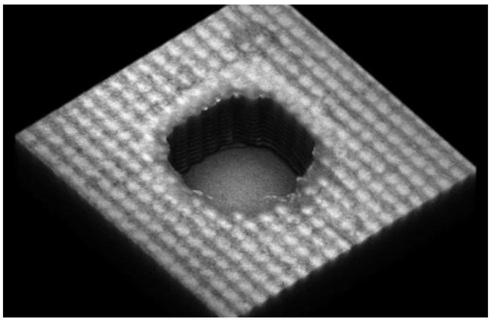 |

| | | | |
|---|---|---|---|
| 39 | Cylinder well | 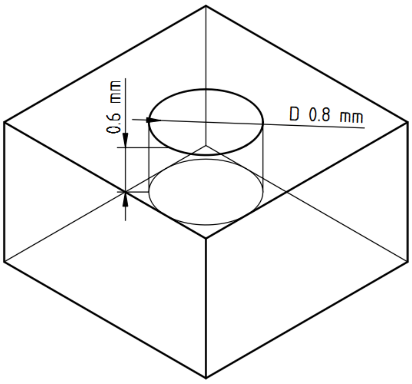 | 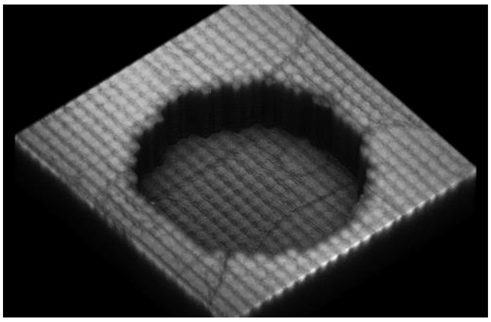 |
| 40 | Cylinder well | 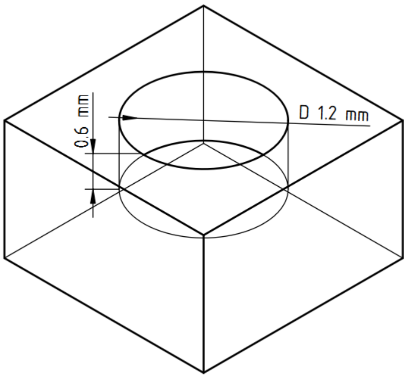 | 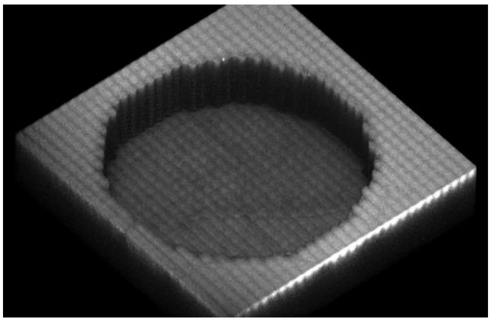 |
| 41 | Cylinder | 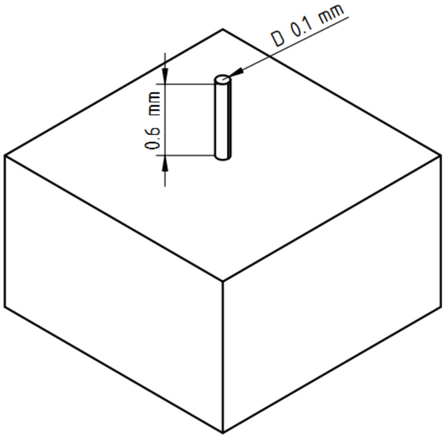 | Printing failure |

| | | | |
|---|---|---|---|
| 42 | Cylinder | 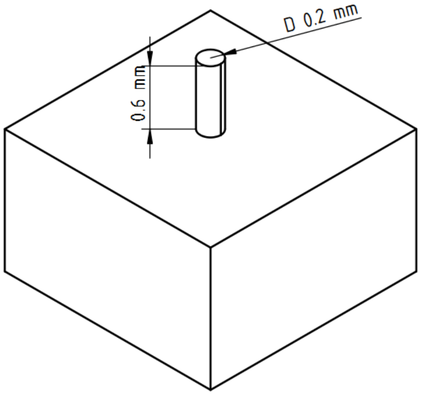  | 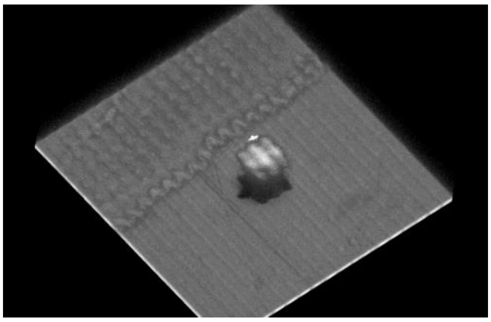 |
| 43 | Cylinder | 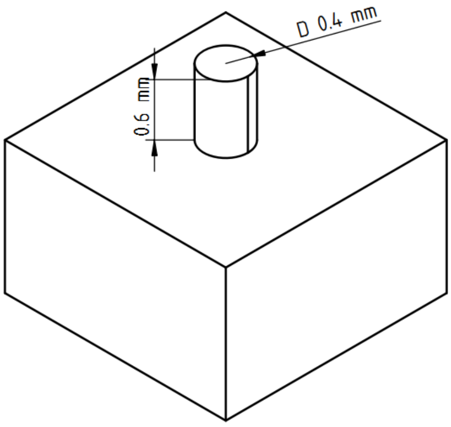  | 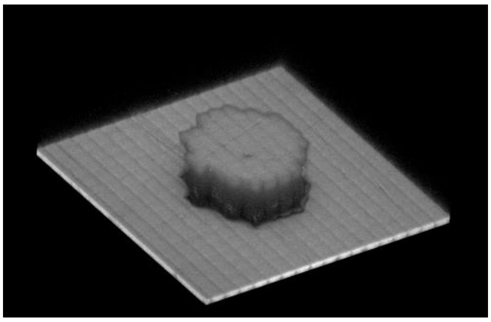 |
| 44 | Cylinder | 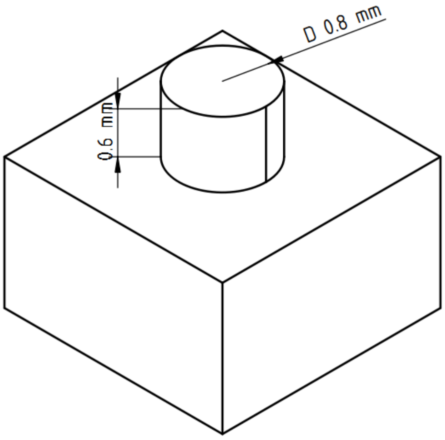  | 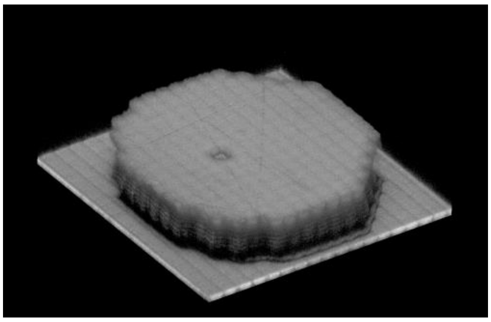 |

| | | | |
|---|---|---|---|
| 45 | Cylinder | 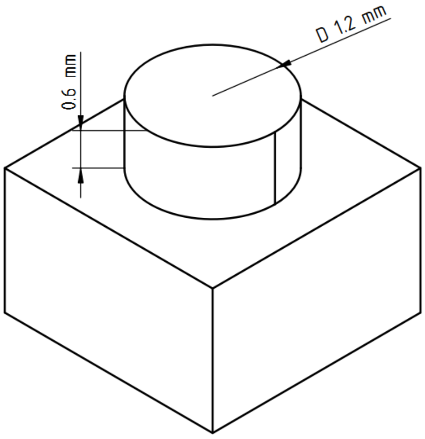 | 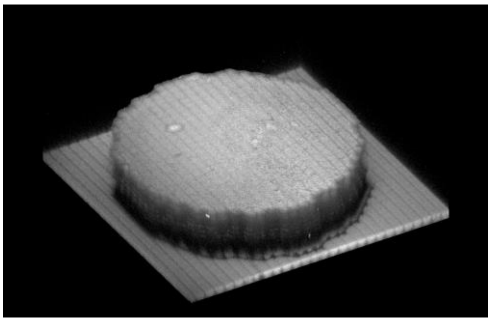 |
| 46 | Cylinder well | 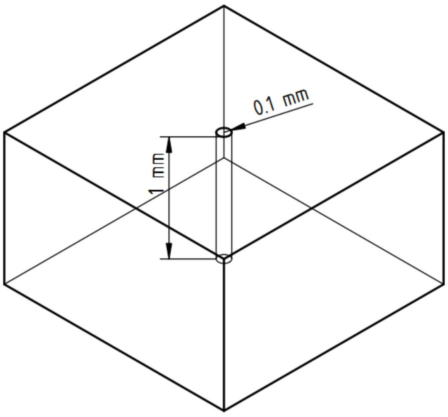 | Printing failure |
| 47 | Cylinder well | 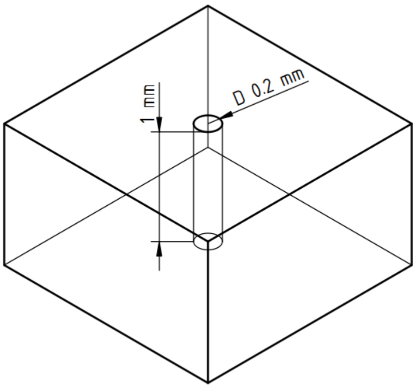 | 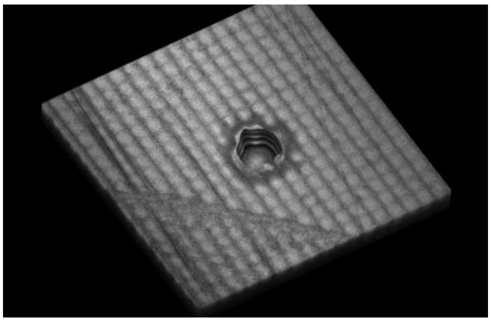 |

| | | | |
|---|---|---|---|
| 48 | Cylinder well | 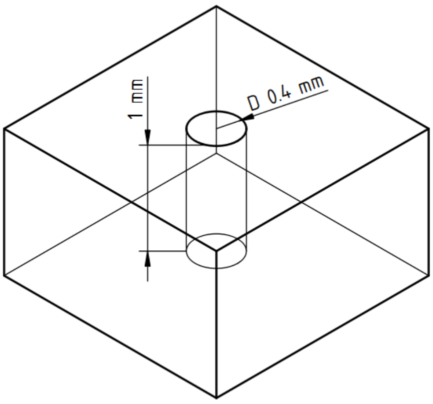  | 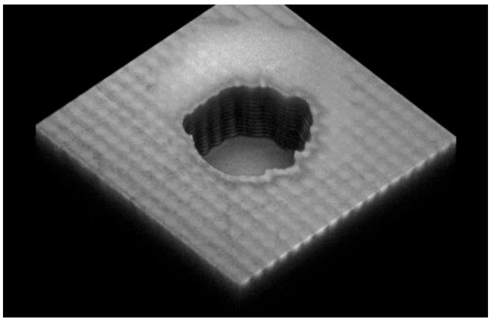 |
| 49 | Cylinder well | 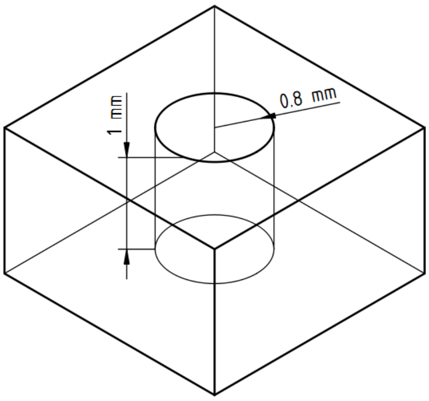  | 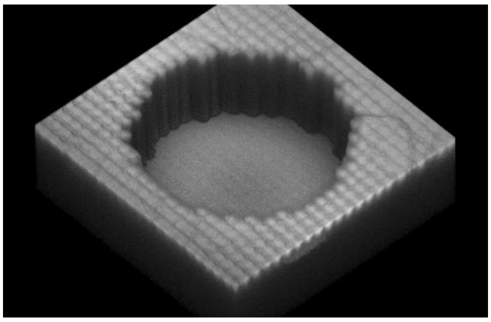 |
| 50 | Cylinder well | 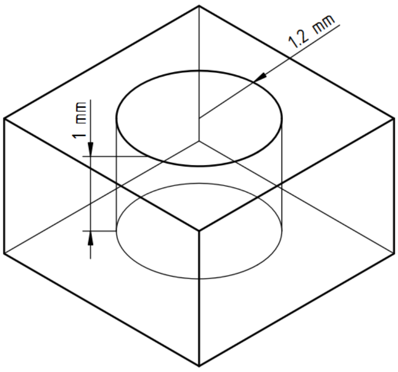  | 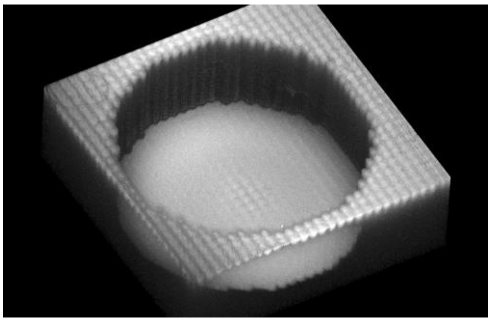 |

| | | | |
|---|---|---|---|
| 51 | Cylinder | 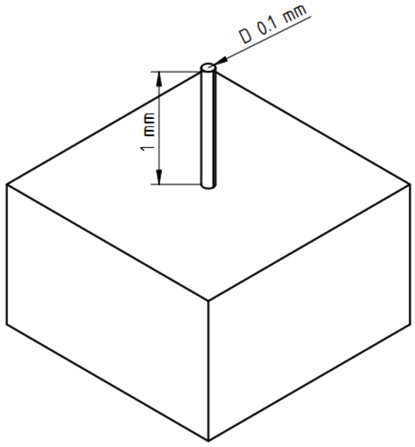  | Printing failure |
| 52 | Cylinder | 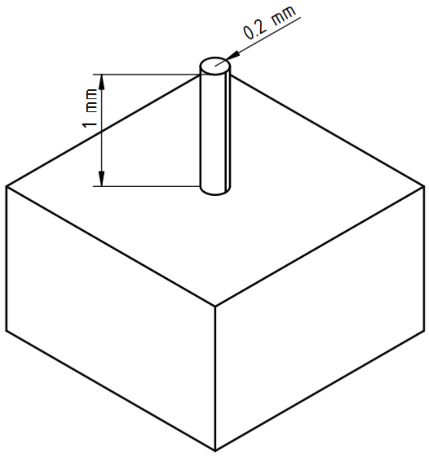  | 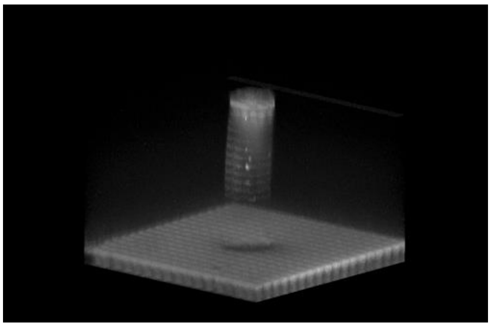 |
| 53 | Cylinder | 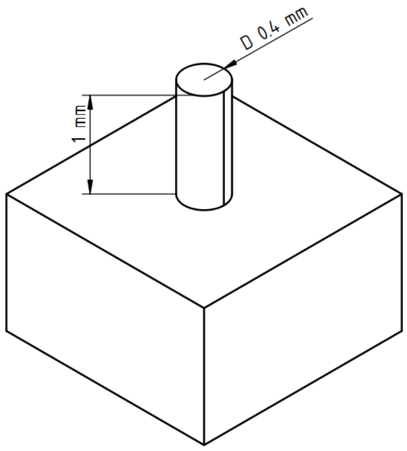  | 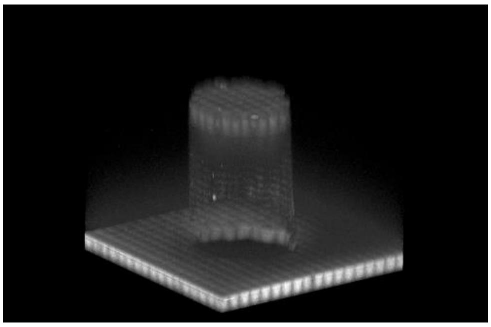 |

<table>
<tr><td>54</td><td>Cylinder</td><td>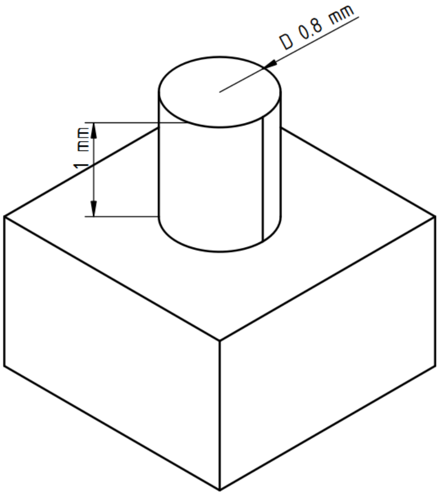
</td><td>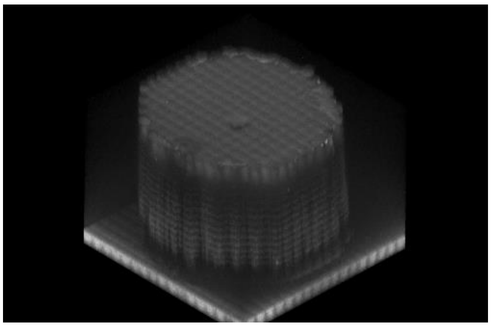</td></tr>
<tr><td>55</td><td>Cylinder</td><td>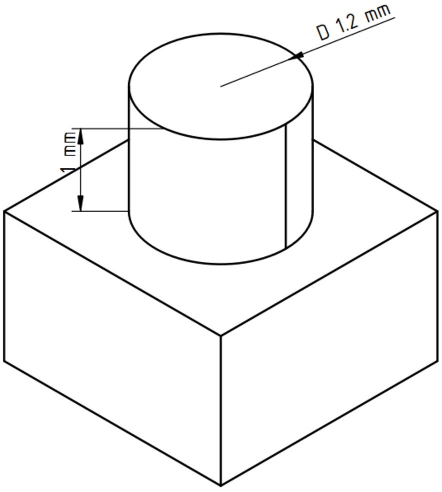
</td><td>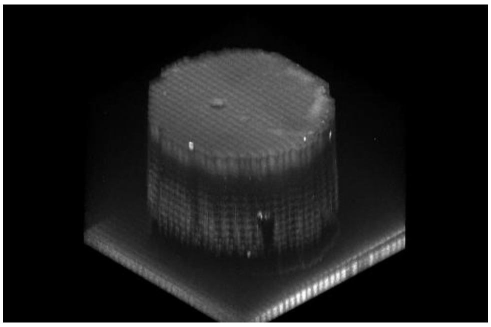</td></tr>
<tr><td>56</td><td>Cone well</td><td>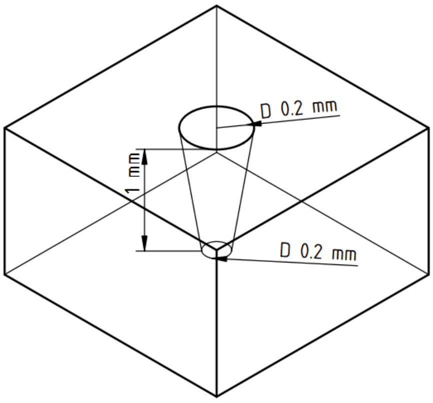
</td><td>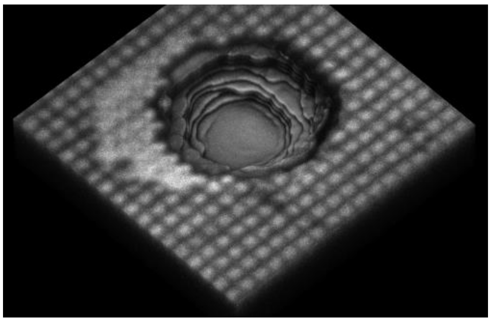</td></tr>
</table>

| | | | |
|---|---|---|---|
| 57 | Cone well | 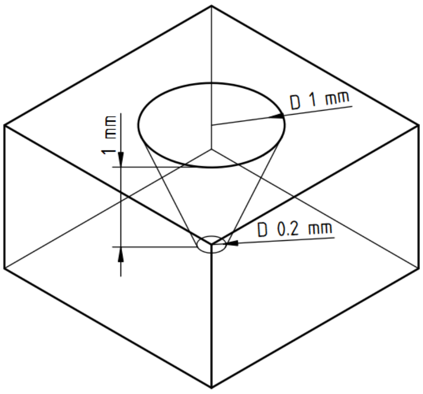  | 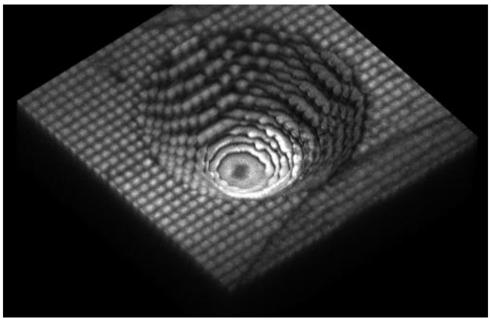 |
| 58 | Cone well | 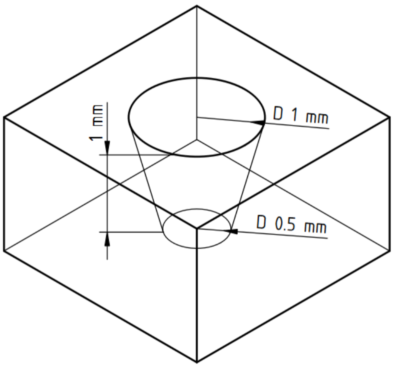  | 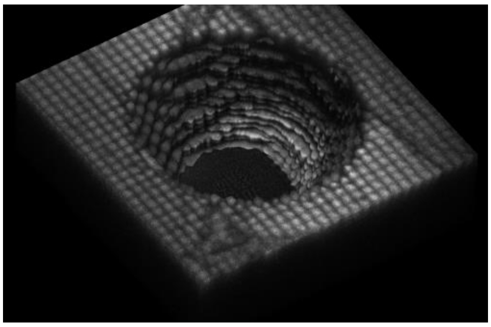 |
| 59 | Cone | 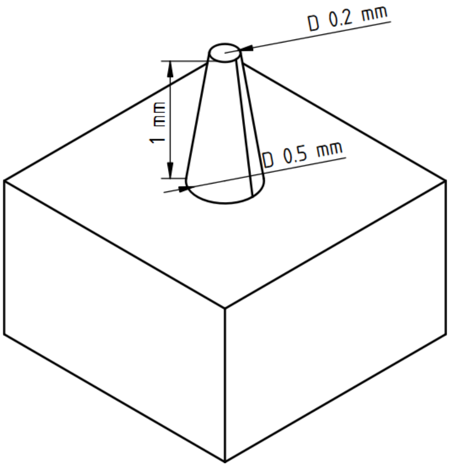  | 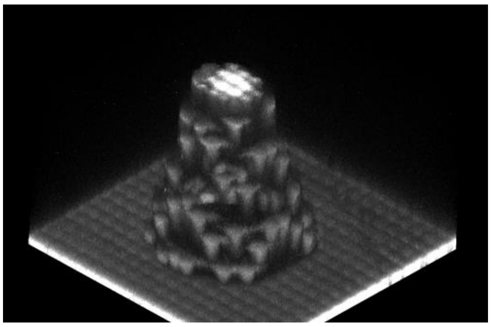 |

| | | | |
|---|---|---|---|
| 60 | Cone | 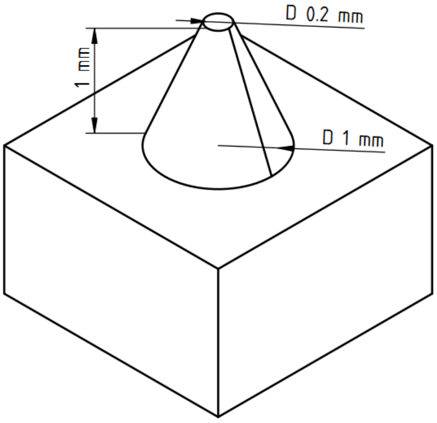  | 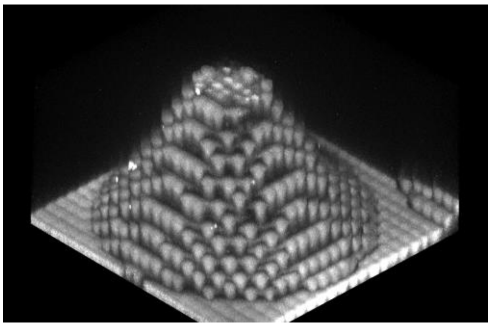 |
| 61 | Cone | 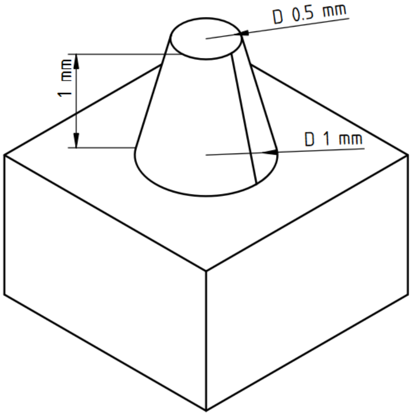  | 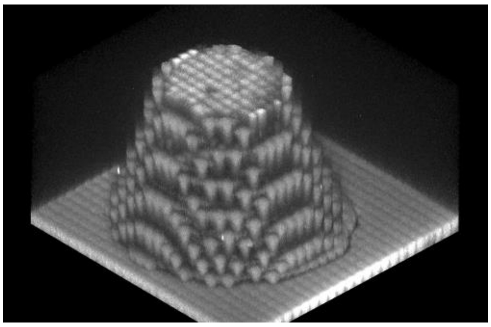 |
| 62 | Cone | 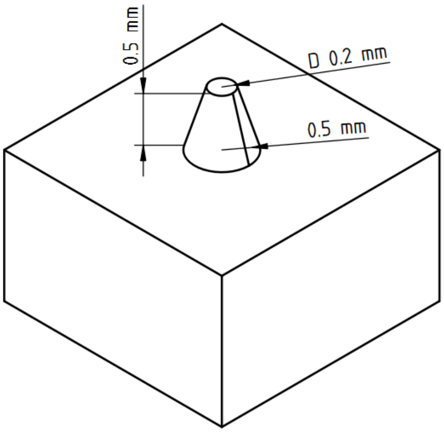  | 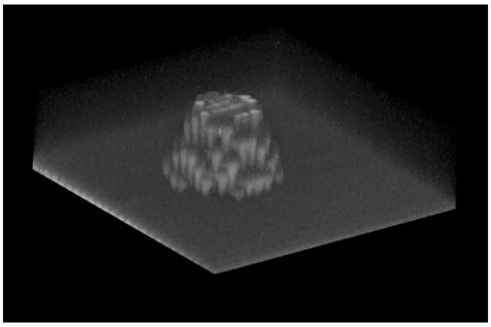 |

| | | | |
|---|---|---|---|
| 63 | Cone | 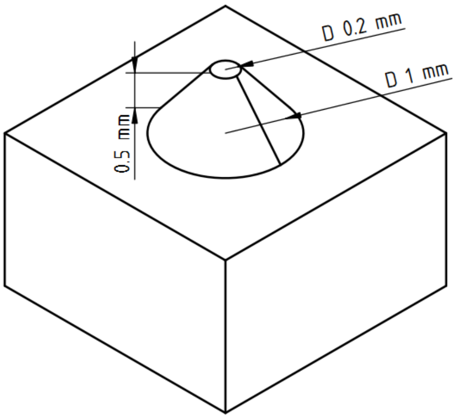 | 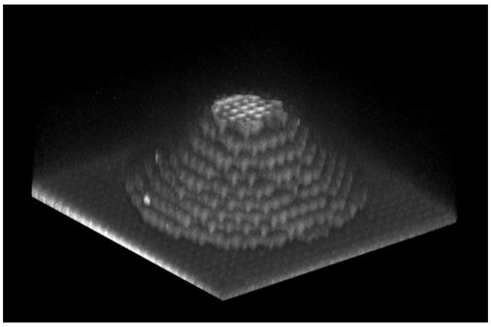 |
| 64 | Cone | 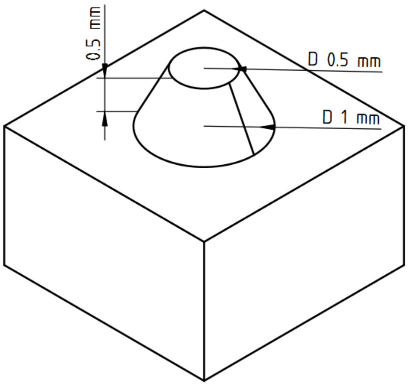 | 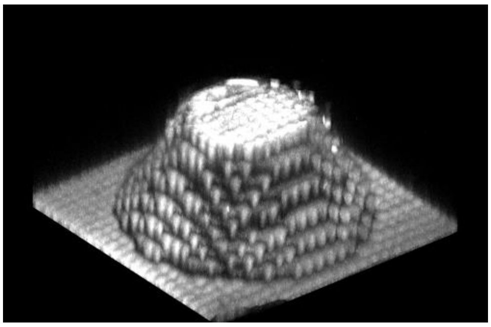 |

### S8. Area distortion

It has already been said that the CAD model is approximated during printing by voxels, which, accordingly, distorts the geometry of the object. However, the question was how strong these distortions are. To check this, test objects with 100 cylinders of different diameters were printed and photographed with a microscope.

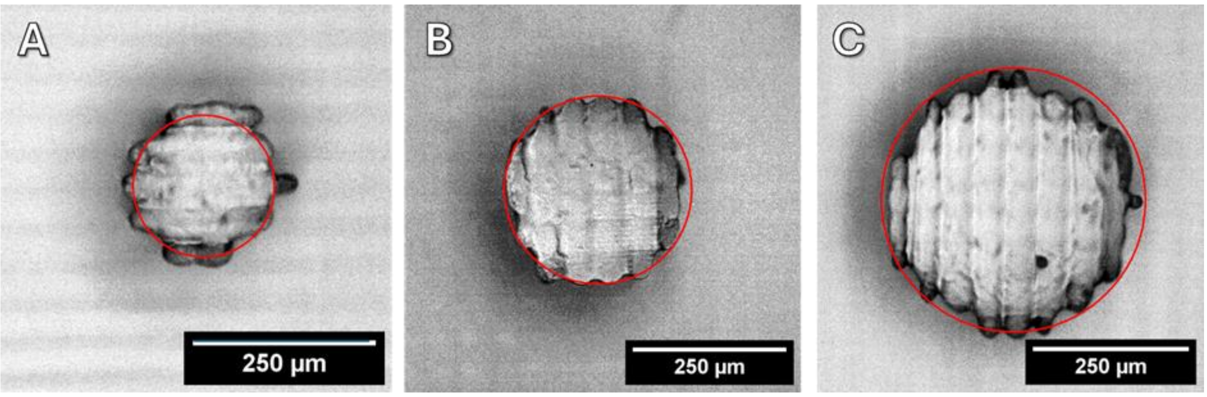

Figure. Cylinders with different diameters. A - 200 µm, B - 300 µm, C - 400 µm. The circle of the corresponding diameter is marked in red.

The microphotographs were further segmented using ilastik and further the geometric dimensions of the objects were measured in the CellProfiler program. The results are summarized in the table and plot below.

| Diameter in CAD model, µm | Measured equivalent diameter, µm |
|---|---|
| 200 | 223 ± 11 |
| 300 | 318 ± 10 |
| 400 | 446 ± 17 |

The presented data show that the obtained diameters are relatively reproducible, the intragroup difference is less than 10%. At the same time, the real diameter of the cylinder slightly differs from the model, but this difference is not so significant if we compare the areas of the objects.

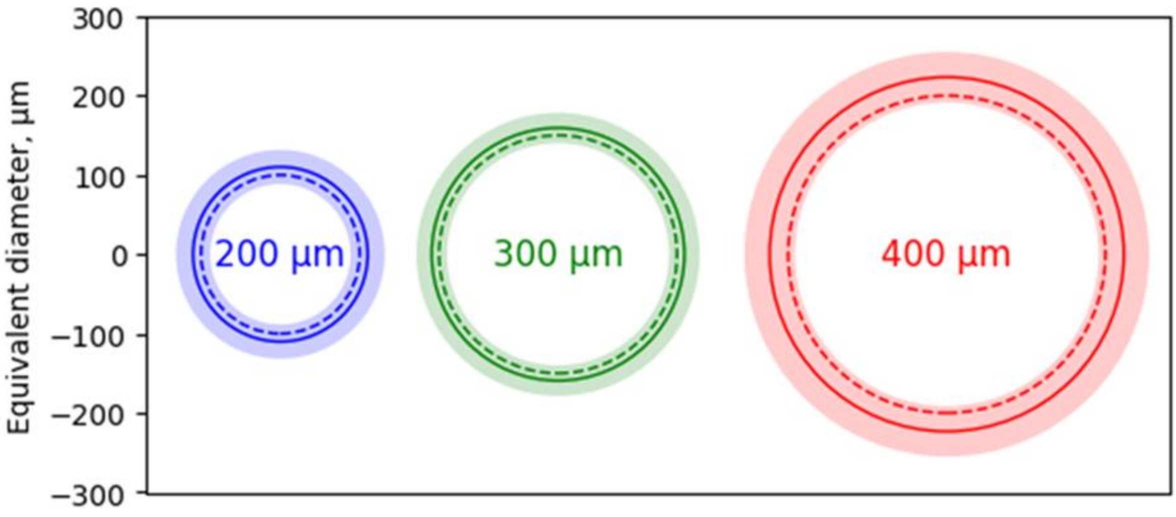


Figure. Graph showing the measured dimensions of the test cylinders. The dashed line is a circle of a given diameter, the solid line is the measured circle of equivalent diameter, the translucent bar is the standard deviation of the measured equivalent diameter.

### S9. Clearing

Using light microscopy, the projected surface area and average fluorescence intensity of the spheroids were measured.

Statistically significant differences were found in each of the characteristics among the fibroblast spheroid groups (Figure 2).

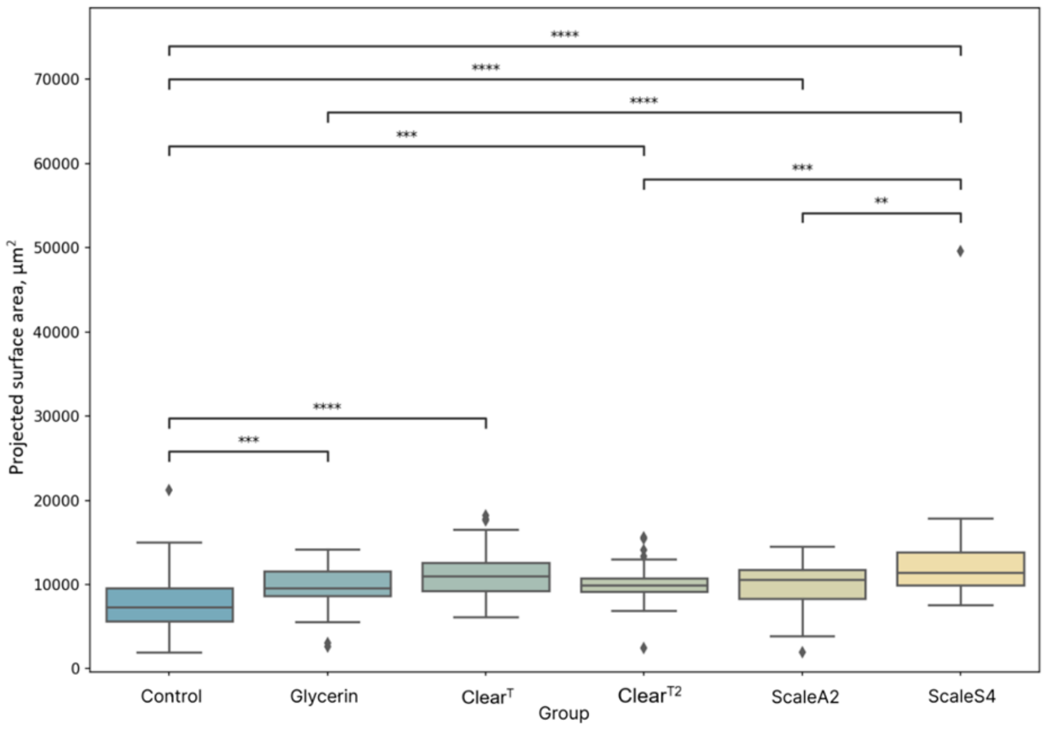


Figure – Projected surface area distribution of fibroblasts spheroids for each optical clearing method. ** - significant difference, 0.001<p<0.01, *** - significant difference, 0.0001<p<0.001; **** - significant difference, p <0.0001. ♦ - outlier

Thus, the projected surface area of the human fibroblast spheroids increased by an average of 37%. These results partially coincide with those published in recent research works [10.1117/1.JBO.23.5.055003]. All authors note that the processing of spheroids and animal tissues causes some increase in size [10.1117/1.JBO.23.5.055003]. Boutin and Hoffman-Kim highlighted that as a result of clearing spheroids according to Scale protocols, the sizes of spheroids in-crease significantly – by up to 40 μm in diameter [10.1089/ten.TEC.2014.0296]. In general, such results are expected, as the main component of Scale solutions, urea, causes tissue hydration.

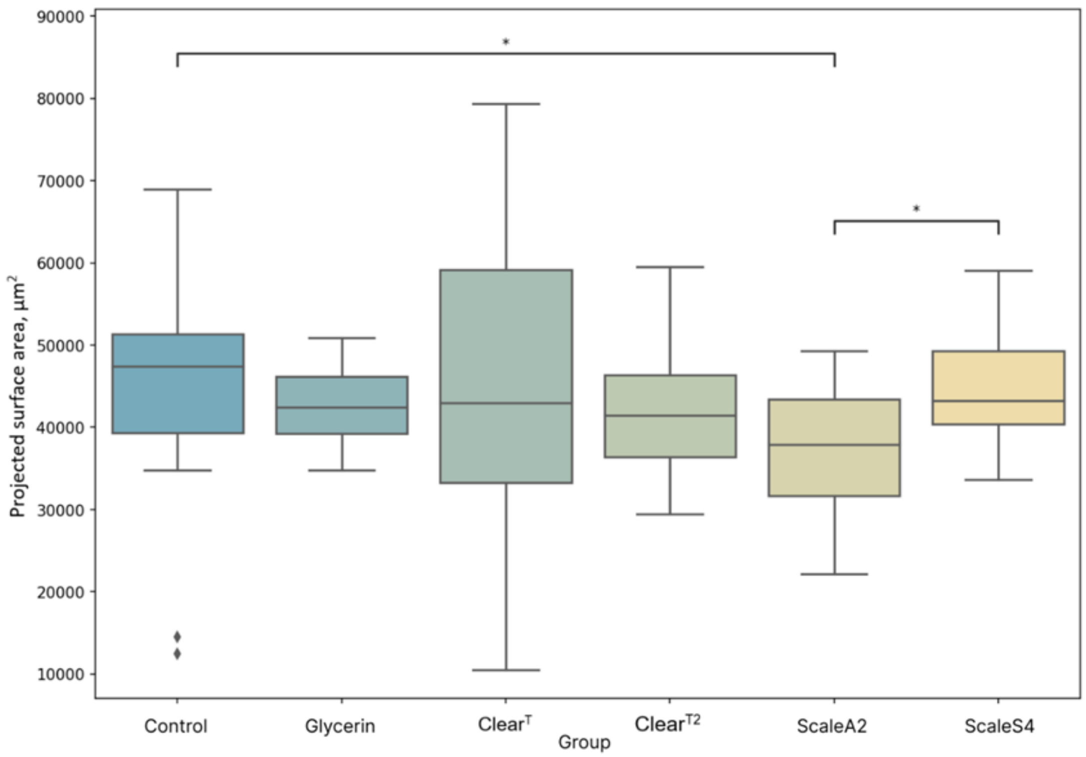


Figure - Projected surface area distribution of rhabdomyosarcoma spheroids for each optical clearing method. * - significant difference, $0.01<p<0.05$. ♦ - outlier

The morphometry results of the spheroids of rhabdomyosarcoma are ambiguous: their projection area has decreased on average by 7.82% (Figure 3). It's not unequivocally clear why the treatment of spheroids of rhabdomyosarcoma with clearing solutions caused a reduction in size. However, rhabdomyosarcoma spheroids have some properties that significantly distinguish them from aggregates of other cultures, they are rather loose, consisting of cells at different stages of differentiation [PMID: 1728419]. Presumably, these features caused atypical changes in morphology after clearing.

The luminous intensity of fibroblast spheroids, or in other words, the transparency in the visible light, of all groups increased by 13.3% on average (Figure 4).

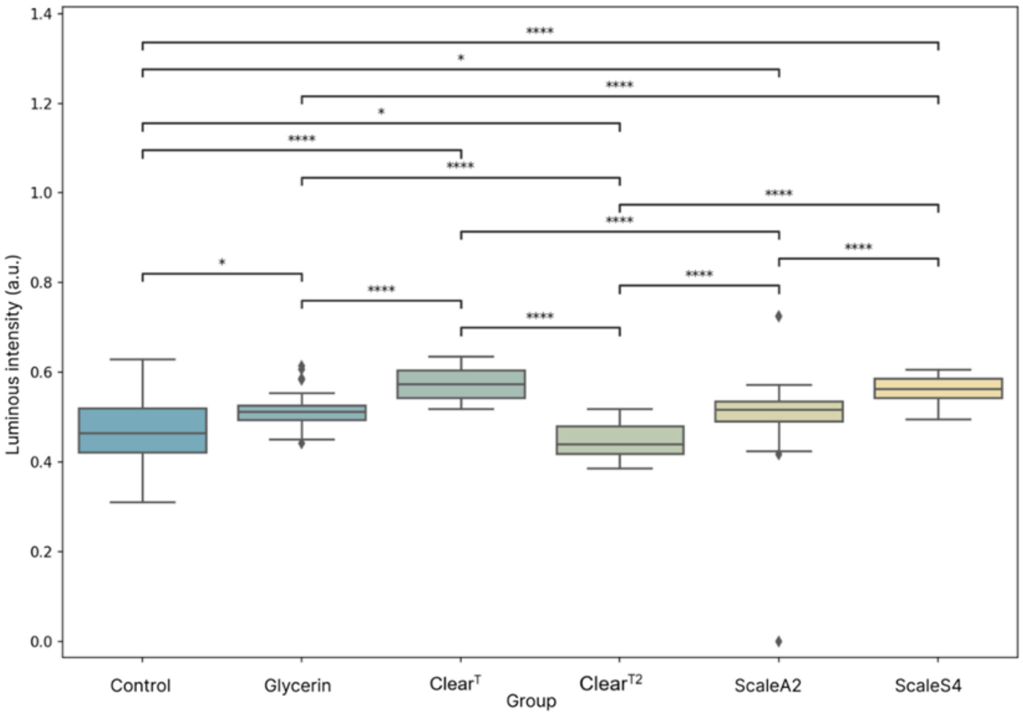


Figure  - Luminous intensity distribution of human fibroblast spheroids for each optical clearing method. * - significant difference, 0.01<p 0.05; **** - significant difference, p<0.0001. ♦ - outlier

The luminous intensity of spheroids of rhabdomyosarcoma cells increased on average by 23.6% after optical clearing (Figure 5). For ClearT spheroids, the transparency slightly decreased (2%).  In this case, although the ClearT group has the highest average transparency of spheroids of human skin fibroblasts, this technique is ineffective for improving transparency in visible light for rhabdomyosarcoma spheroids.

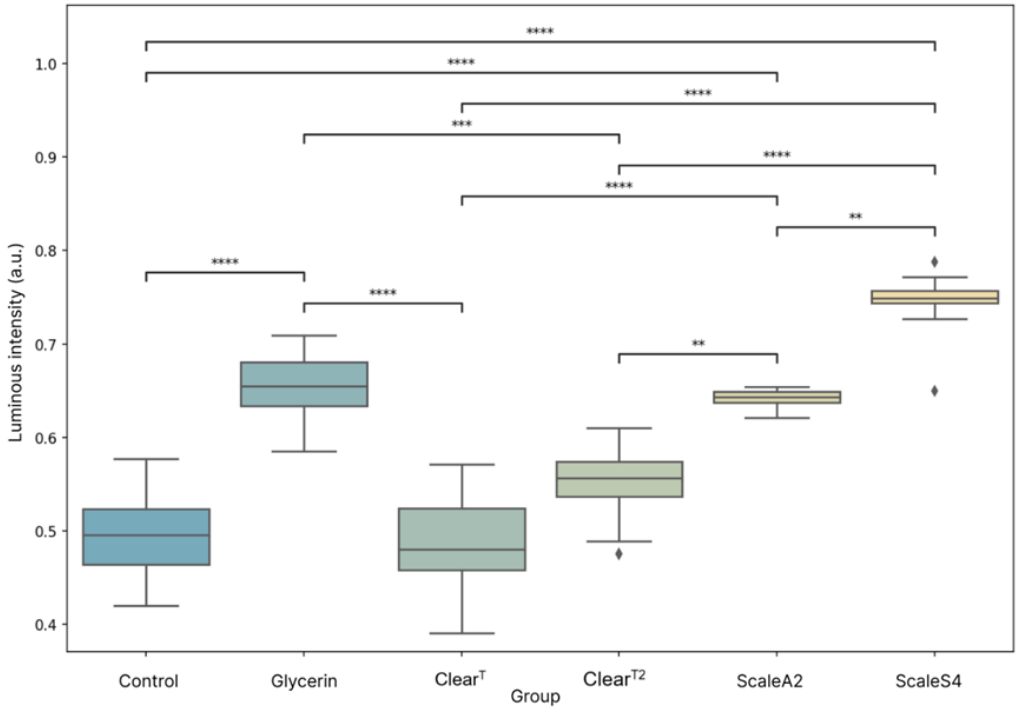

Figure - Luminous intensity distribution of rhabdomyosarcoma spheroids for each optical clearing method. * - significant difference, 0.01<p 0.05; ** - signifi-cant difference, 0.001<p 0.01, *** - significant difference, 0.0001<p 0.001; **** - significant difference, p 0.0001.♦ - outlier

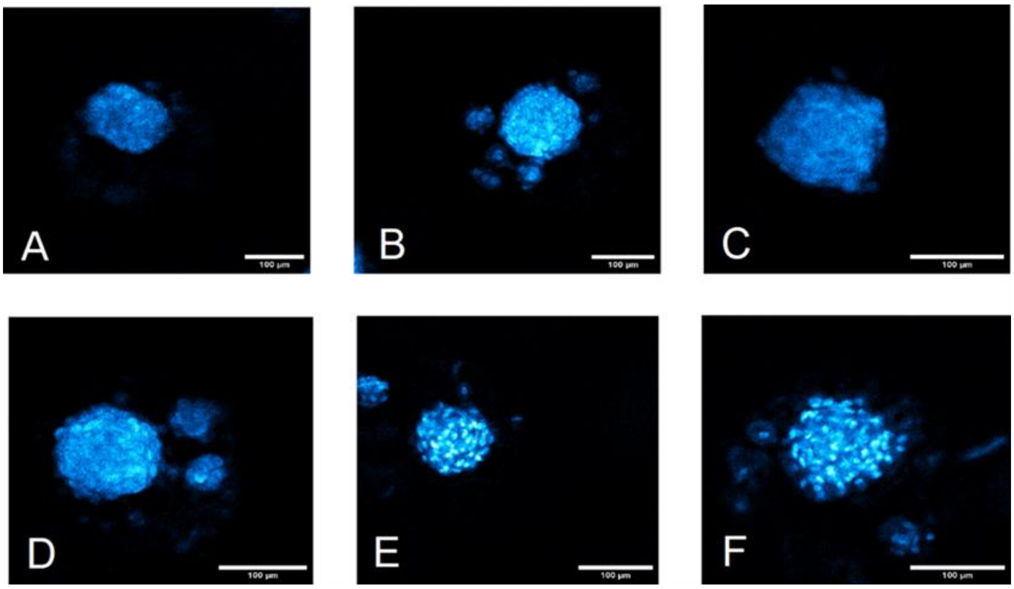


Fluorescent visualization of spheroids of human fibroblasts of different optical cleared groups: A - control, B - glycerol, C - ClearT, D - ClearT2, E - ScaleA2, F - ScaleS4. Hoechst 33342 staining, 10× objective lens.

### S10. Spheroid image processing

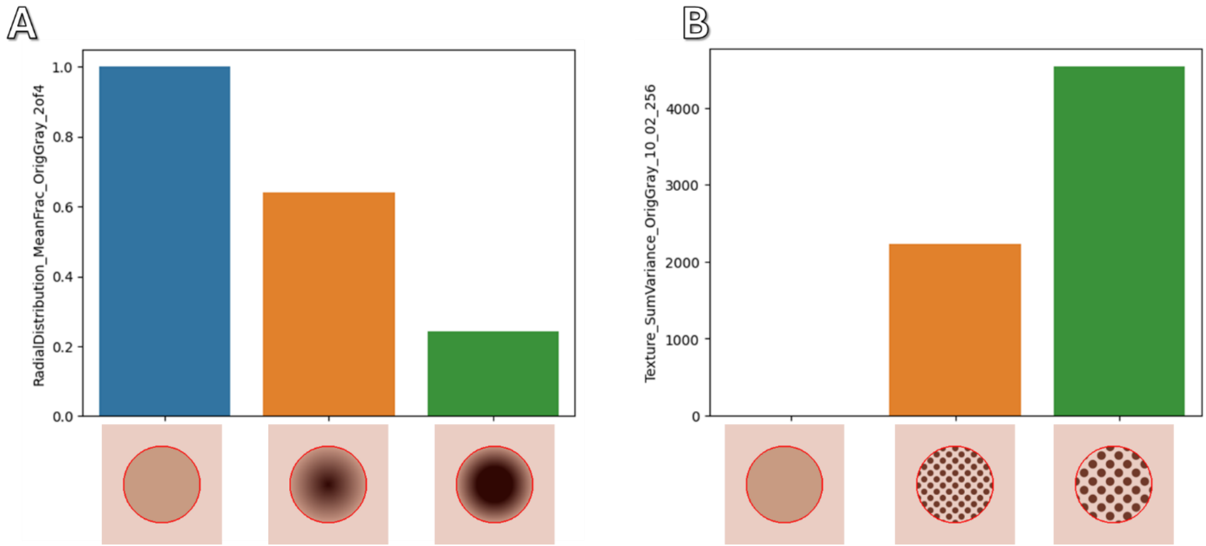


Analysis of radial brightness distribution (A) and texture (B) on synthetic images of spheroids using CellProfiler

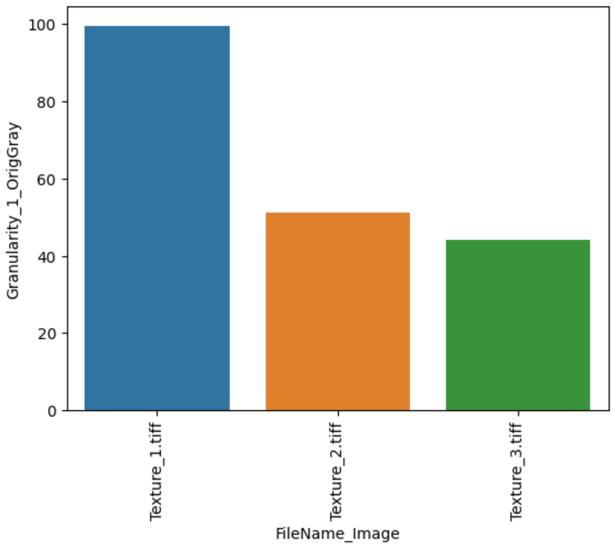


Granularity analysis on synthetic images of spheroids using CellProfiler

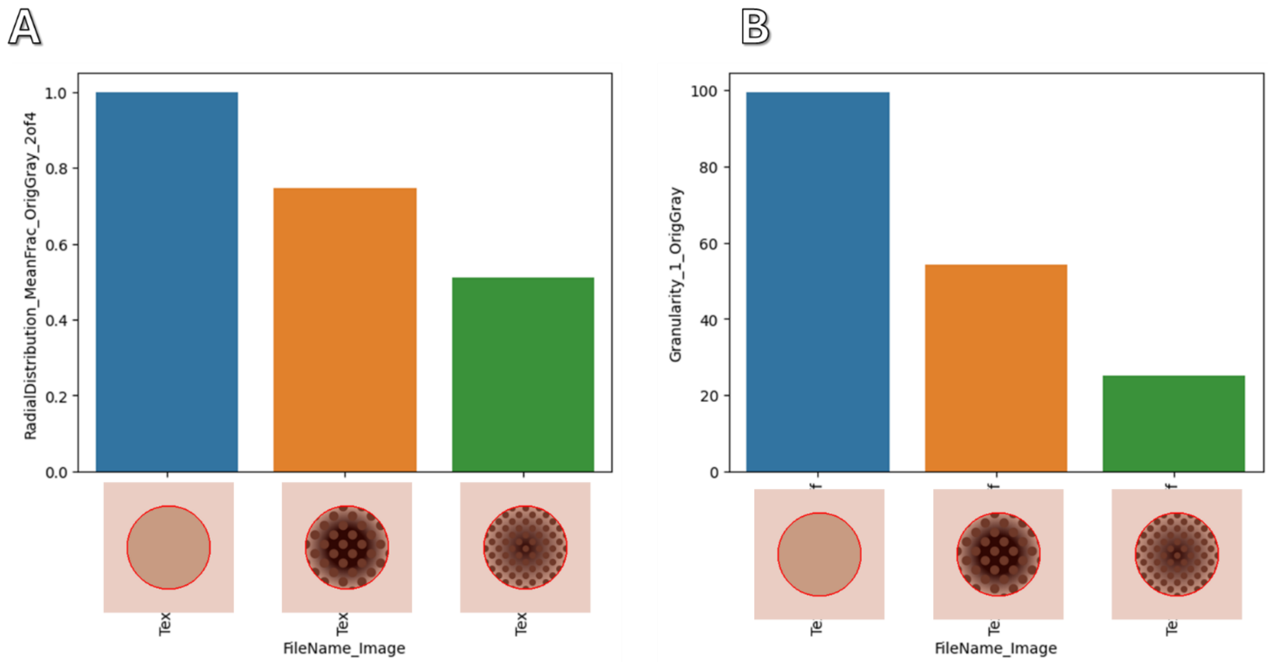


Analysis of radial brightness and granularity distribution in synthetic images of spheroids in which radial variation and optical inhomogeneity (texture) are combined simultaneously using CellProfiler

Table S1. Results of analyzing spheroid image features

| AREA | | | | |
|---|---|---|---|---|
| **Substance/features** | **Class(Time)** | **precision** | **recall** | **f1-score** |
| **Control important features** | | | | |
| AreaShape_Center_Y | | | | |
| AreaShape_BoundingBoxMinimum_Y | | | | |
| AreaShape_BoundingBoxMaximum_Y | | | | |
| AreaShape_BoundingBoxMinimum_X | | | | |

| | | | | |
|---|---|---|---|---|
| AreaShape_Center_X | | | | |
| **Control report** | 0 | 0.86 | 0.75 | 0.80 |
| | 24 | 0.71 | 0.83 | 0.77 |
| | 48 | 1.00 | 1.00 | 1.00 |
| | 72 | 1.00 | 1.00 | 1.00 |
| | accuracy | 0.86 | 0.86 | 0.86 |
| | macro avg | 0.89 | 0.90 | 0.89 |
| | weighted avg | 0.87 | 0.86 | 0.86 |
| **Doxorubicin important features** | | | | |
| AreaShape_MedianRadius | | | | |
| AreaShape_BoundingBoxMaximum_Y | | | | |
| AreaShape_MaximumRadius | | | | |
| AreaShape_Center_Y | | | | |
| AreaShape_Area | | | | |
| **Doxorubicin report** | 0 | 1.00 | 0.50 | 0.67 |
| | 24 | 0.64 | 0.88 | 0.74 |
| | 48 | 0.00 | 0.00 | 0.00 |
| | 72 | 1.00 | 0.80 | 0.89 |
| | accuracy | 0.65 | 0.65 | 0.65 |
| | macro avg | 0.66 | 0.54 | 0.57 |
| | weighted avg | 0.79 | 0.65 | 0.68 |
| **Everolimus important features** | | | | |
| AreaShape_BoundingBoxMinimum_X | | | | |
| AreaShape_BoundingBoxMinimum_Y | | | | |
| AreaShape_Center_X | | | | |
| AreaShape_BoundingBoxMaximum_X | | | | |
| AreaShape_MinFeretDiameter | | | | |
| **Everolimus report** | 0 | 0.67 | 0.29 | 0.40 |
| | 24 | 0.38 | 0.75 | 0.50 |
| | 48 | 0.60 | 0.60 | 0.60 |
| | 72 | 0.80 | 0.80 | 0.80 |
| | accuracy | 0.57 | 0.57 | 0.57 |
| | macro avg | 0.61 | 0.61 | 0.58 |
| | weighted avg | 0.63 | 0.57 | 0.56 |
| **Oxalyplatin important features** | | | | |
| AreaShape_MaximumRadius | | | | |
| AreaShape_MeanRadius | | | | |
| AreaShape_Center_Y | | | | |
| AreaShape_BoundingBoxMinimum_Y | | | | |
| AreaShape_BoundingBoxMaximum_Y | | | | |
| **Oxalyplatin report** | 0 | 1.00 | 0.38 | 0.55 |
| | 24 | 0.45 | 1.00 | 0.63 |
| | 48 | 0.50 | 0.40 | 0.44 |
| | 72 | 1.00 | 1.00 | 1.00 |
| | accuracy | 0.62 | 0.62 | 0.62 |
| | macro avg | 0.74 | 0.69 | 0.65 |

| | | | | |
|---|---|---|---|---|
| | weighted avg | 0.75 | 0.62 | 0.61 |
| **Paclitaxel important features** | | | | |
| AreaShape_BoundingBoxMaximum_Y | | | | |
| AreaShape_BoundingBoxArea | | | | |
| AreaShape_Area | | | | |
| AreaShape_EquivalentDiameter | | | | |
| AreaShape_MedianRadius | | | | |
| **Paclitaxel report** | 0 | 0.89 | 0.89 | 0.89 |
| | 24 | 1.00 | 1.00 | 1.00 |
| | 48 | 0.67 | 1.00 | 0.80 |
| | 72 | 1.00 | 0.75 | 0.86 |
| | accuracy | 0.90 | 0.90 | 0.90 |
| | macro avg | 0.89 | 0.91 | 0.89 |
| | weighted avg | 0.92 | 0.90 | 0.90 |
| **Topotecan important features** | | | | |
| AreaShape_BoundingBoxMaximum_Y | | | | |
| AreaShape_MedianRadius | | | | |
| AreaShape_Center_Y | | | | |
| AreaShape_BoundingBoxArea | | | | |
| AreaShape_Area | | | | |
| **Topotecan report** | 0 | 0.80 | 0.50 | 0.62 |
| | 24 | 0.50 | 0.60 | 0.55 |
| | 48 | 0.50 | 0.67 | 0.57 |
| | 72 | 1.00 | 1.00 | 1.00 |
| | accuracy | 0.62 | 0.62 | 0.62 |
| | macro avg | 0.70 | 0.69 | 0.68 |
| | weighted avg | 0.66 | 0.62 | 0.62 |
| **Docetaxel important features** | | | | |
| AreaShape_Center_X | | | | |
| AreaShape_BoundingBoxMinimum_X | | | | |
| AreaShape_BoundingBoxMaximum_X | | | | |
| AreaShape_Area | | | | |
| AreaShape_EquivalentDiameter | | | | |
| **Docetaxel report** | 0 | 0.83 | 0.63 | 0.71 |
| | 24 | 0.71 | 0.83 | 0.77 |
| | 48 | 0.50 | 0.75 | 0.60 |
| | 72 | 1.00 | 0.75 | 0.86 |
| | accuracy | 0.73 | 0.73 | 0.73 |
| | macro avg | 0.76 | 0.74 | 0.74 |
| | weighted avg | 0.77 | 0.73 | 0.73 |
| **DMSO_30pc important features** | | | | |
| AreaShape_BoundingBoxMinimum_X | | | | |
| AreaShape_MaximumRadius | | | | |
| AreaShape_BoundingBoxMaximum_Y | | | | |
| AreaShape_MinorAxisLength | | | | |
| AreaShape_MedianRadius | | | | |

| | | | | |
|---|---|---|---|---|
| **DMSO_30pc report** | 0 | 0.80 | 0.50 | 0.62 |
| | 24 | 0.67 | 0.33 | 0.44 |
| | 48 | 0.22 | 0.50 | 0.31 |
| | 72 | 0.60 | 0.75 | 0.67 |
| | accuracy | 0.50 | 0.50 | 0.50 |
| | macro avg | 0.57 | 0.52 | 0.51 |
| | weighted avg | 0.62 | 0.50 | 0.52 |
| **Campthotecin important features** | | | | |
| AreaShape_BoundingBoxMaximum_Y | | | | |
| AreaShape_Center_Y | | | | |
| AreaShape_MedianRadius | | | | |
| AreaShape_MeanRadius | | | | |
| AreaShape_MaximumRadius | | | | |
| **Campthotecin report** | 0 | 1.00 | 0.63 | 0.77 |
| | 24 | 0.67 | 0.86 | 0.75 |
| | 48 | 0.50 | 0.75 | 0.60 |
| | 72 | 1.00 | 0.75 | 0.86 |
| | accuracy | 0.74 | 0.74 | 0.74 |
| | macro avg | 0.79 | 0.75 | 0.74 |
| | weighted avg | 0.81 | 0.74 | 0.75 |
| **RADIAL** | | | | |
| **Substance/features** | **Class(Time)** | **precision** | **recall** | **f1-score** |
| **Control important features** | | | | |
| RadialDistribution_MeanFrac_OrigGray_2of4 | | | | |
| RadialDistribution_FracAtD_OrigGray_2of4 | | | | |
| RadialDistribution_MeanFrac_OrigGray_1of4 | | | | |
| RadialDistribution_FracAtD_OrigGray_1of4 | | | | |
| RadialDistribution_MeanFrac_OrigGray_4of4 | | | | |
| **Control report** | 0 | 1.00 | 1.00 | 1.00 |
| | 24 | 1.00 | 1.00 | 1.00 |
| | 48 | 1.00 | 0.67 | 0.80 |
| | 72 | 0.50 | 1.00 | 0.67 |
| | accuracy | 0.91 | 0.91 | 0.91 |
| | macro avg | 0.88 | 0.92 | 0.87 |
| | weighted avg | 0.95 | 0.91 | 0.92 |
| **Doxorubicin important features** | | | | |
| RadialDistribution_FracAtD_OrigGray_2of4 | | | | |
| RadialDistribution_MeanFrac_OrigGray_2of4 | | | | |
| RadialDistribution_FracAtD_OrigGray_4of4 | | | | |
| RadialDistribution_MeanFrac_OrigGray_1of4 | | | | |
| RadialDistribution_FracAtD_OrigGray_1of4 | | | | |
| **Doxorubicin report** | 0 | 1.00 | 1.00 | 1.00 |
| | 24 | 0.86 | 0.75 | 0.80 |
| | 48 | 0.25 | 0.50 | 0.33 |
| | 72 | 0.25 | 0.20 | 0.22 |
| | accuracy | 0.70 | 0.70 | 0.70 |

| | | | | |
|---|---|---|---|---|
| | macro avg | 0.59 | 0.61 | 0.59 |
| | weighted avg | 0.72 | 0.70 | 0.70 |
| **Everolimus important features** | | | | |
| RadialDistribution_FracAtD_OrigGray_1of4 | | | | |
| RadialDistribution_FracAtD_OrigGray_4of4 | | | | |
| RadialDistribution_FracAtD_OrigGray_2of4 | | | | |
| RadialDistribution_MeanFrac_OrigGray_3of4 | | | | |
| RadialDistribution_FracAtD_OrigGray_3of4 | | | | |
| **Everolimus report** | 0 | 1.00 | 1.00 | 1.00 |
| | 24 | 0.29 | 1.00 | 0.44 |
| | 48 | 0.00 | 0.00 | 0.00 |
| | 72 | 0.00 | 0.00 | 0.00 |
| | accuracy | 0.52 | 0.52 | 0.52 |
| | macro avg | 0.32 | 0.50 | 0.36 |
| | weighted avg | 0.39 | 0.52 | 0.42 |
| **Oxalyplatin important features** | | | | |
| RadialDistribution_RadialCV_OrigGray_1of4 | | | | |
| RadialDistribution_MeanFrac_OrigGray_4of4 | | | | |
| RadialDistribution_MeanFrac_OrigGray_1of4 | | | | |
| RadialDistribution_MeanFrac_OrigGray_2of4 | | | | |
| RadialDistribution_ZernikeMagnitude_OrigGray_0_0 | | | | |
| **Oxalyplatin report** | 0 | 0.89 | 1.00 | 0.94 |
| | 24 | 0.67 | 0.80 | 0.73 |
| | 48 | 1.00 | 0.60 | 0.75 |
| | 72 | 0.33 | 0.33 | 0.33 |
| | accuracy | 0.76 | 0.76 | 0.76 |
| | macro avg | 0.72 | 0.68 | 0.69 |
| | weighted avg | 0.78 | 0.76 | 0.76 |
| **Paclitaxel important features** | | | | |
| RadialDistribution_FracAtD_OrigGray_2of4 | | | | |
| RadialDistribution_MeanFrac_OrigGray_2of4 | | | | |
| RadialDistribution_RadialCV_OrigGray_4of4 | | | | |
| RadialDistribution_FracAtD_OrigGray_4of4 | | | | |
| RadialDistribution_FracAtD_OrigGray_3of4 | | | | |
| | 0 | 1.00 | 1.00 | 1.00 |
| | 24 | 0.56 | 1.00 | 0.71 |
| | 48 | 0.50 | 0.50 | 0.50 |
| | 72 | 0.00 | 0.00 | 0.00 |
| | accuracy | 0.75 | 0.75 | 0.75 |
| | macro avg | 0.51 | 0.63 | 0.55 |
| | weighted avg | 0.64 | 0.75 | 0.68 |
| **Topotecan important features** | | | | |
| RadialDistribution_FracAtD_OrigGray_2of4 | | | | |
| RadialDistribution_MeanFrac_OrigGray_4of4 | | | | |
| RadialDistribution_MeanFrac_OrigGray_1of4 | | | | |
| RadialDistribution_MeanFrac_OrigGray_2of4 | | | | |

| | | | | |
|---|---|---|---|---|
| RadialDistribution_MeanFrac_OrigGray_3of4 | | | | |
| **Topotecan report** | 0 | 1.00 | 1.00 | 1.00 |
| | 24 | 0.40 | 0.80 | 0.53 |
| | 48 | 0.00 | 0.00 | 0.00 |
| | 72 | 1.00 | 0.50 | 0.67 |
| | accuracy | 0.62 | 0.62 | 0.62 |
| | macro avg | 0.60 | 0.58 | 0.55 |
| | weighted avg | 0.57 | 0.62 | 0.57 |
| **Docetaxel important features** | | | | |
| RadialDistribution_MeanFrac_OrigGray_4of4 | | | | |
| RadialDistribution_MeanFrac_OrigGray_3of4 | | | | |
| RadialDistribution_MeanFrac_OrigGray_2of4 | | | | |
| RadialDistribution_MeanFrac_OrigGray_1of4 | | | | |
| RadialDistribution_RadialCV_OrigGray_1of4 | | | | |
| **Docetaxel report** | 0 | 1.00 | 1.00 | 1.00 |
| | 24 | 0.63 | 0.83 | 0.71 |
| | 48 | 0.00 | 0.00 | 0.00 |
| | 72 | 0.33 | 0.50 | 0.40 |
| | accuracy | 0.68 | 0.68 | 0.68 |
| | macro avg | 0.49 | 0.58 | 0.53 |
| | weighted avg | 0.59 | 0.68 | 0.63 |
| **DMSO_30pc important features** | | | | |
| RadialDistribution_FracAtD_OrigGray_2of4 | | | | |
| RadialDistribution_FracAtD_OrigGray_3of4 | | | | |
| RadialDistribution_FracAtD_OrigGray_4of4 | | | | |
| RadialDistribution_ZernikeMagnitude_OrigGray_4_0 | | | | |
| RadialDistribution_FracAtD_OrigGray_1of4 | | | | |
| **DMSO_30pc report** | 0 | 0.89 | 1.00 | 0.94 |
| | 24 | 0.33 | 0.33 | 0.33 |
| | 48 | 0.17 | 0.25 | 0.20 |
| | 72 | 0.00 | 0.00 | 0.00 |
| | accuracy | 0.50 | 0.50 | 0.50 |
| | macro avg | 0.35 | 0.40 | 0.37 |
| | weighted avg | 0.44 | 0.50 | 0.47 |
| **Campthotecin important features** | | | | |
| RadialDistribution_FracAtD_OrigGray_2of4 | | | | |
| RadialDistribution_FracAtD_OrigGray_3of4 | | | | |
| RadialDistribution_FracAtD_OrigGray_1of4 | | | | |
| RadialDistribution_MeanFrac_OrigGray_3of4 | | | | |
| RadialDistribution_MeanFrac_OrigGray_2of4 | | | | |
| **Campthotecin report** | 0 | 1.00 | 1.00 | 1.00 |
| | 24 | 0.86 | 0.86 | 0.86 |
| | 48 | 0.33 | 0.25 | 0.29 |
| | 72 | 0.40 | 0.50 | 0.44 |
| | accuracy | 0.74 | 0.74 | 0.74 |
| | macro avg | 0.65 | 0.65 | 0.65 |

| | | | | |
|---|---|---|---|---|
| | weighted avg | 0.74 | 0.74 | 0.74 |
| **GRANULARITY** | | | | |
| **Substance/features** | **Class(Time)** | **precision** | **recall** | **f1-score** |
| **Control important features** | | | | |
| Granularity_2_OrigGray | | | | |
| Granularity_1_OrigGray | | | | |
| Granularity_3_OrigGray | | | | |
| Granularity_7_OrigGray | | | | |
| Granularity_4_OrigGray | | | | |
| **Control report** | 0 | 0.50 | 0.25 | 0.33 |
| | 24 | 0.57 | 0.67 | 0.62 |
| | 48 | 0.25 | 0.17 | 0.20 |
| | 72 | 0.14 | 0.50 | 0.22 |
| | accuracy | 0.36 | 0.36 | 0.36 |
| | macro avg | 0.37 | 0.40 | 0.34 |
| | weighted avg | 0.42 | 0.36 | 0.36 |
| **Doxorubicin important features** | | | | |
| Granularity_1_OrigGray | | | | |
| Granularity_2_OrigGray | | | | |
| Granularity_10_OrigGray | | | | |
| Granularity_7_OrigGray | | | | |
| Granularity_3_OrigGray | | | | |
| **Doxorubicin report** | 0 | 0.83 | 0.63 | 0.71 |
| | 24 | 0.80 | 0.50 | 0.62 |
| | 48 | 0.20 | 0.50 | 0.29 |
| | 72 | 0.43 | 0.60 | 0.50 |
| | accuracy | 0.57 | 0.57 | 0.57 |
| | macro avg | 0.57 | 0.56 | 0.53 |
| | weighted avg | 0.68 | 0.57 | 0.60 |
| **Everolimus important features** | | | | |
| Granularity_1_OrigGray | | | | |
| Granularity_10_OrigGray | | | | |
| Granularity_2_OrigGray | | | | |
| Granularity_6_OrigGray | | | | |
| Granularity_7_OrigGray | | | | |
| **Everolimus report** | 0 | 0.83 | 0.71 | 0.77 |
| | 24 | 0.27 | 0.75 | 0.40 |
| | 48 | 0.67 | 0.40 | 0.50 |
| | 72 | 0.00 | 0.00 | 0.00 |
| | accuracy | 0.48 | 0.48 | 0.48 |
| | macro avg | 0.44 | 0.47 | 0.42 |
| | weighted avg | 0.49 | 0.48 | 0.45 |
| **Oxalyplatin important features** | | | | |
| Granularity_2_OrigGray | | | | |
| Granularity_3_OrigGray | | | | |
| Granularity_1_OrigGray | | | | |

| | | | | |
|---|---|---|---|---|
| Granularity_5_OrigGray | | | | |
| Granularity_4_OrigGray | | | | |
| **Oxalyplatin report** | 0 | 0.80 | 0.50 | 0.62 |
| | 24 | 0.67 | 0.80 | 0.73 |
| | 48 | 0.43 | 0.60 | 0.50 |
| | 72 | 0.00 | 0.00 | 0.00 |
| | accuracy | 0.52 | 0.52 | 0.52 |
| | macro avg | 0.47 | 0.48 | 0.46 |
| | weighted avg | 0.57 | 0.52 | 0.53 |
| **Paclitaxel important features** | | | | |
| Granularity_1_OrigGray | | | | |
| Granularity_11_OrigGray | | | | |
| Granularity_4_OrigGray | | | | |
| Granularity_12_OrigGray | | | | |
| Granularity_3_OrigGray | | | | |
| | 0 | 0.80 | 0.89 | 0.84 |
| | 24 | 1.00 | 0.60 | 0.75 |
| | 48 | 0.50 | 0.50 | 0.50 |
| | 72 | 0.40 | 0.50 | 0.44 |
| | accuracy | 0.70 | 0.70 | 0.70 |
| | macro avg | 0.68 | 0.62 | 0.63 |
| | weighted avg | 0.74 | 0.70 | 0.71 |
| **Topotecan important features** | | | | |
| Granularity_2_OrigGray | | | | |
| Granularity_1_OrigGray | | | | |
| Granularity_11_OrigGray | | | | |
| Granularity_10_OrigGray | | | | |
| Granularity_5_OrigGray | | | | |
| **Topotecan report** | 0 | 0.67 | 0.75 | 0.71 |
| | 24 | 0.67 | 0.80 | 0.73 |
| | 48 | 0.83 | 0.83 | 0.83 |
| | 72 | 0.00 | 0.00 | 0.00 |
| | accuracy | 0.71 | 0.71 | 0.71 |
| | macro avg | 0.54 | 0.60 | 0.57 |
| | weighted avg | 0.65 | 0.71 | 0.68 |
| **Docetaxel important features** | | | | |
| Granularity_2_OrigGray | | | | |
| Granularity_1_OrigGray | | | | |
| Granularity_3_OrigGray | | | | |
| Granularity_4_OrigGray | | | | |
| Granularity_13_OrigGray | | | | |
| **Docetaxel report** | 0 | 0.60 | 0.38 | 0.46 |
| | 24 | 0.60 | 0.50 | 0.55 |
| | 48 | 0.50 | 0.75 | 0.60 |
| | 72 | 0.50 | 0.75 | 0.60 |
| | accuracy | 0.55 | 0.55 | 0.55 |

| | | | | |
|---|---|---|---|---|
| | macro avg | 0.55 | 0.59 | 0.55 |
| | weighted avg | 0.56 | 0.55 | 0.53 |
| **DMSO_30pc important features** | | | | |
| Granularity_2_OrigGray | | | | |
| Granularity_1_OrigGray | | | | |
| Granularity_4_OrigGray | | | | |
| Granularity_8_OrigGray | | | | |
| Granularity_10_OrigGray | | | | |
| **DMSO_30pc report** | 0 | 0.83 | 0.63 | 0.71 |
| | 24 | 1.00 | 0.67 | 0.80 |
| | 48 | 0.50 | 0.50 | 0.50 |
| | 72 | 0.50 | 1.00 | 0.67 |
| | accuracy | 0.68 | 0.68 | 0.68 |
| | macro avg | 0.71 | 0.70 | 0.67 |
| | weighted avg | 0.76 | 0.68 | 0.69 |
| **Campthotecin important features** | | | | |
| Granularity_2_OrigGray | | | | |
| Granularity_1_OrigGray | | | | |
| Granularity_16_OrigGray | | | | |
| Granularity_10_OrigGray | | | | |
| Granularity_7_OrigGray | | | | |
| **Campthotecin report** | 0 | 0.83 | 0.63 | 0.71 |
| | 24 | 0.86 | 0.86 | 0.86 |
| | 48 | 0.50 | 0.75 | 0.60 |
| | 72 | 0.75 | 0.75 | 0.75 |
| | accuracy | 0.74 | 0.74 | 0.74 |
| | macro avg | 0.74 | 0.75 | 0.73 |
| | weighted avg | 0.77 | 0.74 | 0.74 |
| **TEXTURE** | | | | |
| **Substance/features** | **Class(Time)** | **precision** | **recall** | **f1-score** |
| **Control important features** | | | | |
| Texture_Correlation_OrigGray_10_01_256 | | | | |
| Texture_SumVariance_OrigGray_10_01_256 | | | | |
| Texture_SumVariance_OrigGray_10_00_256 | | | | |
| Texture_SumVariance_OrigGray_5_03_256 | | | | |
| Texture_SumVariance_OrigGray_4_00_256 | | | | |
| **Control report** | 0 | 0.89 | 1.00 | 0.94 |
| | 24 | 1.00 | 1.00 | 1.00 |
| | 48 | 1.00 | 0.67 | 0.80 |
| | 72 | 0.67 | 1.00 | 0.80 |
| | accuracy | 0.91 | 0.91 | 0.91 |
| | macro avg | 0.89 | 0.92 | 0.89 |
| | weighted avg | 0.93 | 0.91 | 0.91 |
| **Doxorubicin important features** | | | | |
| Texture_Correlation_OrigGray_5_01_256 | | | | |
| Texture_Correlation_OrigGray_10_01_256 | | | | |

| | | | | |
|---|---|---|---|---|
| Texture_Correlation_OrigGray_3_01_256 | | | | |
| Texture_Correlation_OrigGray_10_03_256 | | | | |
| Texture_Variance_OrigGray_10_03_256 | | | | |
| **Doxorubicin report** | 0 | 1.00 | 1.00 | 1.00 |
| | 24 | 1.00 | 1.00 | 1.00 |
| | 48 | 0.67 | 1.00 | 0.80 |
| | 72 | 1.00 | 0.80 | 0.89 |
| | accuracy | 0.96 | 0.96 | 0.96 |
| | macro avg | 0.92 | 0.95 | 0.92 |
| | weighted avg | 0.97 | 0.96 | 0.96 |
| **Everolimus important features** | | | | |
| Texture_Correlation_OrigGray_5_01_256 | | | | |
| Texture_Correlation_OrigGray_3_01_256 | | | | |
| Texture_Correlation_OrigGray_4_02_256 | | | | |
| Texture_Correlation_OrigGray_5_02_256 | | | | |
| Texture_Correlation_OrigGray_4_01_256 | | | | |
| **Everolimus report** | 0 | 1.00 | 1.00 | 1.00 |
| | 24 | 0.50 | 0.75 | 0.60 |
| | 48 | 0.75 | 0.60 | 0.67 |
| | 72 | 0.50 | 0.40 | 0.44 |
| | accuracy | 0.71 | 0.71 | 0.71 |
| | macro avg | 0.69 | 0.69 | 0.68 |
| | weighted avg | 0.73 | 0.71 | 0.71 |
| **Oxalyplatin important features** | | | | |
| Texture_InverseDifferenceMoment_OrigGray_4_02_256 | | | | |
| Texture_InverseDifferenceMoment_OrigGray_5_02_256 | | | | |
| Texture_InverseDifferenceMoment_OrigGray_5_01_256 | | | | |
| Texture_InverseDifferenceMoment_OrigGray_10_00_256 | | | | |
| Texture_Correlation_OrigGray_4_02_256 | | | | |
| **Oxalyplatin report** | 0 | 1.00 | 1.00 | 1.00 |
| | 24 | 0.83 | 1.00 | 0.91 |
| | 48 | 1.00 | 1.00 | 1.00 |
| | 72 | 1.00 | 0.67 | 0.80 |
| | accuracy | 0.95 | 0.95 | 0.95 |
| | macro avg | 0.96 | 0.92 | 0.93 |
| | weighted avg | 0.96 | 0.95 | 0.95 |
| **Paclitaxel important features** | | | | |
| Texture_Correlation_OrigGray_4_02_256 | | | | |
| Texture_Correlation_OrigGray_3_01_256 | | | | |
| Texture_InfoMeas2_OrigGray_3_02_256 | | | | |
| Texture_SumAverage_OrigGray_5_00_256 | | | | |
| Texture_Correlation_OrigGray_5_02_256 | | | | |
| | 0 | 1.00 | 1.00 | 1.00 |
| | 24 | 0.80 | 0.80 | 0.80 |
| | 48 | 0.20 | 0.50 | 0.29 |
| | 72 | 1.00 | 0.25 | 0.40 |

| | | | | |
|---|---|---|---|---|
| | accuracy | 0.75 | 0.75 | 0.75 |
| | macro avg | 0.75 | 0.64 | 0.62 |
| | weighted avg | 0.87 | 0.75 | 0.76 |
| **Topotecan important features** | | | | |
| Texture_SumEntropy_OrigGray_4_03_256 | | | | |
| Texture_SumVariance_OrigGray_4_00_256 | | | | |
| Texture_SumEntropy_OrigGray_3_00_256 | | | | |
| Texture_SumVariance_OrigGray_4_01_256 | | | | |
| Texture_InverseDifferenceMoment_OrigGray_4_02_256 | | | | |
| **Topotecan report** | 0 | 1.00 | 1.00 | 1.00 |
| | 24 | 1.00 | 1.00 | 1.00 |
| | 48 | 1.00 | 1.00 | 1.00 |
| | 72 | 1.00 | 1.00 | 1.00 |
| | accuracy | 1.00 | 1.00 | 1.00 |
| | macro avg | 1.00 | 1.00 | 1.00 |
| | weighted avg | 1.00 | 1.00 | 1.00 |
| **Docetaxel important features** | | | | |
| Texture_Correlation_OrigGray_5_02_256 | | | | |
| Texture_Correlation_OrigGray_3_00_256 | | | | |
| Texture_Correlation_OrigGray_4_02_256 | | | | |
| Texture_Correlation_OrigGray_5_03_256 | | | | |
| Texture_Correlation_OrigGray_3_02_256 | | | | |
| **Docetaxel report** | 0 | 1.00 | 1.00 | 1.00 |
| | 24 | 1.00 | 1.00 | 1.00 |
| | 48 | 1.00 | 1.00 | 1.00 |
| | 72 | 1.00 | 1.00 | 1.00 |
| | accuracy | 1.00 | 1.00 | 1.00 |
| | macro avg | 1.00 | 1.00 | 1.00 |
| | weighted avg | 1.00 | 1.00 | 1.00 |
| **DMSO_30pc important features** | | | | |
| Texture_Correlation_OrigGray_3_01_256 | | | | |
| Texture_Correlation_OrigGray_5_01_256 | | | | |
| Texture_Correlation_OrigGray_10_01_256 | | | | |
| Texture_Variance_OrigGray_4_01_256 | | | | |
| Texture_Contrast_OrigGray_3_03_256 | | | | |
| **DMSO_30pc report** | 0 | 0.89 | 1.00 | 0.94 |
| | 24 | 0.75 | 1.00 | 0.86 |
| | 48 | 1.00 | 0.75 | 0.86 |
| | 72 | 1.00 | 0.50 | 0.67 |
| | accuracy | 0.86 | 0.86 | 0.86 |
| | macro avg | 0.91 | 0.81 | 0.83 |
| | weighted avg | 0.89 | 0.86 | 0.85 |
| **Campthotecin important features** | | | | |
| Texture_SumVariance_OrigGray_3_02_256 | | | | |
| Texture_InfoMeas2_OrigGray_10_01_256 | | | | |
| Texture_SumVariance_OrigGray_3_00_256 | | | | |

| | | | | |
|---|---|---|---|---|
| Texture_Variance_OrigGray_10_01_256 | | | | |
| Texture_Correlation_OrigGray_3_02_256 | | | | |
| **Campthotecin report** | 0 | 1.00 | 1.00 | 1.00 |
| | 24 | 1.00 | 1.00 | 1.00 |
| | 48 | 1.00 | 0.50 | 0.67 |
| | 72 | 0.67 | 1.00 | 0.80 |
| | accuracy | 0.91 | 0.91 | 0.91 |
| | macro avg | 0.92 | 0.88 | 0.87 |
| | weighted avg | 0.94 | 0.91 | 0.91 |
